# Beyond accessibility: ATAC-seq footprinting unravels kinetics of transcription factor binding during zygotic genome activation

**DOI:** 10.1101/869560

**Authors:** Mette Bentsen, Philipp Goymann, Hendrik Schultheis, Kathrin Klee, Anastasiia Petrova, René Wiegandt, Annika Fust, Jens Preussner, Carsten Kuenne, Thomas Braun, Johnny Kim, Mario Looso

## Abstract

While footprinting analysis of ATAC-seq data can theoretically enable investigation of transcription factor (TF) binding, the lack of a computational tool able to conduct different levels of footprinting analysis has so-far hindered the widespread application of this method. Here we present TOBIAS, a comprehensive, accurate, and fast footprinting framework enabling genome-wide investigation of TF binding dynamics for hundreds of TFs simultaneously. As a proof-of-concept, we illustrate how TOBIAS can unveil complex TF dynamics during zygotic genome activation (ZGA) in both humans and mice, and explore how zygotic Dux activates cascades of TFs, binds to repeat elements and induces expression of novel genetic elements. TOBIAS is freely available at: https://github.com/loosolab/TOBIAS.

## Background

Epigenetic mechanisms governing chromatin organization and transcription factor (TF) binding are critical components of transcriptional regulation and cellular transitions. In recent years, rapid improvements of pioneering sequencing methods such as ATAC-seq (Assay of Transposase Accessible Chromatin) ^1^, have allowed for systematic, global scale investigation of epigenetic mechanisms controlling gene expression. While ATAC-seq can uncover accessible regions where TFs might bind, reliable identification of specific TF binding sites (TFBS) still relies on chromatin immunoprecipitation methods such as ChIP-seq. However, ChIP-seq methods require high input cell numbers, are limited to one TF per assay, and are further restricted to TFs for which antibodies are readily available. Latest improvements of ChIP based methods ^2^ can circumvent some of these technical drawbacks, but the limitation of only being able to identify binding sites of one TF per assay persists. Therefore, it remains costly, or even impossible, to study the binding of multiple TFs in parallel.

Current limits to the investigation of TF binding become particularly apparent when investigating processes involving a very limited number of cells such as preimplantation development (PD) of early zygotes. During PD, the fertilized egg forms the zygote, which undergoes a series of cell divisions to finally constitute the blastocyst, a structure built by the inner cell mass (ICM) and trophectoderm (Figure 1a). Within this process, maternal and paternal mRNAs are degraded prior to zygotic genome activation (ZGA) (reviewed in ^3^), a transformation which eventually leads to the transcription of thousands of genes ^4^. Integration of multiple omics-based profiling methods have revealed a set of key TFs that are expressed at the onset of and during ZGA including Dux ^5, 6^, Zscan4 ^7^, and other homeobox-containing TFs ^8^. However, due to the limitations of ChIP-seq, the exact genetic elements bound and regulated by different TFs during PD remain to be fully discovered. Consequently, the global network of TF binding dynamics throughout PD remains mostly obscure.

**Figure 1:**
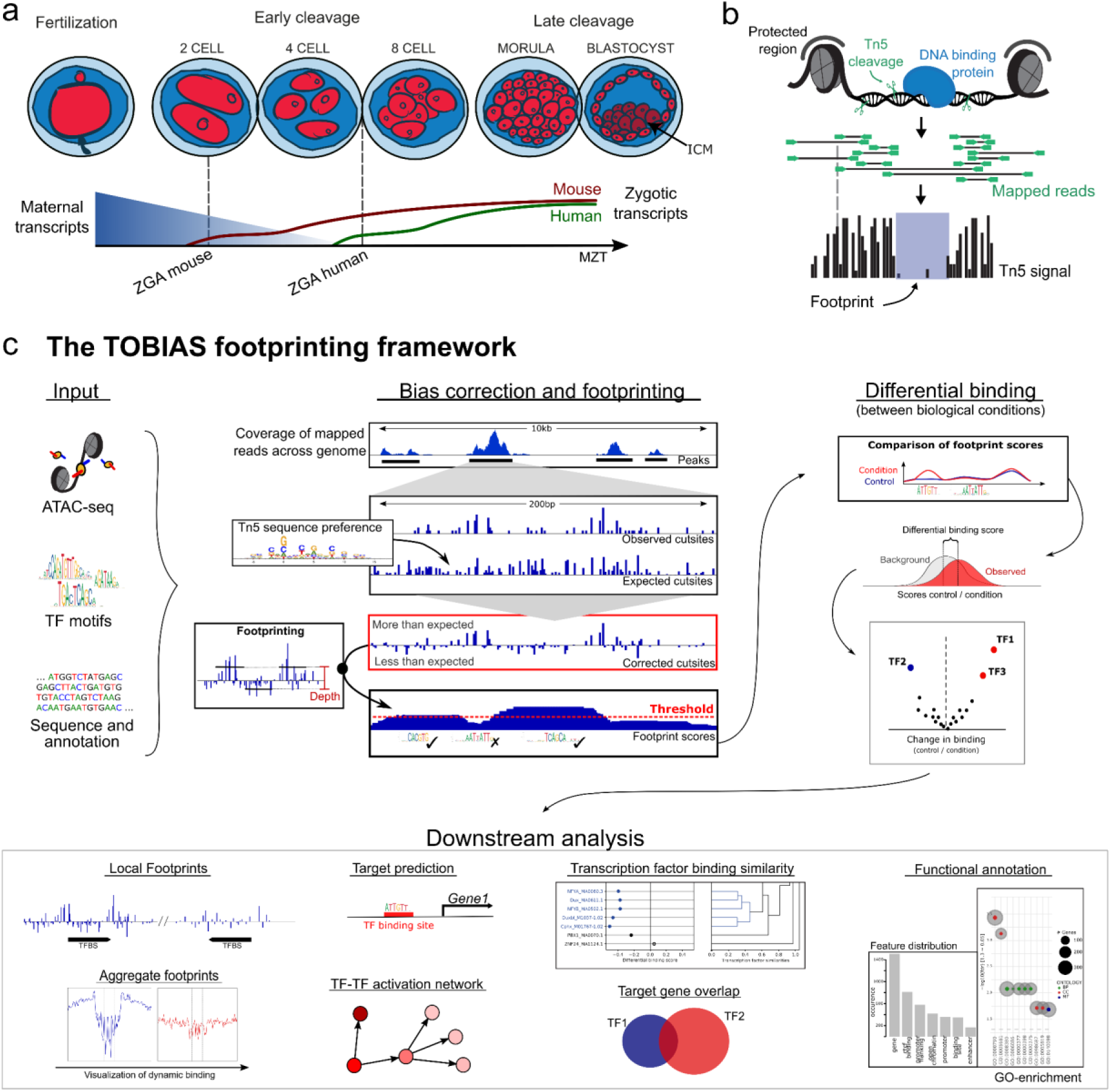
The use of chromatin accessibility assays to investigate early developmental processes. **(a) Early embryonic development in human and mouse.** The fertilized egg undergoes a series of divisions ultimately creating the structure of the blastocyst. While maternal transcripts are depleted, the zygotic genome is activated in waves. ZGA initiates in mouse at 2-cell stage and in human at the 4-8-cell stage. **(b) The concept of footprinting using ATAC-seq**. The Tn5 transposase cleaves and inserts sequencing adapters in open chromatin, but is unable to cut in chromatin occupied by e.g. nucleosomes or transcription factors. The mapped sequencing reads are used to create a signal of single Tn5 insertion events, in which binding of transcription factors is visible as depletion of signal (the footprint). **(c) The TOBIAS digital genomic footprinting framework**. Using an input of sequencing reads from ATAC-seq, transcription factor motifs and sequence information, the TOBIAS footprinting framework detects local and global changes in transcription factor binding. Bias-correction of the Tn5 sequence preference enables detection of local chromatin footprints and matching to individual TFBS. Footprint scores are compared between conditions to define differentially bound TFs. The global binding map allows for a variety of downstream analysis such as visualization of local and aggregated footprints across conditions, prediction of target genes for each TF as well as comparison of binding specificity between several transcription factors. Functional annotation such as GO enrichment can be used to infer biological meaning of target gene sets.

A computational method known as *digital genomic footprinting* (DGF) ^9^ has emerged as an alternative means, which can overcome some of the limits of ChIP-based methods. DGF is a computational analysis of chromatin accessibility assays such as ATAC-seq, which employs DNA effector enzymes that only cut accessible DNA regions. Similarly to nucleosomes, bound TFs hinder cleavage of DNA, resulting in defined regions of decreased signal strength within larger regions of high signal - known as *footprints* ^10^ (Figure 1b).

Surprisingly, although this concept shows considerable potential to survey genome-wide binding of multiple TFs in parallel from a single experiment, DGF analysis is rarely applied when investigating TF binding mechanisms. The skepticism towards DGF has been driven by the discovery that enzymes used in chromatin accessibility assays (e.g. DNase-I) are biased towards certain sequence compositions, an effect which has been well characterized for DNase-seq ^11, 12^. The influence of Tn5 transposase bias in the context of ATAC-seq footprinting has, however, only been described very recently ^13, 14^ and still represents an uncertainty during discovery of true footprints. Besides the identification of footprints, comparing footprints across biological conditions remains challenging as well. While there have been efforts to estimate differential TF binding on a genomewide scale ^15, 16^, investigation of epigenetic processes often require more in-depth information on the individual differentially bound TFBS and genes targeted by these TFs, which is not provided by these methods. Furthermore, many footprinting methods suffer from performance issues due to missing support for multiprocessing, inflexible software architecture prone to software dependency issues, and the use of non-standard file-formats. These obstacles complicate the assembly of different tools for advanced analysis workflows. Consequently, despite its compelling potential, these issues have rendered footprinting on ATAC-seq cumbersome to apply to biological questions. Essentially, a comprehensive framework enabling large-scale ATAC-seq footprinting is missing.

Here, we describe and apply TOBIAS (**T**ranscription factor **O**ccupancy prediction **B**y **I**nvestigation of **A**TAC-seq **S**ignal), a comprehensive computational framework that we created for footprinting analysis (Figure 1c). TOBIAS is a collection of command-line tools, which provides functionality to perform all levels of footprinting analysis including bias correction, footprinting, and comparison between conditions (Supp. Figure 1; Footprinting pipeline). Furthermore, TOBIAS includes a variety of auxiliary tools such as TF network inference and visualization of footprints, which can be combined for more targeted downstream analysis (Supp. Figure 1; Supporting tools). To couple individual tools, we provide scalable analysis workflows implemented in Snakemake 17 and NextFlow 18, including a cloud computing compatible version making use of the de.NBI cloud ^19^. These pipelines utilize a minimal input of ATAC-seq reads, TF motifs and genome information to enable complete footprinting analysis and comparison of TF binding even for complex experimental designs (e.g. time series).

## Results

### Classification and validation of TOBIAS

A computational DGF framework able to perform footprinting on ATAC-seq data and handle complex experimental designs autonomously does not exist. Nonetheless, to demonstrate the advantages of TOBIAS, we compared the individual framework features to published footprinting tools for ATAC-seq footprinting where applicable. While we found that some functionalities are overlapping between tools, we found a substantial set of features exclusively covered by TOBIAS (Table 1). As sequencing costs will continue to decrease, allowing for ever more data to be created, it is worth noting that TOBIAS is the only tool supporting differential footprinting for more than two conditions. Additionally, TOBIAS is one of only two footprinting tools applying multiprocessing to speed up computation, resulting in the lowest runtime among the compared set of tools.

**Table 1:** Comparison of features for ATAC-seq footprinting tools.

|  | Footprinting tools for ATAC-seq |  |  |  |  |
| --- | --- | --- | --- | --- | --- |
|  | TOBIAS | HINT-ATAC | MsCentipede | PIQ | Wellington |
| **Overview:** |  |  |  |  |  |
| Year of publication | - | 2019 | 2015 | 2014 | 2013 |
| Tool availability | Github | Github | Github | Bitbucket | Github |
| Programming language | Python | Python | Python | R | Python |
| Type of footprinting<br>(D: De novo, M: Motif-centric) | D | D | M | M | D |
| **Features:** |  |  |  |  |  |
| Footprinting | ✓ | ✓ | ✓ | ✓ | ✓ |
| Tn5 bias-correction | ✓ | ✓ | ✗ | ✗ | ✗ |
| Size-adjustable footprinting algorithm | ✓ | ✗ | ✗ | ✗ | ✗ |
| Differential footprinting | ✓ | ✓ | ✗ | ✗ | ✓ |
| Time series footprinting<br>(comparison of 2+ conditions) | ✓ | ✗ | ✗ | ✗ | ✗ |
| Calculation of TFBS (from motifs) | ✓ | ✓ | ✗ | ✓ | ✗ |
| TFBS clustering | ✓ | ✗ | ✗ | ✗ | ✗ |
| Consensus motifs for clustered motifs | ✓ | ✗ | ✗ | ✗ | ✗ |
| Output of genomic tracks | ✓ | ✓ | ✗ | ✗ | ✓ |
| Adjustable plotting of aggregate footprints | ✓ | ✗ | ✗ | ✗ | ✗ |
| Visualization of locus footprints | ✓ | ✗ | ✗ | ✗ | ✗ |
| Inference of TF-binding networks | ✓ | ✗ | ✗ | ✗ | ✗ |
| Predict bound/unbound state per TFBS | ✓ | ✗ | ✓ | ✓ | ✗ |
| **Usability:** |  |  |  |  |  |
| Uses standard file formats | ✓ | ✓ | ✗ | ✗ | ✓ |
| Parallel computing | ✓ | ✗ | ✗ | ✗ | ✓ |
| Complete workflow available | ✓ | ✗ | ✗ | ✗ | ✗ |
| - Snakemake | ✓ | ✗ | ✗ | ✗ | ✗ |
| - Nextflow | ✓ | ✗ | ✗ | ✗ | ✗ |
| Cloud computing supported | ✓ | ✗ | ✗ | ✗ | ✗ |
| Time to execute* (min) | 7.2 | 46 | 2808 | 329 | 8 |
| Package manager/installer | PyPI<br>Bioconda | PyPI<br>Bioconda | - | - | PyPI<br>Bioconda |

\* CPU time (using 30 cores if applicable) to perform bias correction (if applicable) and footprinting for GM12878 chromosome 1 using 54 transcription factors matched to ENCODE ChIP-seq.

To compare the footprinting capabilities of individual tools, we utilized 218 paired ChIP-seq / ATAC-seq datasets across four different cell types. Here, the ChIP-seq data represents the true binding sites for each TF, which we used for validating each method after application to the matched ATAC-seq data (see Methods; Validation). In terms of Tn5 bias correction, as well as visualization of aggregate footprints, we found that TOBIAS clearly outperforms other bias-correction tools in uncovering footprints and thereby distinguishing between bound/unbound sites (Supp. Figure 2a, Supp. File 1). For the task of protein binding prediction, we found that TOBIAS significantly outperformed the other de novo tools HINT-ATAC, PIQ and Wellington (Supp. Figure 2b) and performed equally well as msCentipede overall (Supp. Figure 2c). Notably, TOBIAS also showed robust performance across individual cell types (Suppl. Figure 2d). Looking at individual TFs, TOBIAS outperforms msCentipede for factors with a notable gain of footprints after Tn5 bias correction (Supp. Figure 2e), once again highlighting the advantage of taking Tn5 bias into account. Although msCentipede implements a motif centric learning approach, which can take TF specific binding patterns into account, it did not yield overall higher accuracy in comparison to TOBIAS. Additionally, the approach of building individual TF models took 300 times longer to compute than performing footprinting using TOBIAS (Supp. Figure 2f and Table 1). Such learning approaches are therefore greatly limited in the number of TFs and conditions that can realistically be included in an analysis. In conclusion, we find that the TOBIAS framework shows unprecedented accuracy and performance in the field of ATAC-seq footprinting.

In order to confirm the improvement of footprint detection after Tn5 bias correction, we made use of another exemplary dataset derived from hESC ^20^. Importantly, besides cases where the footprint was hidden by Tn5 bias (Supp. Figure 3a; ZSCAN4), we also identified TFs for which the motif itself disfavors Tn5 integration, thereby creating a false-positive footprint in uncorrected signals, which disappears after Tn5 bias correction (Supp. Figure 3a; HLX). Utilizing a footprint depth metric as described by ^16^ (Supp. Figure 3b) across uncorrected, expected and corrected Tn5 signals, we found a high correlation between uncorrected and expected footprinting depths (Supp. Figure 3c). In contrast, this effect vanished after TOBIAS correction (Supp. Figure 3d), effectively uncovering TF footprints which were superimposed by Tn5 bias. From a global perspective, taking 590 TFs into account, TOBIAS generated a measurable footprint for 64% of the TFs (Supp. Figure 3e). This is in contrast to previous reports wherein it has been suggested that only 20% of all TFs leave measurable footprints ^16^. To summarize, we found that TOBIAS exceeded other tools in terms of uncovering footprints and correctly identifying bound TF binding sites.

### Footprinting uncovers transcription factor binding dynamics in mammalian ZGA

To demonstrate the full potential of TOBIAS, in particular in the investigation of processes involving only few cells, we analyzed a series of ATAC-seq datasets derived from both human and murine preimplantation embryos at different developmental stages ranging from 2C, 4C, 8C to ICM in addition to embryonic stem cells of their respective species ^20, 21^. Altogether, TOBIAS was used to calculate footprint scores for a list of 590 and 464 individual TFs across the entire process of PD of human and mouse embryos, respectively. After clustering TFs into co-active groups within one or multiple developmental timepoints, we first asked whether the predicted timing of TF activation reflects known processes in human PD. Intriguingly, we found 10 defined clusters of specific binding patterns, the majority of which peaked between 4C and 8C, fully concordant with the transcriptional burst and termination of ZGA (Figure 2a and Supp. Table 1).

**Figure 2:**
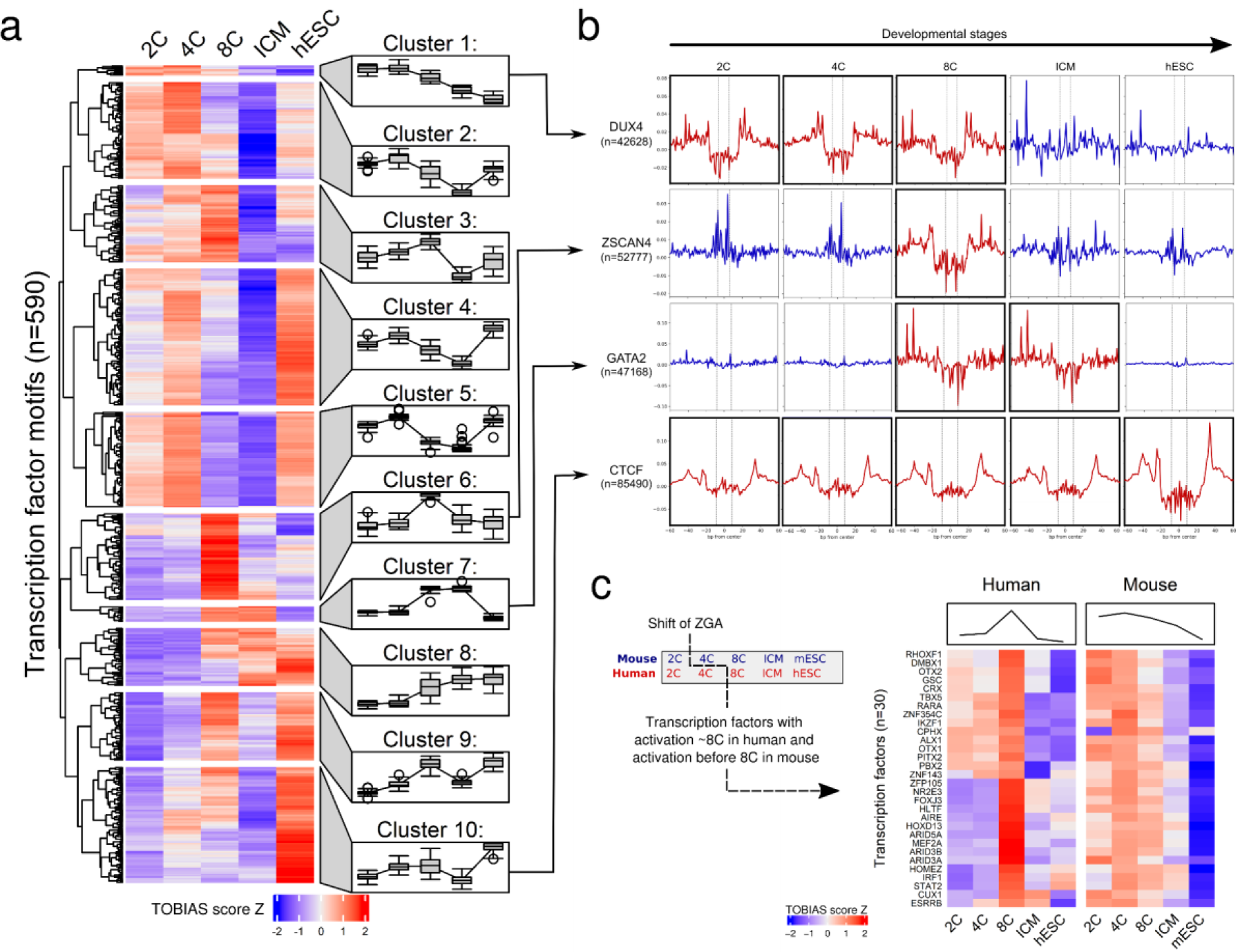
TOBIAS enables investigation of global changes in transcription factor binding. **(a) Clustering of transcription factor activities throughout development**. Each row represents one TF, each column a developmental stage; blue color indicates low activity, red color indicates high activity. In order to visualize cluster trends, each cluster is associated with a mean trend line and time point specific boxplots. **(b) Bias-corrected ATAC-seq footprints reveal dynamic TF binding**. Aggregated footprinting plot matrix for transcription factor binding sites. Plots are centered around binding motifs (n=* relates to the number of binding sites). Rows indicate TFs DUX4, ZSCAN4, GATA2, and CTCF; columns illustrate developmental stages from left to right. Active binding of the individual TFs is visible as depletion in the signal around the binding site (highlighted in red). Upper three TFs are related to developmental stages, CTCF acts as a universal control, generating a footprint in all conditions. See Supplementary Figure 3f for uncorrected footprints. **(c) TF activity is shifted by ZGA onset in human and mouse**. Heatmaps show activity of known ZGA-related TFs for human (left) and mouse (right) across matched timepoints 2C / 8C / ICM / hESC (mESC). Mean TF activity (top panel) peaks at 4-8C stage in human and is shifted to 2-4C stage in mouse by the earlier ZGA onset.

Two clusters of TFs (Cluster 1+2; n=83) displayed highest activity at the 2-4C stage and strongly decreased thereafter, suggesting that factors within these clusters are likely involved in ZGA initiation. We set out to classify these TFs, and observed a high overlap with known maternally transferred transcripts ^22^ (LHX8, BACH1, EBF1, LHX2, EMX1, MIXL1, HIC2, FIGLA, SALL4, ZNF449), explaining their activity before ZGA onset. Importantly, DUX4 and DUXA, which are amongst the earliest expressed genes during ZGA ^5, 6^, were also contained in these clusters. Additional TFs included HOXD1, which is known to be expressed in human unfertilized oocytes and preimplantation embryos ^23^ and ZBTB17, a TF mandatory to generate viable embryos ^24^. Cluster 6 (n=67) displayed a particularly prominent 8C specific signature, that harbored well known TFs involved in lineage specification such as PITX1, PITX3, SOX8, MEF2A, MEF2D, OTX2, PAX5 and NKX3.2. Furthermore, overlapping TFs within Cluster 6 with RNA expression datasets ranging from the germinal vesicle to cleavage stage ^5^, 12 additional TFs (FOXJ3, HNF1A, ARID5A, RARB, HOXD8, TBP, ZFP28, ARID3B, ZNF136, IRF6, ARGFX, MYC, ZSCAN4) were confirmed to be exclusively expressed within this time frame. Taken together, these data show that TOBIAS reliably uncovers massively parallel TF binding dynamics at specific time points during early embryonic development.

### Transcription factor scores correlate with footprints and gene expression

To confirm that TOBIAS-based footprinting scores are indeed associated with leaving *bona fide* footprints we utilized the ability to visualize aggregated footprint plots as implemented within the framework. Indeed, bias corrected footprint scores were highly congruent with explicitly defined footprints (Figure 2b) of prime ZGA regulators at developmental stages in which these have been shown to be active ^7^. For example, footprints associated with DUX4, a master inducer of ZGA, were clearly visible from 2C-4C, decreased from 8C onwards and were completely lost in later stages, consistent with known expression levels ^20^ and ZGA onset in humans. Footprints for ZSCAN4, a primary DUX4 target ^5^, were exclusively visible at the 8C stage. Interestingly, GATA2 footprints were exhibited from 8C to ICM stages which is in line with its known function in regulating trophoblast differentiation ^25^. As expected, CTCF creates footprints across all timepoints. Strikingly, we observed that these defined footprints were not detectable without TOBIAS mediated Tn5 bias correction (Supp. Figure 3f). These data show that footprint scores can be reliably confirmed by footprint visualizations, which further allow to infer TF binding dynamics.

To test if the global footprinting scores of individual TFs correlate with the incidence and level of their RNA expression, we matched them to RNA expression datasets derived from individual timepoints throughout zygotic development, taking TF motif similarity into account. Indeed, we found that TOBIAS scores for the majority of TFs either correlated well with the timing of their expression profiles or displayed a slightly delayed activity after expression peaked (Supp. Figure 4a). This is important because it shows that in conjunction with expression data, TOBIAS can unravel the kinetics between TF expression (mRNA) and the actual binding activity of their translated proteins. The value of this added information becomes particularly apparent when analyzing activities of TFs that did not correlate with the timing of their RNA expression (Supp. Figure 4a; not correlated).

For example, within the non-correlated cluster 13 TFs were identified which are of putative maternal origin ^22^ including SALL4. In mice, Sall4 protein is maternally contributed to the zygote, subsequently degraded at 2C and then reexpressed after zygotic transcription has initiated ^26^. Consistent with this, SALL4 expression increases dramatically from 8C onwards (Supp. Table 2). In contrast to the expression values, TOBIAS predicted SALL4 to have the highest activity in 2C and second-highest activity in hESC (on-off-on-pattern), which is in line with the presence of maternal SALL4 in the zygote. These data show that TOBIAS can predict true on-off-on-patterns, and can infer significant insight into TF activities, in particular for those where determining their expression patterns alone does not suffice to explain when they exert their biological function.

### Differential footprint analysis reveals functional divergence between human and mouse ZGA

The timing of ZGA varies between mice (2C) and humans (4C to 8C) (reviewed in ^27 28^). By integrating the TOBIAS scores from human and mouse (Supp. Figure 4b and Supp. Table 3), and instrumentalizing the capability of TOBIAS to generate differential TF binding plots for all time points automatically, we investigated similarities and differences of PD between these species. Firstly, reflecting the shift of ZGA onset, we identified 30 TFs which appeared to be ZGA specific in both human and mouse (Figure 2c) including several homeobox factors which already have described functions within ZGA ^29^ as well as ARID3A which has been shown to play a role in cell fate decisions in creating trophectoderm ^30^.

Next, we used the differential TF binding plots to display differences in ZGA at the transition between 2C and 4C in mouse (Figure 3a), and human 8C and ICM (Figure 3b) (Supp. File 2 + 3 for all pairwise comparisons). In mice, we observed a shift of Obox-factor activity in 2C to an activation of Tead (Tead1-4) and AP-2 (Tfap2a/c/e) motifs in 4C. Notably, AP-2/Tfap2c is required for normal embryogenesis in mice ^31^ and was also recently shown to act as a chromatin modifier that opens enhancers proximal to pluripotency factors in human ^32^. We observed a similar shift of TF activity for homeobox factors such as PITX1-3, RHOXF1, CRX and DMBX1 at the human 8C stage towards higher scores in ICM for known pluripotency factors such as POU5F1 (OCT4) and other POU-factors. Taken together, these results highlight the ability of TOBIAS to capture differentially bound TFs, not only across the whole timeline, but also between individual conditions and species.

**Figure 3:**
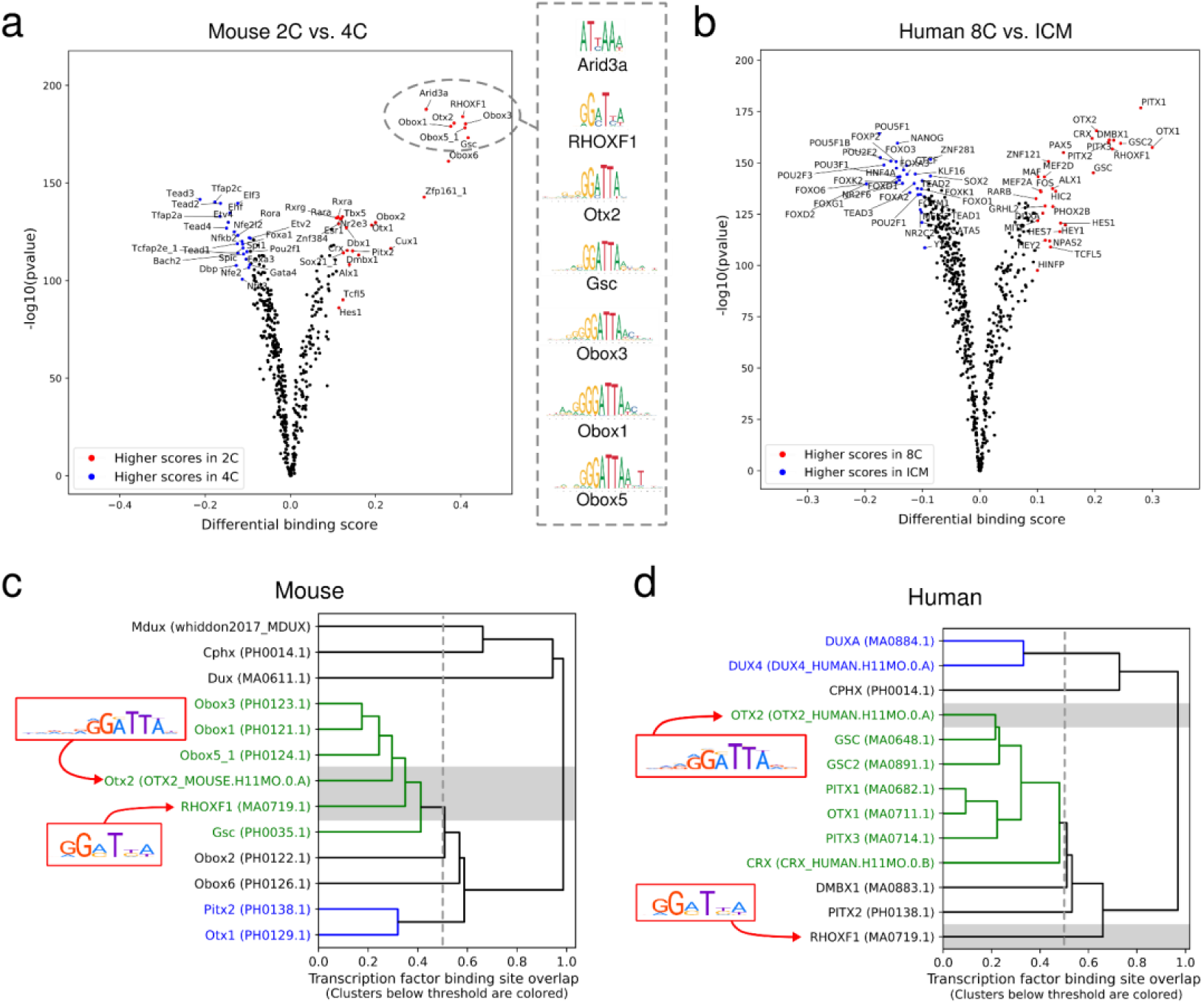
Comparison of binding site overlaps shows specification of ZGA functions between mouse and human. **(a-b) Pairwise comparison of TF activity between developmental stages**. The volcano plots show the differential binding activity against the -log10(pvalue) (as provided by TOBIAS) of the investigated TF motifs; each dot represents one motif. For (A) 2C stage specific/significant TFs are labeled in red, 4C specific factors are given in blue. For (B) 8C stage specific/significant TFs are labeled in red, ICM specific factors are given in blue. **(c-d) Clustering of TF motifs based on binding site overlap**. Excerpt of the global TF clustering based on TF binding location, illustrating individual TFs as rows. The trees indicate genomic positional overlap of individual TFBS with a tree-depth of 0.2 representing an overlap of 80% of motifs. Each TF is indicated by name and unique ID in brackets. Clusters of TFs with more than 50% overlap (below 0.5 tree distance) are colored. (C) shows overlap of motifs included in the mouse analysis, and (D) shows clustering of human motifs. Complete TF trees are provided in Supp. Files 2 and 3.

Throughout the pairwise comparisons, we observed that TFs from the same families often display similar binding kinetics within species, which is not surprising since they often possess highly similar binding motifs (Figure 3a right). To characterize TF similarity, TOBIAS clusters TFs based on the overlap of TFBS within investigated samples (Figure 3c+3d). This enables quantification of the similarity and clustering of individual TFs that appear to be active at the same time. Thereby, we observed a group of homeobox motifs which cluster together with more than 50% overlap of their respective binding sites in mouse (Figure 3c). In contrast, other TFs such as Tead and AP-2 cluster separately, indicating that these factors utilize independent motifs (Supp. File 2+3). While this might appear trivial, this clustering of TFs in fact also highlights differences in motif usage between human and mouse. One prominent example is the RHOXF1 motif, which shows high binding-site overlap with Obox 1/3/5 and Otx2 binding sites in mouse (Figure 3c; ∼60% overlap), but does not cluster with OTX2 in human (Figure 3d; ∼35% overlap). This observation suggests important functional differences of RHOX/Rhox TFs between mice and humans. In support of this hypothesis RHOXF1, RHOXF2 and RHOXF2B genes are exclusively expressed at 8C and ICM in humans, whereas Rhox factors are not expressed in corresponding developmental stages of preimplantation in mouse (Supp. Table 4). Conceivably, this observation, together with the finding that murine Obox factors share the same motif as RHOX-factors in humans, suggests that Obox TFs might function similarly to RHOX-factors during ZGA. Altogether, the TOBIAS mediated TF clustering based on TFBS overlap allows for quantification of target-similarity and divergence of TF function between motif families.

### Dux expression induces massive changes of chromatin accessibility, transcription and TF networks

Throughout the investigations of human and mouse development we became particularly interested in the Dux/DUX4 TF, which TOBIAS predicted to be one of the earliest factors to be active in both organisms (Figure 2a, Supp. Figure 4b and Supp. Table 1+3). Interestingly, despite the fact that Dux has already been proved to play a prominent role in ZGA ^5-7, 33, 34^, there is still a poor understanding of how Dux regulates its primary downstream targets, and consequently its secondary targets, during this process. We therefore applied TOBIAS to identify Dux binding sites utilizing an ATAC-seq dataset of Dux overexpression (DuxOE) in mESC ^5^.

As expected, the differential TF activity predicted by TOBIAS showed an increase in activity of Dux, Obox and other homeobox-TFs (Figure 4a, Supp. File 4). Interestingly, this was accompanied by a massive loss of TF binding for pluripotency markers such as Nanog, Pou5f1 (OCT4) and Sox2 upon DuxOE, indicating that Dux renders previously accessible chromatin sites associated with pluripotency inaccessible.

**Figure 4:**
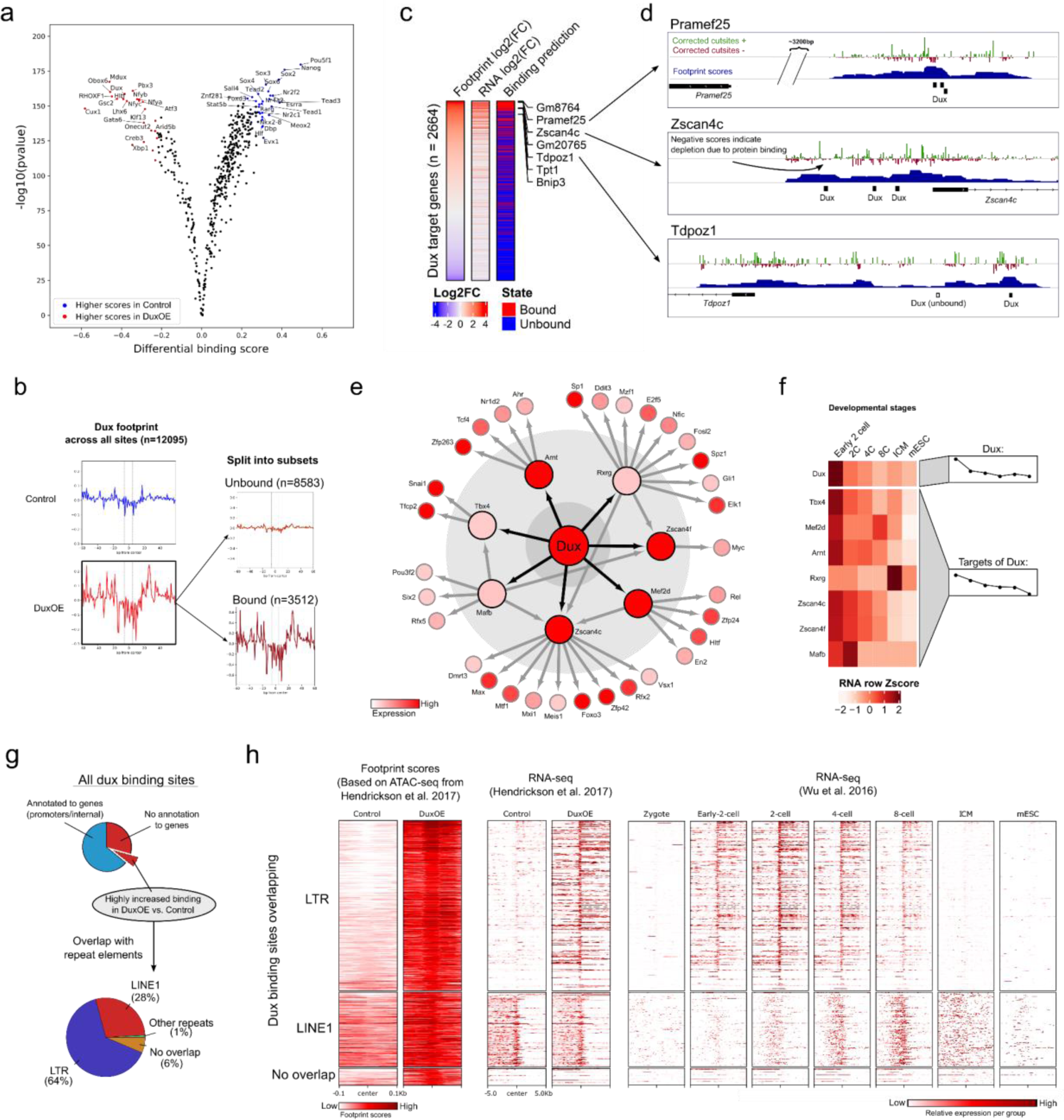
Dux binding induces transcription at gene promoters and LTR sequences in mouse. **(a) Volcano plot comparing TF activities between mDux GFP-(Control) and mDux GFP+ (DuxOE)**. Volcano plot showing the TOBIAS differential binding score on the x-axis and -log10 (p value) on the y-axis; each dot represents one TF. DuxOE specific TFs are labeled in red and Control specific TFs are labeled in blue. **(b) Aggregated footprint plots for Dux**. The aggregate plots are centered on the predicted binding sites for Dux between Control and DuxOE condition. The total possible binding sites for DuxOE (n=12095) are separated into bound and unbound sites (right). The dashed line represents the edges of the Dux motif. **(c) Change in expression of genes near Dux binding sites**. The heatmap shows 2664 Dux binding sites found in gene promoters. Footprint log2(FC) and RNA log2(FC) represent the changes between Control and DuxOE for footprints and gene expression, respectively. Log2(FC) is calculated as log2(DuxOE/Control). The column “Binding prediction” depicts whether the binding site was predicted by TOBIAS to be bound/unbound in the DuxOE condition. **(d) Genomic tracks showing footprint scores of Dux-binding**. Genomic tracks indicating three DUX target gene promoters (one per row) and respective tracks for cut site signals (red/blue), TOBIAS footprints (blue), detected motifs (black boxes), and gene locations (solid black boxes with arrows indicating gene strand). **(e) Dux transcription factor network**. The TF-TF network is built of all TFBS with binding in TF promoters with increasing strength in DuxOE (log2(FC)>0). Sizes of nodes represent the level of the network starting with Dux (Large: Dux, Medium: 1st level, Small: 2nd level). Nodes are colored based on RNA level in the OE condition. **(f) Correlation of the Dux transcription factor network to expression during development**. The heatmap depicts the in vivo gene expression during developmental stages from ^21^. The right-hand group annotation highlights the difference in mean expression for each timepoint. The heatmap is split into Dux and target genes of Dux. **(g) Dux binding sites overlap with repeat elements**. All potential Dux binding sites are split into sites either overlapping promoters/genes or without annotation to any known genes. The bottom pie chart shows a subset of the latter, additionally having highly increased binding (log2(FC)>1), and overlapping LTR/LINE1 elements. **(h) Dux induces expression of transcripts specific for preimplantation**. Genomic signals for the Dux binding sites which are bound in DuxOE with log2(FC) footprint score >1 (i.e. upregulated in DuxOE) are split into overlapping either LTR, LINE1 or no known genetic elements (top to bottom). Footprint scores (+/- 100bp from Dux binding sites) indicate the differential Dux binding between control and DuxOE. RNA-seq shows the normalized read-counts from ^5^ and ^21^ within +/- 5kb of the respective Dux binding sites where red color indicates high expression.

Consistently, Dux footprints (Figure 4b; left) were clearly evident upon DuxOE. In comparison to existing bias-correction methods, we found TOBIAS to be superior in uncovering this footprint between Control and DuxOE conditions (Supp. Figure 5a). Importantly, TOBIAS additionally discriminated ∼30% of all potential binding sites within open chromatin regions to be bound in the DuxOE condition, which further demonstrates the specificity of this method (Figure 4b; right). To rank the biological relevance of the individually changed binding sites between control and DuxOE conditions, we linked all annotated gene loci to RNA expression. A striking correlation between the gain-of-footprint and gain-of-expression of corresponding loci was clearly observed and mirrored by the TOBIAS predicted bound/unbound state (Figure 4c). Amongst the genes within the list of bound Dux binding sites (Supp. Table 5 for full Dux target list) were well known Dux targets including *Zscan4c* and *Pramef25* ^35^, for which local footprints for Dux were clearly visible (Figure 4d). The high resolution of footprints is particularly pronounced for *Tdpoz1* which harbors two potential Dux binding sites of which one is clearly footprinted in the score track, while the other is predicted to be unoccupied (Figure 4d; bottom). In line with this, *Tdpoz1* expression is significantly upregulated upon DuxOE as revealed by RNA-seq (log2FC: 6,95). Consistently, *Tdpoz1* expression levels are highest at 2C in zygotes and decrease thereafter, strongly indicating that *Tdpoz1* is likely a direct target of Dux during PD both *in vitro* and *in vivo* ^21, 36^ (Supp. Table 5). Footprinting scores also directly correlated with ChIP-seq peaks for Dux in the Tdpoz1 promoter (Supp. Figure 5b), an observation which we also found at many other positions (Examples shown in Supp. Figure 5c+d).

Many of the TOBIAS-predicted Dux targets encode TFs themselves. Therefore, we applied the TOBIAS network module to subset and match all activated binding sites to TF target genes with the aim of inferring how these TF activities might connect. Thereby, we could model an intriguing pseudo timed TF activation network. This directed network uncovered a TF activation cascade initiated by Dux, resulting in the activation of 7 primary TFs which appear to subsequently activate 32 further TFs (first three layers depicted in Figure 4e). As Dux is a regulator of ZGA, we asked how the *in vitro* activated Dux network compared to gene expression throughout PD *in vivo*. Strikingly, the in vivo RNA-seq data of the developmental stages ^21^ confirmed an early 2C specific expression for Dux, followed by a slightly shifted activation pattern for all direct Dux targets except for *Rxrg* (Figure 4f). However, it is of note that *Rxrg* is significantly upregulated in the *in vitro* DuxOE from which the network is inferred (Supp. Table 5), pointing to both the similarities and differences between the *in vivo* 2C and *in vitro* 2C-like stages induced by Dux. In conclusion, these data show that beyond identifying specific target genes of individual TFs, TOBIAS can infer biological insight by predicting entire TF activation networks.

Notably, many of the predicted Dux binding sites (40%) are not annotated to genes (Figure 4g), raising the question what role these sites play in ZGA. Dux is known to induce expression of repeat regions such as LTRs ^5^ and consistently, we found that more than half of the DUX-bound sites without annotation to genes are indeed located within known LTR sequences (Figure 4g) which were transcribed both *in vitro* and *in vivo* (Figure 4h). Interestingly, we additionally found that 28% of all non-annotated Dux binding sites overlap with genomic loci encoding LINE1 elements. Although LINE1 expression does not appear to be altered in mESC cells, there is a striking pattern of increasing LINE1 transcription from 4C-8C (Figure 4h) *in vivo*, pointing to a possible role of LINE1 regulation throughout PD. Finally, we found a portion of the Dux binding sites which do not overlap with any annotated gene nor with putative regulatory repeat sequences, even though transcription clearly occurs at these sites (Figure 4h; bottom). One example is a predicted Dux binding site on chromosome 13, which coincides with a spliced region of increased expression between control mESC/DuxOE and comparable high expression in 2C, 4C and 8C (Supp. Figure 6). These data clearly indicate the existence of novel transcribed genetic elements, the function of which remains unknown, but which are likely controlled by Dux and could play a role during PD.

In conclusion, TOBIAS predicted the exact locations of Dux binding in promoters of target genes, and could unveil how Dux initiates TF-activation networks and induces expression of repeat regions. Importantly, these data further show that TOBIAS can identify any TFBS with increased binding, not only those limited to annotated genes, which aids in uncovering novel regulatory genetic elements.

## Discussion

### Footprint scores reveal true characteristics of protein binding

To the best of our knowledge, this is the first application of a DGF approach to visualize gain and loss of individual TF footprints in the context of time series, TF overexpression, and TF-DNA binding for a wide-range of TFs in parallel. Importantly, we found that these advances could in large part be attributed to the framework approach we took in developing TOBIAS, which enabled us to simultaneously compare global TF binding across samples and quantify changes in TF binding at specific loci. The modularity of the framework also allowed us to apply a multitude of downstream analysis tools to easily visualize footprints and gain even more information about TF binding dynamics as exemplified by the discovery of the Dux TF-activation network.

The power of this framework to handle time-series data becomes especially apparent when correlating the TOBIAS-based prediction of TF binding to RNA-seq data from the same time points. For instance, TOBIAS could infer when the maternally transferred TF SALL4 is truly active while its gene expression pattern alone does not allow to make such conclusions. Along this line, TOBIAS is also powerful in circumstances where gene expression of a particular TF appears to be anticorrelated with its binding activity. It is tempting to speculate that TFs for which footprinting scores are low, even though their RNA expression is high, might act as transcriptional repressors, because footprinting relies on the premise that TFs will increase chromatin accessibility around the binding site. In support of this hypothesis, recent investigations have suggested that repressors display a decreased footprinting effect in comparison to activators ^37^. Therefore, the integration of ATAC-seq footprinting and RNA-seq is an important step in revealing additional information such as classification TFs into repressors and activators, as well as the kinetics between expression and binding.

### Species-specific TFs use common ZGA motifs in mice and human

By integration of human and murine TF activities using both differential footprinting and species-specific TFBS overlaps, our analyses revealed that the majority of TF motifs are active at corresponding timepoints of human and mouse ZGA. This is not necessarily surprising since homologous TFs that exert the same functions usually use similar motifs (e.g Pou2f1/POU2F1, Otx1/OTX1 and/or Foxa3/FOXA3). Interestingly though, we found that this is not the case for all TF motifs. We found that the human RHOXF1 motif (Figure 2b) is likely not utilized by Rhox proteins in mice even though more than 30 Rhox genes exist. Evidently, throughout multiple duplications, Rhox genes seem to have obtained other functionalities in mouse ^38^ in comparison to the two human RHOX genes that are expressed in reproductive tissues ^39^. Therefore, although we found the human RHOXF1 motif to be highly active in mice, this motif is most likely utilized by other proteins such as the mouse specific Obox proteins. In support of this conclusion, expression patterns of Obox proteins appear to be tightly regulated during PD ^40^ (^21^). High expression of Obox 1/2/5/7 is observed from the zygote to 4C stage, while Obox3/6/8 are expressed and peak at later stages (Supp. Table 4). Notably, there is a significant sequence similarity of the homeobox domains but not in the other parts of the RHOXF1 and Obox protein sequences, which supports the similarity in binding specificity. Although the potential functional overlap of RHOXF1 and Obox factors remains unresolved, our inter-species analysis suggests an unappreciated function of these factors and their targets during PD, warranting an in depth investigation.

In the context of TF target prediction, the power of TOBIAS was particularly highlighted by the fact that the analysis could identify almost all known Dux targets. In addition to coding genes, our analysis disclosed novel Dux binding sites and significant footprint scores at LINE1 encoding genomic loci, which appear to be activated at the 4C/8C stage. This finding is especially interesting because a recent study has shown that LINE1 RNA can interact with Nucleolin and Kap1 to repress Dux expression ^41^. Therefore, our findings give rise to a kinetics driven model in which Dux not only initiates ZGA but also regulates its own termination by a temporally delayed negative feedback loop. Exactly how this feedback loop is controlled remains to be determined.

### Limitations and outlook of footprinting analysis

Despite the striking capability of DGF analysis, some limitations and dependencies of this method still remain. Amongst these is the need of high-quality TF motifs for matching footprint scores to individual TFs with high confidence. In other words, while the binding of a TF might create an effect that can be interpreted as a footprint, without a known motif, this effect cannot be matched to the corresponding TF. This becomes evident in the context of DPPA2/4, a TF described by several groups to act in PD and even upstream of Dux ^34^. DPPA2/4 targets GC rich sequences ^34^, but its canonical binding motif remains unknown. It also needs to be noted that footprinting analysis cannot take effects into account that arise from heterogeneous mixtures of cells wherein TFs are bound in some cells and in others not. Therefore, if not separated, the classification of differential binding will be an observation averaged across many cells, possibly masking subpopulation effects. Recent advances have enabled the application of ATAC-seq in single cells ^42^, but this generates sparse matrices, rendering footprinting approaches on single cells elusive. However, we speculate that by creating aggregated pseudo-bulk signals from large clustered SC ATAC datasets, DGF analysis might also become possible in single cells.

## Conclusions

Here, we have illustrated the TOBIAS framework as a versatile tool for ATAC-seq footprinting analysis which helps to unravel transcription factor binding dynamics in complex experimental settings that are otherwise difficult to investigate. We showed that entire networks of TF binding, which have previously been explored using a combination of omics methods, can be recapitulated to a great extent by DGF analysis, which requires only ATAC-seq and TF motifs. From a global perspective, we provided new insights into PD by quantifying the stage-specific activity of specific TFs. Furthermore, we highlighted the usage of TOBIAS to study specific transcription factors as exemplified by our investigations on Dux. Finally, we used the specific TF target predictions to gain insights into the local binding dynamics of Dux in the context of TF-activation networks, repeat regions and novel genetic elements.

In conclusion, we present TOBIAS as the first comprehensive software that performs all steps of DGF analysis, natively supports multiple experimental conditions and performs visualization within one single framework. Although we utilized the process of PD as a proof of principle, the modularity and universal nature of the TOBIAS framework enables investigations of various biological conditions beyond PD. We believe that continued work in the field of DGF, including advances in both software and wet-lab methods, will validate this method as a resourceful tool to extend our understanding of a variety of epigenetic processes involving TF binding.

## Declarations

### Ethics approval and consent to participate

Not applicable.

### Consent for publication

Not applicable.

### Availability of data and materials

The TOBIAS software is available on GitHub at: https://github.com/loosolab/TOBIAS.

Excerpts of the data analyzed here are accessible for dynamic visualization at: http://loosolab.mpi-bn.mpg.de/tobias-meets-wilson. All raw data analyzed are available from

GEO or ENCODE as described in Methods. The complete TOBIAS output for the analysis of the Dux overexpression dataset can be downloaded from: https://figshare.com/projects/Digital_Genomic_Footprinting_Analysis_of_ATAC-seq_dataset_from_preimplantation_timepoints_via_TOBIAS/69959.

### Authors’ contributions

MB, CK, JK and ML wrote the manuscript. MB, PG, HS, AP, KK, RW, AF and JP performed the bioinformatics analysis. JK, TB and ML directed, coordinated and supervised the work.

## Acknowledgements

We would like to thank the IT-group at MPI-BN for continued support with IT-infrastructure. We would also like to thank Marius Dieckmann, the administrator of the Kubernetes cluster in Gießen, for his support and help in implementing the TOBIAS-Nextflow Cloud version.

## Methods

### Datasets

| Organism | Deposited data | Source | Identifier |
| --- | --- | --- | --- |
| Mouse | ATAC-seq, RNA-seq and ChIP-seq from mESC control and Dux overexpression | <sup>5</sup> | GEO:<br>GSE85632 |
| Mouse | ATAC-seq and RNA-seq from various preimplantation stages | <sup>21</sup> | GEO:<br>GSE66390 |
| Human | ATAC-seq and RNA-seq from various preimplantation stages | <sup>20</sup> | GEO:<br>GSE101571 |

For all public data sets used in this study (see table above), raw files were obtained from the European Nucleotide Archive ^43^ and processed as described in the methods section. See also methods section “Comparison of TOBIAS to existing methods” for links to the ENCODE data used for method validation.

### Processing of ATAC-seq data

Raw sequencing fastq files were assessed for quality, adapter content and duplication rates with FastQC v0.11.7, trimmed using cutadapt ^44^ and aligned with STAR v2.6.0c ^45^ (parameters: “--alignEndsType EndToEnd --outFilterMismatchNoverLmax 0.1 -- outFilterScoreMinOverLread 0.66 --outFilterMatchNminOverLread 0.66 –outFilterMatchNmin 20 --alignIntronMax 1 --alignSJDBoverhangMin 999 --alignEndsProtrude 10 ConcordantPair --alignMatesGapMax 2000 --outMultimapperOrder Random --outFilterMultimapNmax 999 -- outSAMmultNmax 1”) to either the mouse or human genome using Mus_musculus.GRCm38 or Homo_sapiens.GRCh38 versions from Ensembl ^46^. Accessible regions were identified by peak calling for each sample separately using MACS2 (parameters: “--nomodel --shift -100 -- extsize 200 --broad”) ^47^. Peaks from each sample were merged to a set of union peaks across all conditions using “bedtools merge”. Each union peak was annotated to the transcriptional start site of genes (GENCODE ^48^) in a distance of -10000/+1000 from the TSS using UROPA^49^.

### Processing of RNA-seq data

Raw reads were assessed for quality, adapter content and duplication rates with FastQC v0.11.7, trimmed using cutadapt ^44^ and aligned with STAR v2.6.0c ^45^ (parameters: “-- outFilterMismatchNoverLmax 0.1 --outFilterScoreMinOverLread 0.9 -- outFilterMatchNminOverLread 0.9 --outFilterMatchNmin 20 --alignIntronMax 200000 -- alignMatesGapMax 2000 --alignEndsProtrude 10 ConcordantPair --outMultimapperOrder Random --outFilterMultimapNmax 999”) to either the mouse or human genome using Mus_musculus.GRCm38 or Homo_sapiens.GRCh38 versions from Ensembl ^46^. Differentially expressed genes were identified using DESeq2 v1.22 ^50^. Only genes with a minimum log2 fold change of ±1, a maximum Benjamini–Hochberg corrected P-value of 0.05 and a minimum combined mean of five reads were classified as significantly differentially expressed.

### Processing of ChIP-seq data

Raw sequencing files in fastq format were quality assessed by Trimmomatic by trimming reads after a quality drop below a mean of Q15 in a window of 5 nucleotides ^51^. All reads longer than 15 nucleotides were aligned versus the mouse genome version mm10, keeping just unique alignments (parameters: --outFilterMismatchNoverLmax 0.2 --outFilterScoreMinOverLread --outFilterMatchNminOverLread 0.66 --outFilterMatchNmin 20 --alignIntronMax 1 -- alignSJDBoverhangMin 999 --outFilterMultimapNmax 1 --alignEndsProtrude 10 ConcordantPair) by using the STAR mapper ^45^. Read deduplication was done by Picard (http://broadinstitute.github.io/picard/).

### Processing of transcription factor motifs

TF motifs were downloaded from JASPAR CORE 2018 ^52^, the JASPAR PBM HOMEO collection and Hocomoco V11 ^53^ databases. We further included the human ARGFX_3 motif from footprintDB 54 which originates from a HT-SELEX assay 55. In annotation to the Dux/Dux4 motifs of JASPAR and Hocomoco, we also included two TF motifs for MDUX/DUX4 created using MEME-ChIP ^56^ with standard parameters on the ChIP-seq peaks of ^35^ (GSE87279).

JASPAR motifs were linked to Ensembl gene ids by mapping the provided “Uniprot id” to the “Ensembl gene id” through biomaRt ^57^. Hocomoco motifs were likewise linked to genes through the provided HGNC/MGI annotation. Due to the redundancy of motifs between JASPAR and Hocomoco, we further filtered the TF motifs to one motif per gene, preferentially choosing motifs originating from mouse/human respectively. For each TOBIAS run, we created sets of expressed TFs as estimated from RNA-seq in the respective conditions. This amounted to 590 motifs for the dataset on human preimplantation stages, 464 motifs for the dataset on mouse preimplantation, and 459 for the DuxOE dataset.

### Maternal genes

Maternal genes for human and mouse were downloaded from the REGULATOR database ^22^. Entrez gene ids were converted to Ensembl gene ids using biomaRt ^57^ and subsequently matched to available TF motifs as previously explained.

### Overlap of Dux binding sites to repeat elements

Repeat elements for mm10 were downloaded from UCSC (http://hgdownload.soe.ucsc.edu/goldenPath/mm10/database/rmsk.gz). Overlap of Dux sites to individual repeat elements (as seen in figure 4G) was performed using “Bedtools intersect”. The sum of overlaps were counted by repeat class (LINE1/LTR).

### Visualization

All TF-score heatmaps were generated by R Version 3.5.3 and complex heatmap package version 3.6 ^58^. Individual gene views were generated by loading TOBIAS output tracks into IGV version 2.6.2 ^59^ or using the TOBIAS PlotTracks module, which is a wrapper for the svist4get visualization tool ^60^. TF networks were drawn with Cytoscape version 3.7.1 ^61^. Heatmaps of genomic signal density were generated using Deeptools version 3.3.0 ^62^. All other figures, such as footprint plots, volcano plots and motif clustering dendrograms were generated by the TOBIAS visualization modules as described below.

### The TOBIAS framework

In developing TOBIAS, we found that there were six main areas of DGF which had not been comprehensively addressed in the context of ATAC-seq footprinting analysis:

- All-in-one framework including bias correction, footprinting, quantification of protein binding and visualization
- Investigation of TF binding on a global level (which TFs are more bound globally) as well as the locus-specific level (which TF binds to which genomic locations including statistics on differential binding)
- Consideration of the redundancy and similarity of known TF binding motifs in the context of footprinting
- A scoring model for TF-DNA binding taking into account the potential lack of a canonical footprint effect
- Comparison and quantification of TF binding activity within complex experimental settings (multiple conditions or time series)
- Automated workflows for recurring analysis tasks

Modules enabling these individual analysis steps are included in the TOBIAS package, which is publicly available at Github (https://github.com/loosolab/TOBIAS) as well as on PyPI and Bioconda. Besides the examples given in the repository README, we also provide a Wiki (https://github.com/loosolab/TOBIAS/wiki) which introduces some of the individual software modules. We used the pre-defined workflows in Snakemake and NextFlow to run the full analysis. The single modules are explained in more detail below.

#### Bias correction (TOBIAS ATACorrect module)

Each Tn5-cut site is defined as the 5’ end of the read shifted by +5 at the plus strand and -4 at the minus strand to center the transposase event. Using the mapped reads from closed chromatin, ATACorrect builds a dinucleotide weight matrix ^63^ representing the preference of Tn5 insertion. In contrast to the classical position weight matrix (PWM) the dinucleotide weight matrix (DWM) captures the inter-base relationships which arise due to the palindromic nature of the bias. A background model is similarly built by shifting all reads +100bp as described by^64^.

Reads within open chromatin peaks are then corrected by estimating the expected number of cuts per base pair and subtracting this from the observed cut sites as follows (modified from ^65^):

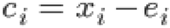

where

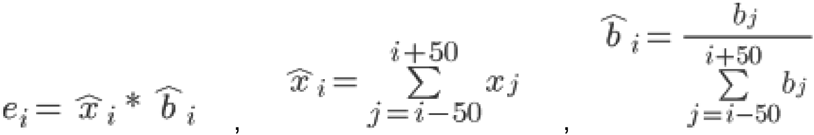

where *x*_*i*_ is the observed number of cuts, *e*_*i*_ is the expected number of cuts, *b*_*i*_ is the calculated bias level, and *c*_*i*_ is the corrected number of cuts at position i. To limit the influence of low-bias positions in the calculation of 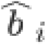, a lower limit is set for *b*_*i*_ by calculating the fit of cutsites vs. bias to a rectified linear unit function (ReLu) in moving 100bp-windows and setting every *b*_*i*_ below the linear fit to 0. This calculation is performed for all base pairs within open chromatin, setting all other positions to 0. Lastly, each *c*_*i*_ is rescaled to fit the original sum of cuts 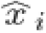 for each window.

#### Footprinting (TOBIAS ScoreBigwig module)

We estimate footprint scores across open chromatin regions by calculating:

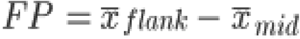

where

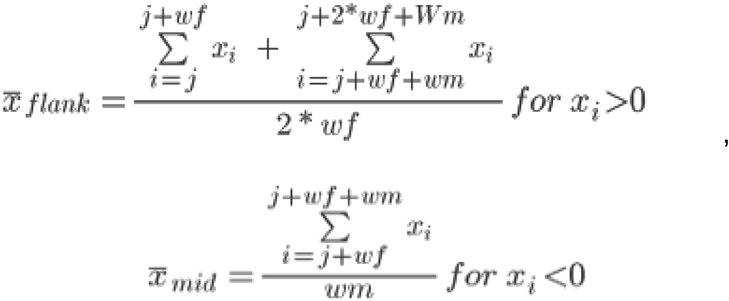

*x*_*i*_ is the number of cuts at position i, *wf* = width of flank in bp, *wm* = width of middle (footprint) in bp. The defaults used are: *wf* = [10;30], *wm* = [20;50].

The term 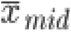 will be negative and will therefore raise the score if there is a high depletion of cuts in the footprint (middle). If there is no depletion, the score will simplify to the mean of cuts in the flanking regions, representing accessibility. It is therefore not necessary to see a canonical footprint shape for the footprint score to be high. The footprint score can be interpreted as higher scores being more evidence that a protein was bound at a given position.

#### Estimation of transcription factor states and pairwise comparison between conditions (TOBIAS BINDetect module)

To match the calculated footprint scores to potential binding sites, TOBIAS BINDetect integrates genomic sequence, footprint scores from several conditions and motifs to identify up- and down regulated TFs based on footprint scores.

In the first step of the algorithm, the MOODS library (https://github.com/jhkorhonen/MOODS ^66^) is used to detect TF binding sites (within peaks) with a p-value threshold of 1e-4. Background base pair probabilities are estimated from the input peak set. Subsequently, each binding site is matched to footprint scores for each condition. Simultaneously, a background distribution of values is built by randomly subsetting peak regions at ∼200bp intervals, and the scores from each condition are normalized to each other using quantile normalization. These values are used to calculate a distribution of background log2FCs for each pairwise comparison of conditions.

Overlaps between the TFBS identified in the first step are quantified by creating a distance matrix of TFs. The distance between a TF pair (TF1;TF2) is calculated as:s

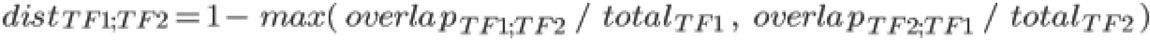

where *total*_*TF*1_ and *total*_*TF*2_ are the total base pairs of all *TF1* and TF2 sites respectively *overlap*_*TF*1;*TF*2_ and is the amount of base pairs of *TF1* which overlap with *TF2* sites. The max-statement ensures that the overlap is calculated with regards to the shortest TF motif.

In the second step of the algorithm, every TF binding site found (for each motif given as input) is split into bound and unbound sites based on a score threshold per condition. The threshold is set at the level of significance of a normal-distribution fit to the background distribution of scores (user-defined p-value). As well as the per-condition split, each site is assigned a log2FC (fold change) per comparison, which represents whether the binding site has larger/smaller footprint scores in comparison. The global distribution of log2FC’s per TF is compared to the background distributions to calculate a *differential binding score*, which is calculated as:

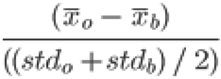

where 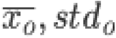 and 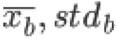 are the means and standard deviations of the observed and background log2FC distributions respectively. A p-value is also calculated by subsampling 100 log2FCs from the background and calculating the significance of the observed change (Python’s scipy.stats.ttest_1samp). By comparing the observed log2FC distribution to the background log2FC, the effects of any global differences due to sequencing depth, noise etc. are controlled.

The differential binding scores and p-values are visualized as a volcano plot per condition-comparison. All TFs with -log10(p-value) above the 95% quantile or differential binding scores smaller/larger than the 5% and 95% quantiles (top 5% in each direction) are colored and shown with labels. Below the plot, hierarchical clustering of the TFBS-distance matrix is shown and all TFs with distances less than 0.5 (overlap of 50% of bp) are colored as separate clusters.

The result of BINDetect is a folder-structure containing an overview of all potential binding sites (as .bed as well as excel-files), the predicted split into bound and unbound sites, and a global overview of differentially bound TFs per condition-comparison.

#### Visualizing aggregate plots and calculation of footprint depth (TOBIAS PlotAggregate module)

Footprints are visualized using the subtool “TOBIAS PlotAggregate”. Aggregate footprints are created by aligning genomic signals centered on all binding sites (taking into account strandedness), to create a matrix of (*n* sites) x (*n* bp). The aggregate signal is calculated as the mean of each column (each bp). The default of +/- 60bp from the motif center was used throughout this manuscript.

The aggregate footprinting depth (FPD), which is applied in Supp. Figure 2c-d, was calculated for each TF as:

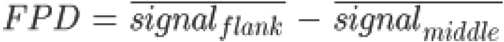

where 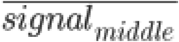 is the mean of the signal centered on the TFBS (30bp) and 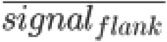 is the mean of the signal in the remaining flanks ([-60;-15] and [+15;+60] bp) (See Supp. Figure 2b). Similarly to the investigations in previous literature ^16^, we applied a mixture model from the Mixtools R package ^67^ to estimate the fractions of TFs with/without measurable footprints (Supp. Figure 2e).

#### Transcription factor binding network (TOBIAS CreateNetwork module)

The TF-TF network for Dux was built by subsetting all binding sites on the following characteristics: Bound in the promoter of a target gene, labeled “Unbound” in Control, labeled “Bound” in DuxOE, and log2FC footprint score increasing for DuxOE vs. Control. All targets were further reduced to only include genes encoding TFs with available motifs. Motifs were matched to genes as explained in the methods section “Processing of transcription factor motifs”. The network was then created using “TOBIAS CreateNetwork”. The result is a network of source and target nodes with directed edges, which in words can be described as: *Source TF* binds in the promoter of *Target TF*.

#### TOBIAS framework output structure

The output generated by the TOBIAS framework is organized in a hierarchical folder structure, which increases clarity of all steps of the analysis. The folder structure specifically organizes input data, pre-processing output like peak-calling and annotation, genomic tracks such as bias correction and footprints, as well as the local and global TF predictions. Particularly, the output for every individual TF investigated is arranged into separate folders containing TF specific plots, annotations and binding predictions. This structure makes it simple to use the output for further downstream analysis, as was showcased in this work. An exemplary output of the complete framework can be found at: https://figshare.com/projects/Digital_Genomic_Footprinting_Analysis_of_ATAC-seq_dataset_from_preimplantation_timepoints_via_TOBIAS/69959.

### Validation

#### Comparison of TOBIAS to existing methods

Although footprinting tools for DNase-seq exist ^68-70 65, 71-73 74^, not all can be applied to paired-end ATAC-seq data. We have focused our comparison on tools which are easily obtainable and installable, do not require ChIP-seq training-data, and are explicitly supporting ATAC-seq. We have additionally added two metrics for “Accessibility” and “PWM score” to compare TOBIAS to other footprinting-free metrics. The validation datasets and usage of existing tools are described in the following sections.

#### Datasets

The TOBIAS framework was benchmarked using ATAC-seq data from four human cell types: GM12878 (GEO: GSE47753), A549 (GEO: GSE114202), K562 (ENA: PRJNA288801) and HEPG2 (ENA: PRJEB30461). ATAC-seq data was trimmed using cutadapt ^44^ and mapped using Bowtie2 ^75^. All reads with a quality score <30 as well as non-proper paired reads were removed. All replicates were merged to one joined .bam-file of reads. Peaks were called using MACS2 ^47^ with parameters “--nomodel --shift -100 --extsize 200 --broad --qvalue 0.01 --broad-cutoff 0.01”. ChIP-seq peak regions (narrowPeak format) were downloaded from ENCODE and associated to motifs from Jaspar CORE 2018 using “MEME Centrimo” ^76^. Only ChIP-seq experiments with motif enrichment > 1.0e-10 (Centrimo E-value) were kept. In case of more than one ChIP-seq experiment for the same target in the same cell type, the one with the highest motif enrichment was chosen. After filtering, there were 12 TFs for A549, 54 TFs for GM12878, 64 TFs for HepG2, and 87 TFs for K562 for a total of 217 ChIP-seq experiments matched to ATAC-seq. Bound binding sites per TF were defined as any TFBS within +/- 50bp from the paired ChIP-seq peak summit. In case of two or more binding sites per peak, the one closest to the summit was set to bound, and others were excluded from the analysis. Unbound binding sites were defined as any TFBS not overlapping any ChIP-seq peak, as well as not overlapping bound sites from any other factors for this cell type. Bound and unbound sites were further filtered to only include TFBS falling within ATAC-seq peaks for the cell type in question.

#### Bias correction approaches

TOBIAS was compared to the existing bias correction methods as follows:

- **seqOutBias (**^**77**^) The seqOutBias software was downloaded from GitHub (https://github.com/guertinlab/seqOutBias). Following the vignette for ATAC-seq, mappability files were created and ATAC-seq reads were corrected for plus/minus strand reads separately. After correction, we further shifted the positive and negative tracks +5 and -4bp respectively, as this was not performed by the tool itself.
- **HINT-ATAC (**^**14**^) The HINT software was downloaded from PyPI as part of the RGT software suite. Bias-correction was performed from the ATAC-seq reads using the command “rgt-hint tracks --bc --bigWig <bam>”.

Aggregate footprints for each method across all (within peaks), bound and unbound binding sites (see explanation above) were visualized using “TOBIAS PlotAggregate”.

#### Footprinting

Existing footprinting tools were applied as follows:

- **msCentipede (**^**78**^) The msCentipede software was downloaded from GitHub (https://github.com/rajanil/msCentipede). For each TF, the binding model was built using the 5000 TFBS with the highest PWM score genomewide. For model learning, the “--mintol” parameter was set to 1e-3 as a tradeoff between accuracy and speed. The resulting models were then used to infer the posterior binding-probability of TFBS in peaks.
- **Wellington (**^**70**^) The pyDNase software was downloaded from PyPI. Footprints in ATAC-seq peaks were estimated using “wellington_footprints.py” with the “-A” option for ATAC-seq mode.
- **PIQ (**^**79**^) The PIQ software was downloaded from Bitbucket (https://bitbucket.org/thashim/piq-single/). The script *bam2rdata*.*r* was used to bring the input .bam-file into the correct data format. Likewise, the script *pwmmatch*.*exact*.*r* was used to predict genomewide TFBS. Finally, footprinting scores for each TF were obtained using the script *pertf*.*r* for each motif/cell type pair. The purity score was taken as the probability for a certain TFBS to be bound.
- **HINT-ATAC (**^**14**^) The HINT software was downloaded from PyPI as part of the RGT software suite. Footprints were identified using the command “rgt-hint footprinting --atac-seq --paired-end --organism=hg38 <bam> <peaks>”. The output of HINT-ATAC footprinting is a.bed-file of footprint ranges ranked by tag count. All TFBS overlapping a footprint with more than 2/3 of the TFBS bases was assumed to be bound and scored using the tag count of the footprint. The rest of the TFBS (within peaks) were set to score 0 (low chance of protein binding). The auROC was calculated based on the ability of these scores to predict true protein binding. It should be noted that this affects the shape of the ROC curve, as all TFBS without overlaps are assumed to have the same probability of being bound. However, this is a characteristic of the method, and HINT-ATAC was therefore evaluated on the same premise as other tools.
- **Accessibility** The “Accessibility” metric is defined as the sum of Tn5 insertions in a 300 basepair window centered at the binding site. This score represents the accessibility of a certain region not taking into account local footprint information.
- **PWM score** The score of the motif-sequence match at the specific TFBS. As this is based on sequence alone, the PWM-score is independent of chromatin accessibility.

Due to high computational times for some tools, the validation was limited to binding sites on human chromosome 1. On the basis of the ChIP-seq labels, the area under the ROC curve (auROC) was used to evaluate the predictive power of each method.

## Supplemental Information

### List of Supplementary Files

*Supplementary File 1: Visualization of different methods for Tn5 bias correction across 36 TFs with matched ChIP-seq. Each page contains footprints for a specific TF across all binding sites (in peaks), bound sites (overlapping ChIP-seq) and unbound sites (not overlapping ChIP-seq) for uncorrected/expected/corrected signals from different bias correction methods*.

*Supplementary File 2: The direct output file of the “TOBIAS BINDetect”-module containing differential binding plots across all pairwise-comparisons of human developmental stages*.

*Supplementary File 3: The direct output file of the “TOBIAS BINDetect”-module containing differential binding plots across all pairwise-comparisons of mouse developmental stages*.

*Supplementary File 4: The direct output file of the “TOBIAS BINDetect”-module containing differential binding plots between control (mESC) and DuxOE samples*.

### List of Supplementary Tables

*Supplementary Table 1: Prediction of transcription factor binding across human 2C/4C/8C/ICM/hESC clustered into co-active TFs. Each transcription factor is further linked to expression of the factor based on RNA-seq*.

*Supplementary Table 2: TOBIAS TF scores for human PD timepoints, correlated to corresponding RNA expression*.

*Supplementary Table 3: Prediction of transcription factor binding across mouse 2C/4C/8C/ICM/mESC clustered into co-active TFs. Each transcription factor is further linked to expression of the factor based on RNA-seq*.

*Supplementary Table 4: Human and Mouse RNA expression for Obox and RHOX/Rhox genes during preimplantation developmental stages*.

*Supplementary Table 5: Full list of the predicted Dux binding sites as well as their change between mESC and DuxOE as predicted by TOBIAS*.

## Figures and figure legends

**Supplementary Figure 1:**
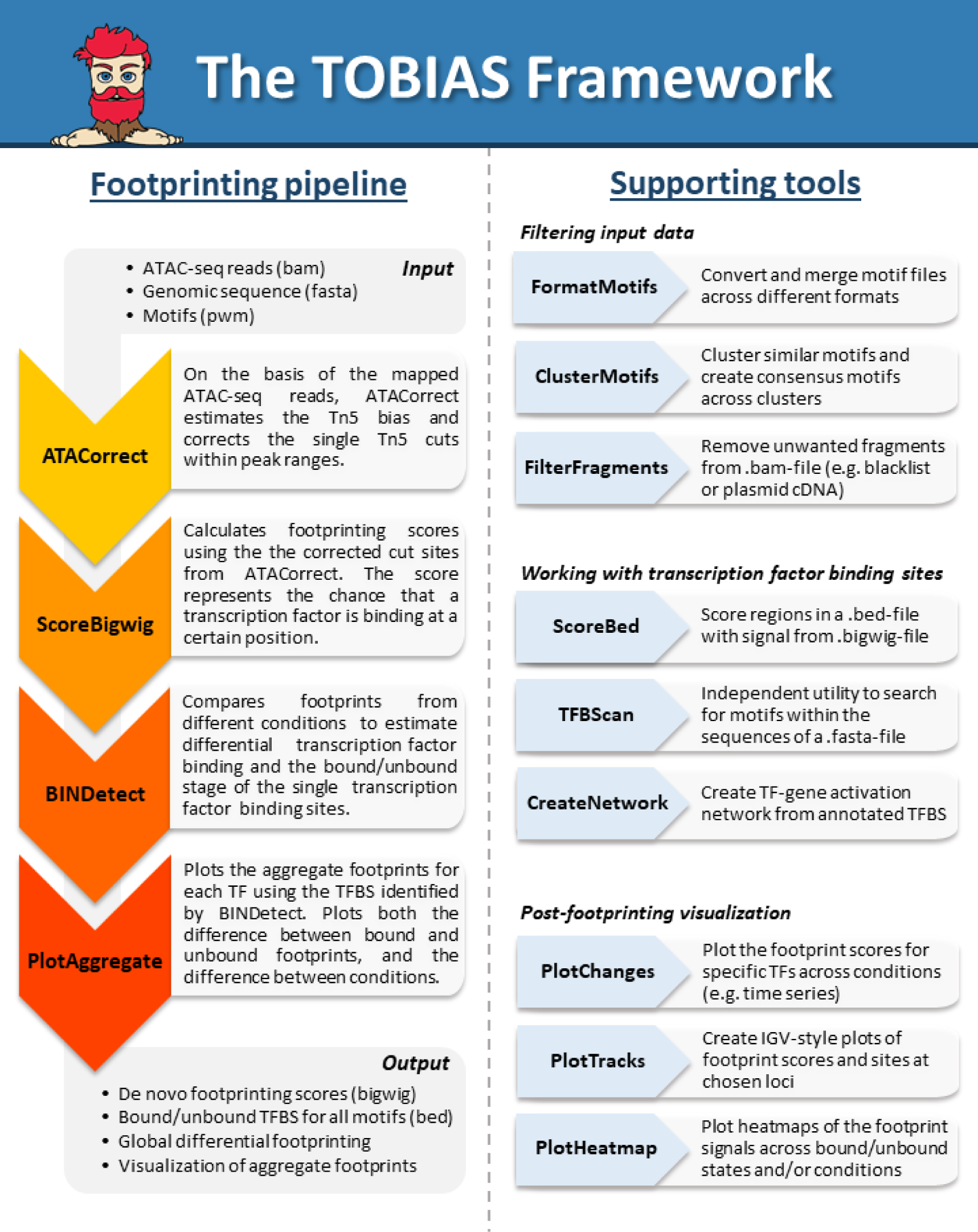
Overview of the TOBIAS framework tools. The TOBIAS tools are intended for use in a standardized pipeline as shown on the left. ATACorrect and ScoreBigWig corrects Tn5 cuts and calculates footprint scores respectively. Next, BINDetect introduces information from different transcription factor binding motifs to predict binding sites both within and across conditions. PlotAggregate can be used to visualize the single footprints. Furthermore, a large variety of supporting tools can be used at different stages of the pipeline, such as pre-filtering of .bam-files using FilterFragments or plotting of locus-specific footprints using PlotTracks.

**Supplementary Figure 2:**
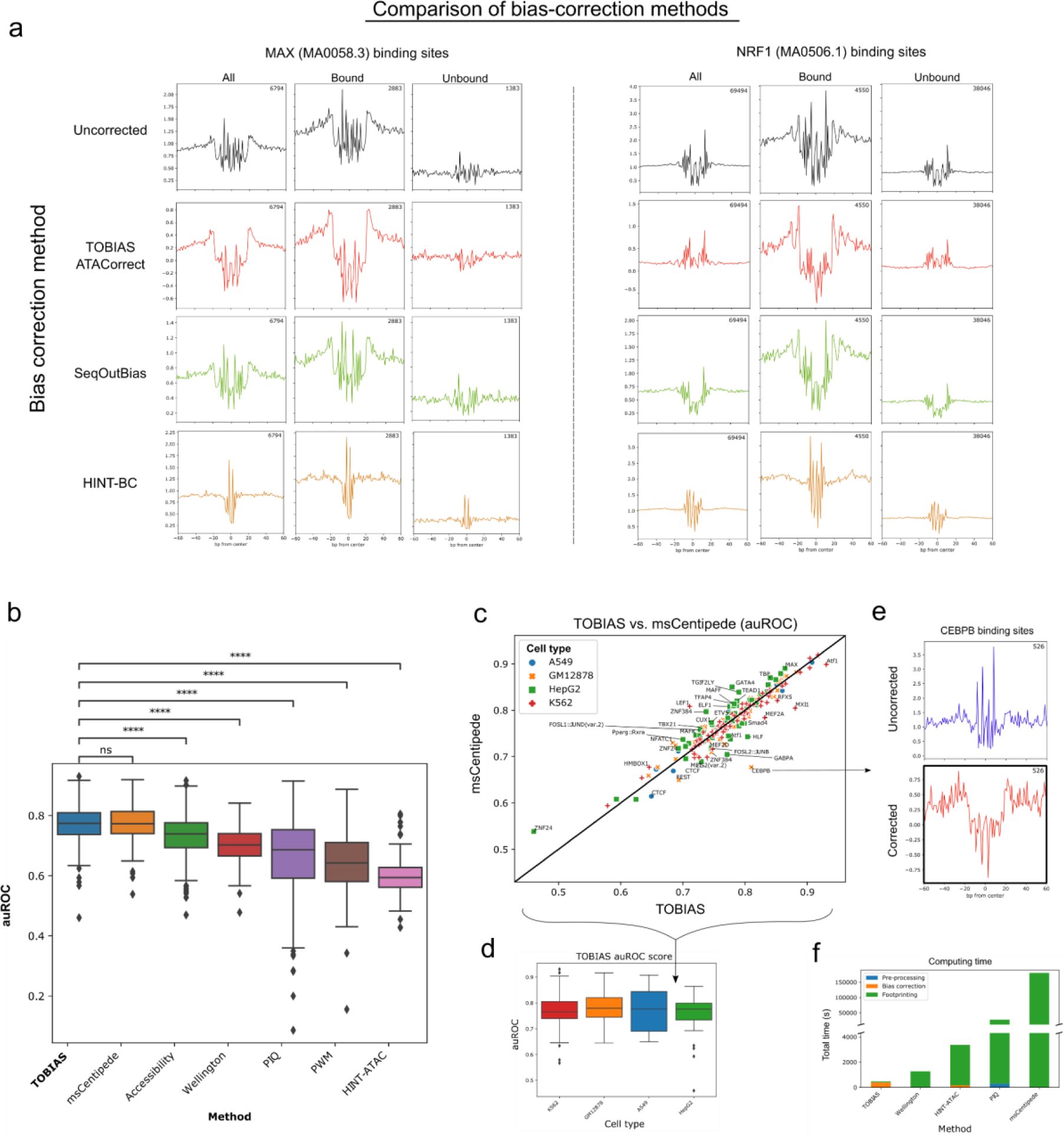
Comparison of existing bias-correction and footprinting methods. **(a) Comparison of aggregate footprints for different bias-correction methods**. Bound and unbound transcription factor binding sites for MAX and NRF1 are shown across uncorrected signal (pileup of Tn5 insertions), TOBIAS ATACorrect, SeqOutBias and HINT-BC correction methods. An overview of all included TFs from cell type GM12878 can be found in Supplementary File 1. **(b) Comparison of predictive ability across different footprinting methods**. The auROC is calculated based on ENCODE ChIP-seq for 217 TFs and compared across methods. Significance (Mann-Whitney U test, **** equals p<=1.0e^-4^) is indicated as asterisk. **(c) Scatterplot comparing the auROC of TOBIAS and msCentipede**. Each point represents one TF, which is colored and marked dependent on cell type. The diagonal line represents equal auROC between TOBIAS and msCentipede. **(d) Validation of TOBIAS across cell types**. The auROC of TOBIAS predictions across cell types K562 (n=67), GM12878 (n=54), HepG2 (n=64) and A549 (n=11). **(e) Aggregate footprints for CEBPB**. The aggregate footprints for true CEBPB binding sites (bound sites verified by ChIP-seq). Whereas the uncorrected ATAC-seq is insufficient to uncover a footprint, the corrected ATAC-seq signal exhibits a clear footprint for CEBPB binding sites. **(f) Comparison of computing times for footprinting tools**. The CPU run time for each tool is measured across the three tasks of “pre-processing” (only for PIQ), “bias-correction” (only for TOBIAS and HINT-ATAC) and footprinting (all tools).

**Supplementary Figure 3:**
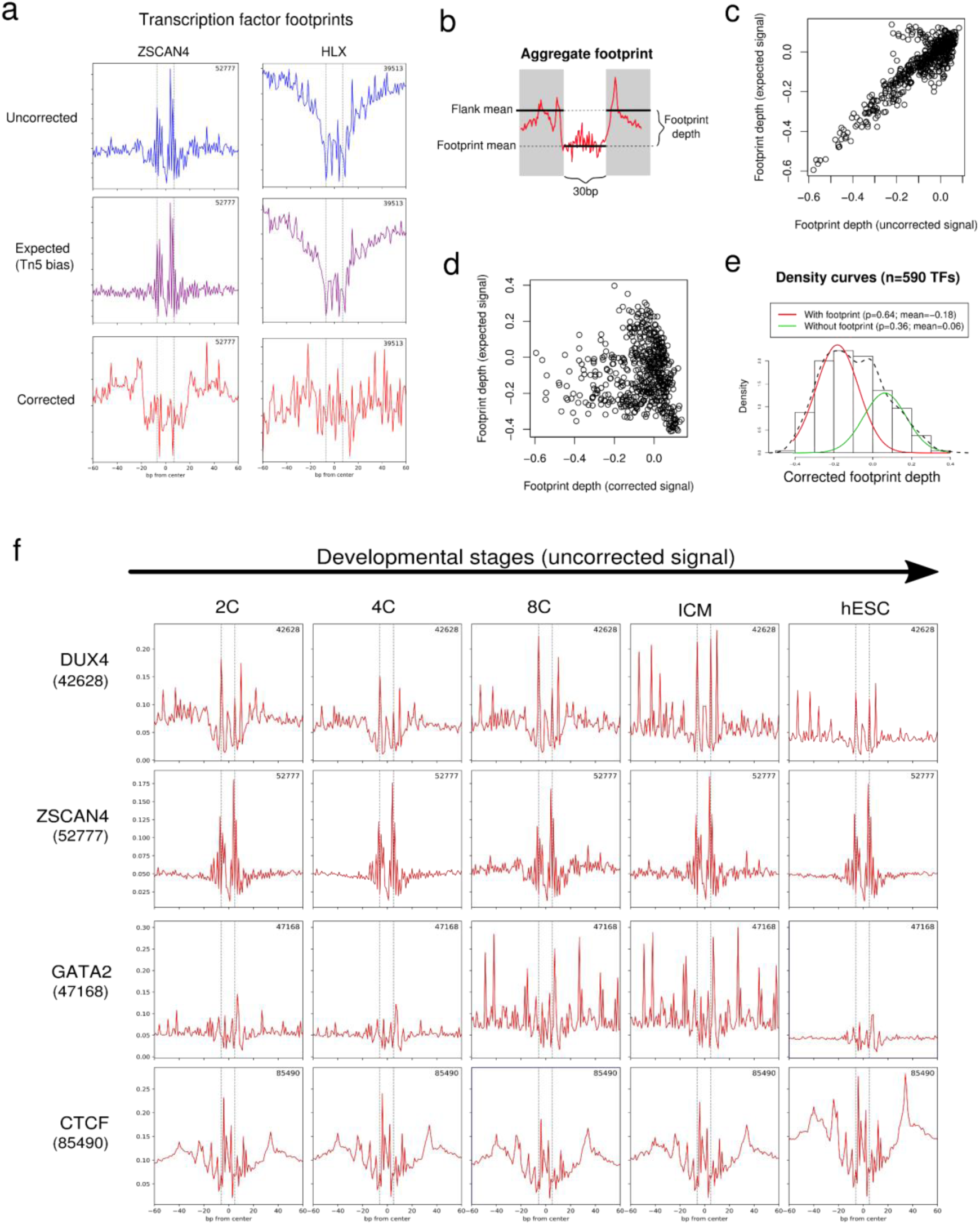
Tn5-bias correction is important for visualization of footprints from ATAC-seq. **(a) Examples of Tn5-bias correction using “expected”-intermediates**. The figure shows the aggregate footprints for transcription factors ZSCAN4 and HLX across the uncorrected, expected and corrected Tn5 signals. The number in the right-hand corner represents number of binding sites included in the plot. **(b) Aggregate footprint depth model**. The footprint depth is calculated using a similar metric as described in ^16^. **(c-d) Uncorrected and corrected Tn5-bias**. The scatter plots show the correlation between depth of footprints for uncorrected vs. expected footprints (c) and corrected vs. expected footprints (d). **(e) Mixture model of all footprinting depths**. The mixture model shows that 65% of motifs fall into the category of a measurable footprint in the aggregated profile. Data is based on 590 motifs in hESC. **(f) A depiction of uncorrected footprint aggregates across time points for transcription factors DUX4, ZSCAN4, GATA2 and CTCF**. In contrast to the corresponding corrected signals seen in Figure 2A, the footprints are hardly visible in the uncorrected aggregates.

**Supplementary Figure 4:**
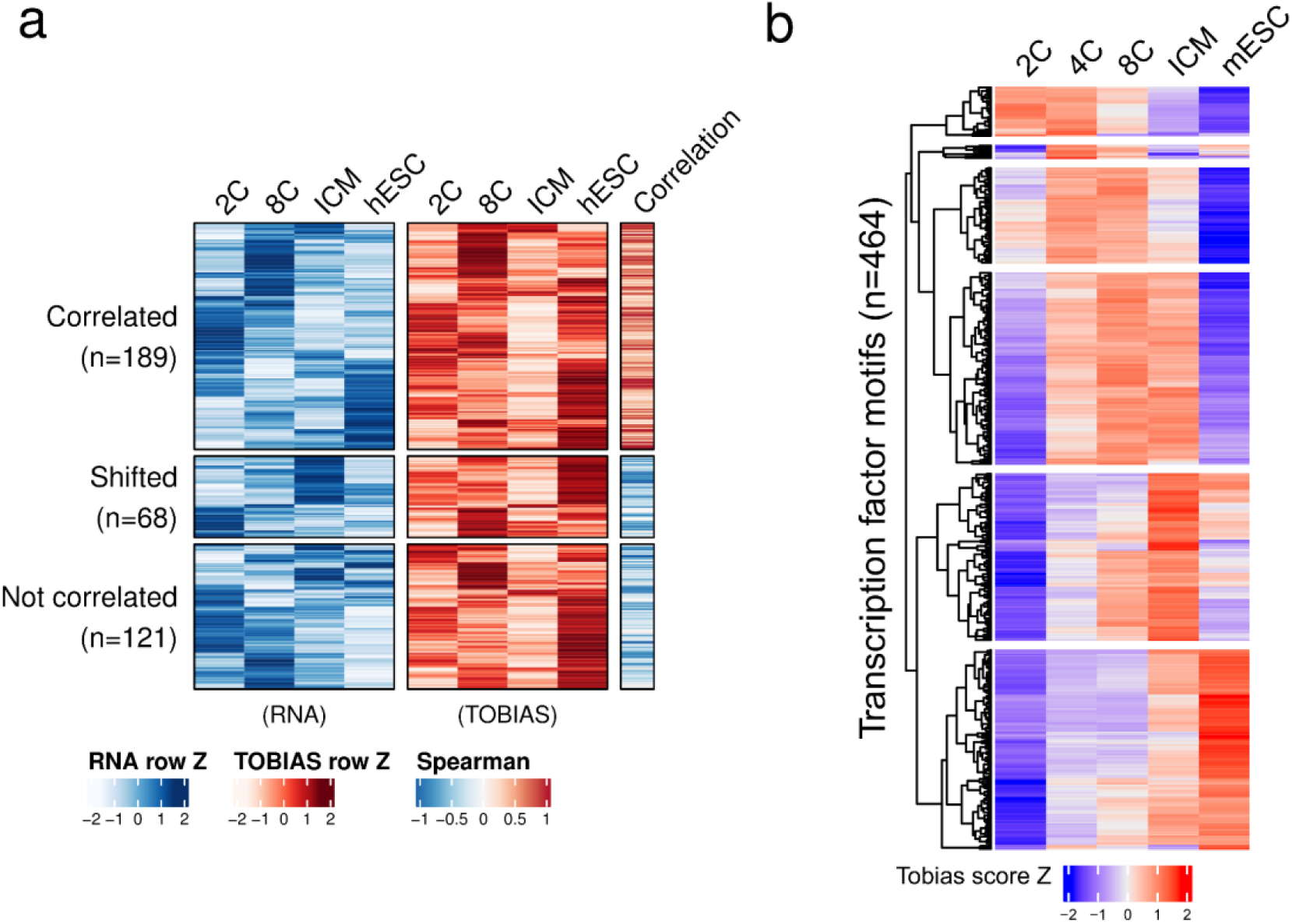
Transcription factor activity and expression during mouse and human development. **(a) Correlation of footprints and RNA-seq**. The left heatmap (blue) depicts expression of transcription factor clusters in the respective human developmental stages. The left heatmap (red) depicts the corresponding TOBIAS scores. Spearman column represents the spearman correlation between TOBIAS/RNA. The TF clusters are grouped into “Correlated” (Spearman≥0.2), “shifted” (RNA max value appears before TOBIAS max value) and “Not correlated” (Spearman<0.2 with no apparent shift in RNA). **(b) Dynamic transcription factor binding during mouse embryonic development**. Similarly to figure 2A, the heatmap depicts the TOBIAS-predicted footprint scores for 464 motifs during the time points 2C, 4C, 8C, ICM and mESC. The rows are clustered into 6 clusters using hierarchical clustering. Individual cluster members are given in Supplementary Table 3.

**Supplementary Figure 5:**
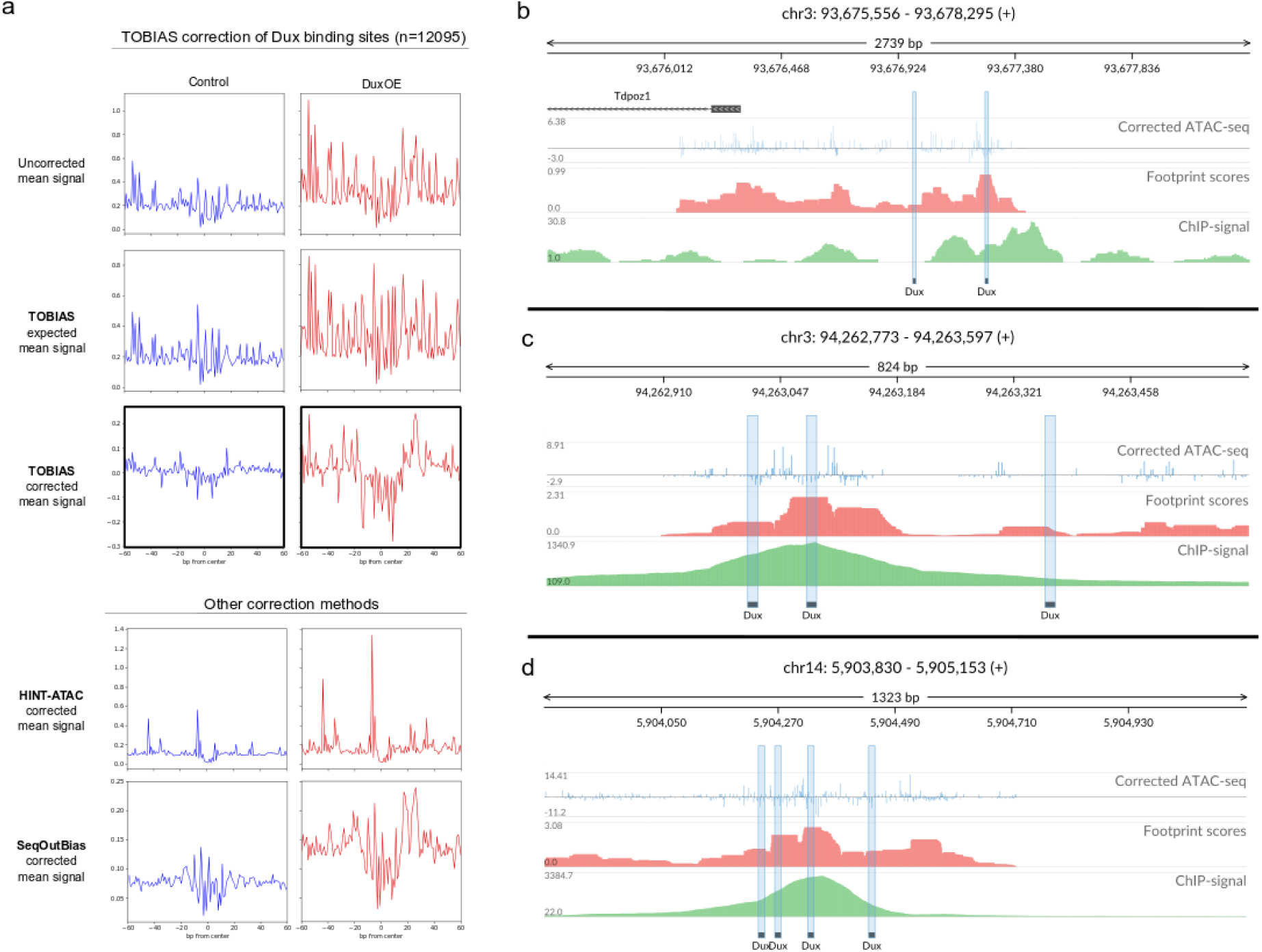
Dux binding is visible as footprints and correlate with ChIP-signal. **(a) Correction of the Dux footprint using different bias correction methods**. The aggregate footprints for 12095 Dux binding sites (within ATAC-seq peaks) are shown between Control and DuxOE conditions. The top three panels depict the uncorrected, expected and corrected signals as calculated by TOBIAS. The bottom panels depict the same sites corrected by either HINT-ATAC or SeqOutBias methods. **(b) A view of the footprinting scores in the promoter of Tdpoz1**. Genomic tracks show corrected ATAC-seq cutsites at 1bp resolution (blue), footprint scores as calculated by TOBIAS (red), and pileup of reads from Dux ChIP-seq of ^5^ (green). Potential Dux binding sites are highlighted in blue. **(c-d) Footprinting correlates with ChIP-signal at multiple genomic loci**. Genomic tracks are the same as described for (a).

**Supplementary Figure 6:**
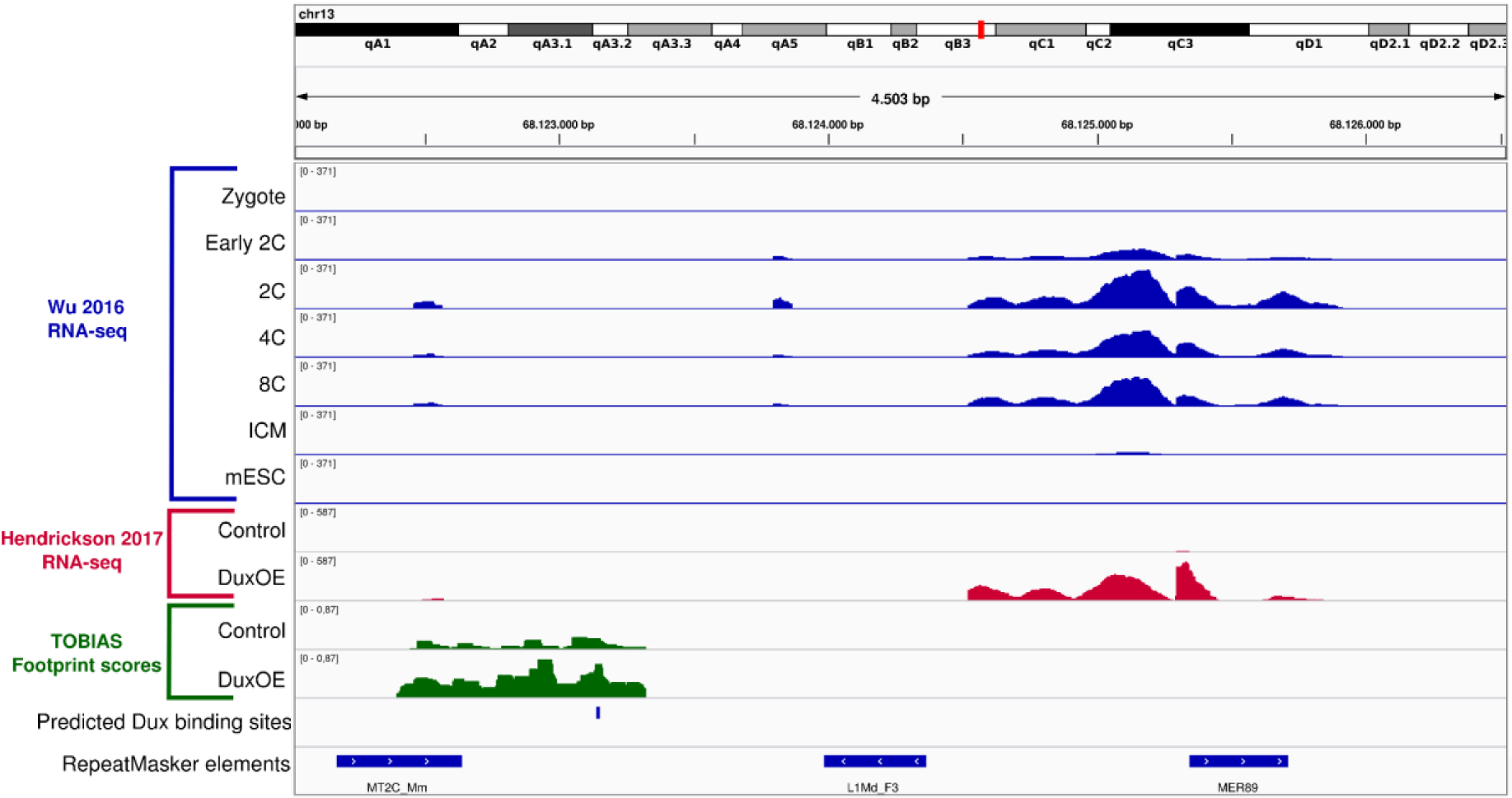
Predicted Dux binding site correlates with increase in expression of close-by non-annotated regions. The figure shows genomic tracks of RNA-seq from ^21^ (blue) and ^5^ (red), TOBIAS footprint scores predicted from ATAC-seq (green) (^5^), predicted Dux binding site as well as known repeats as annotated by RepeatMasker (Smit, AFA, Hubley, R & Green, P. RepeatMasker Open-4.0).

