## Supplemental File 1 for "Beyond accessibility: ATAC-seq footprinting unravels kinetics of transcription factor binding during zygotic genome activation"

### Aggregate footprints for TF BHLHE40 (ENCSR987MTA)

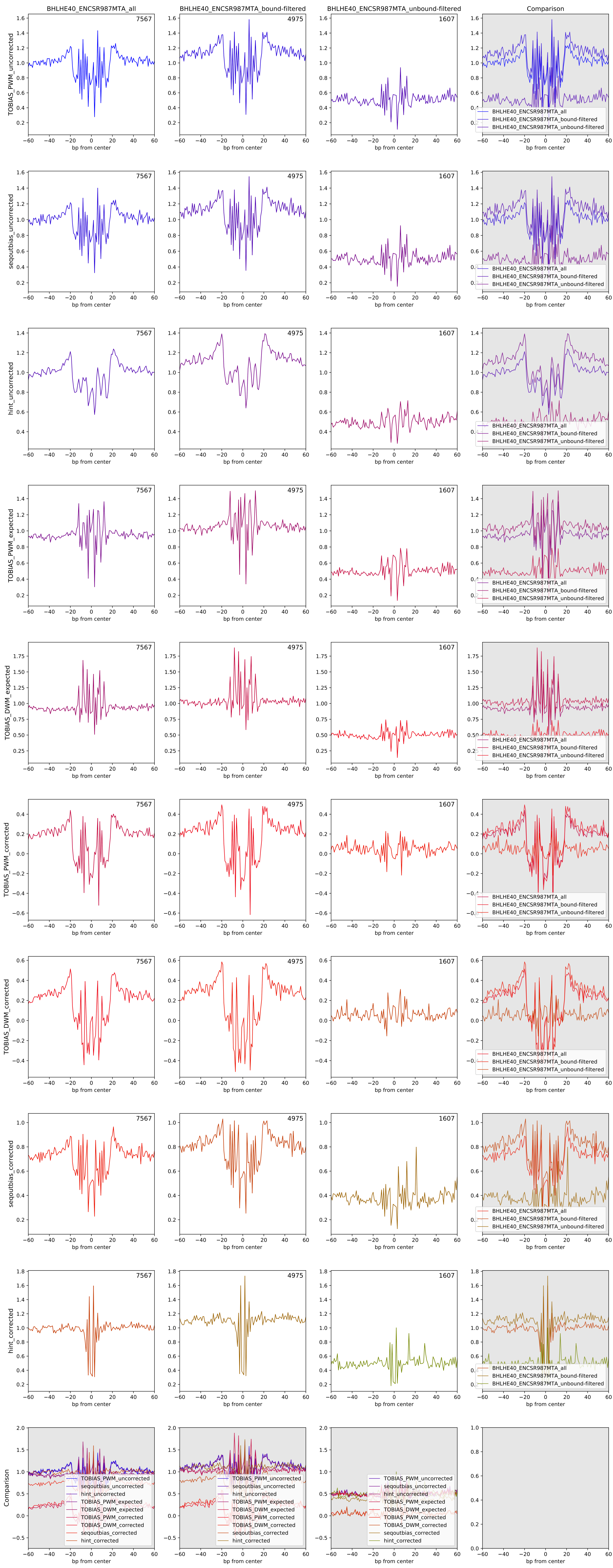

Aggregate footprints for TF CEBPB (ENCSR681NOM)

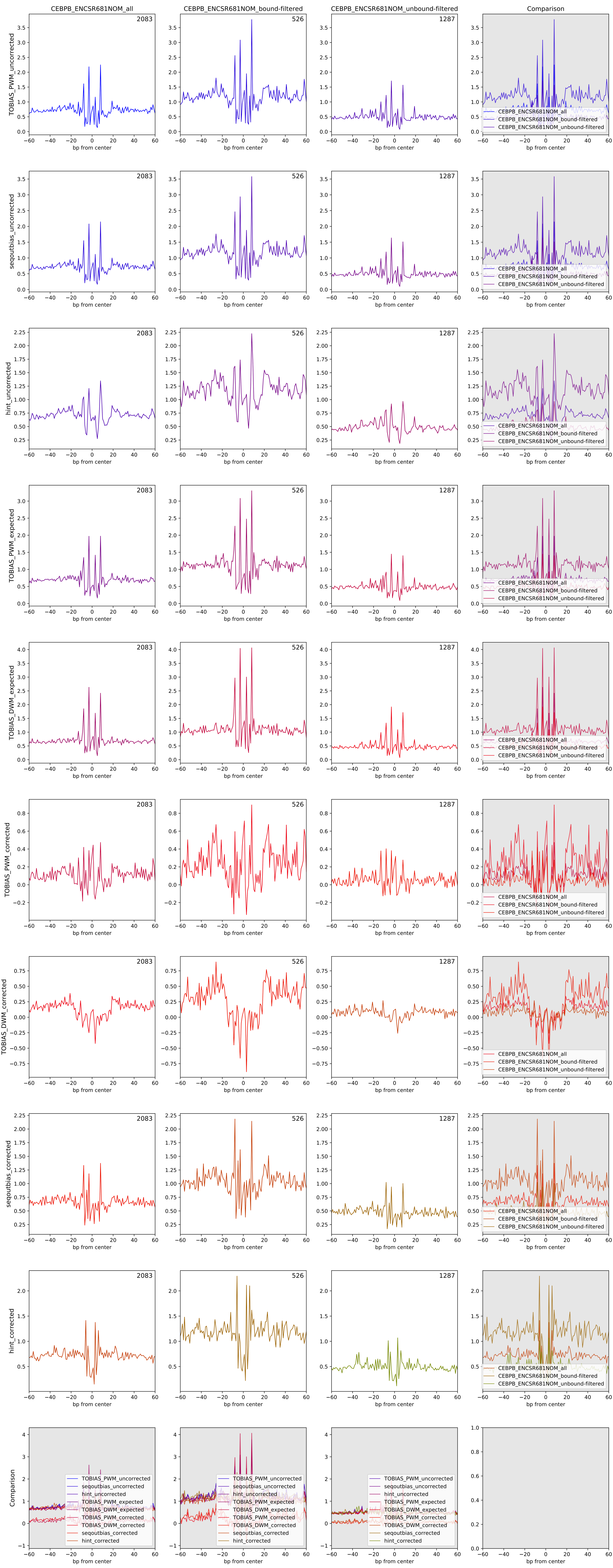

Aggregate footprints for TF CREM (ENCSR839XZU)

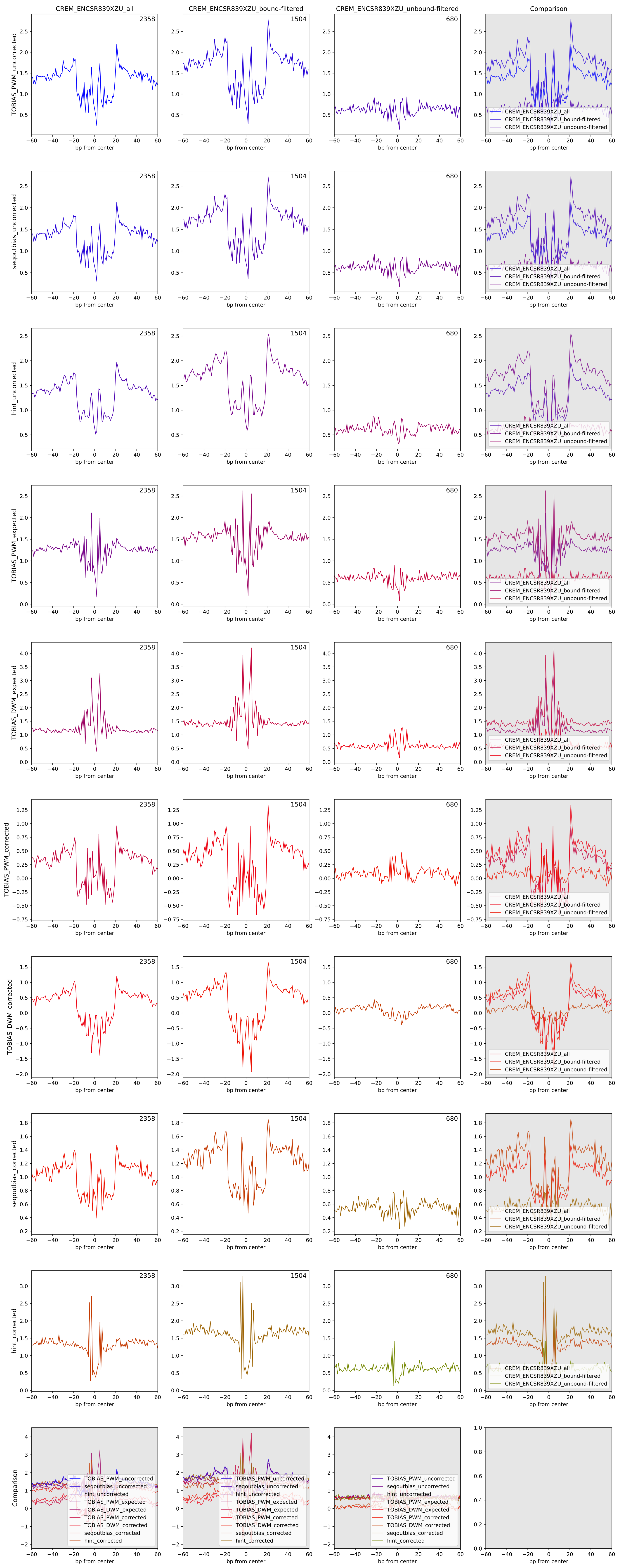

Aggregate footprints for TF CTCF (ENCSR000DZN)

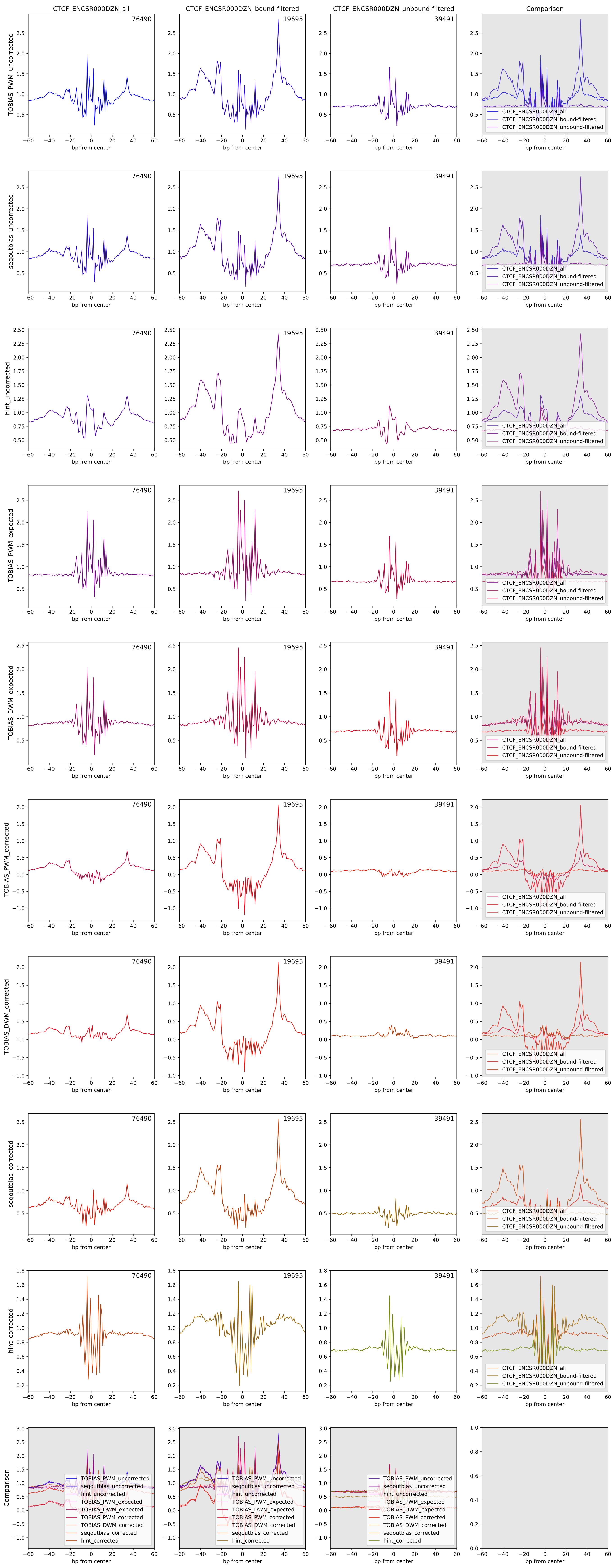

Aggregate footprints for TF EBF1 (ENCSR000DZQ)

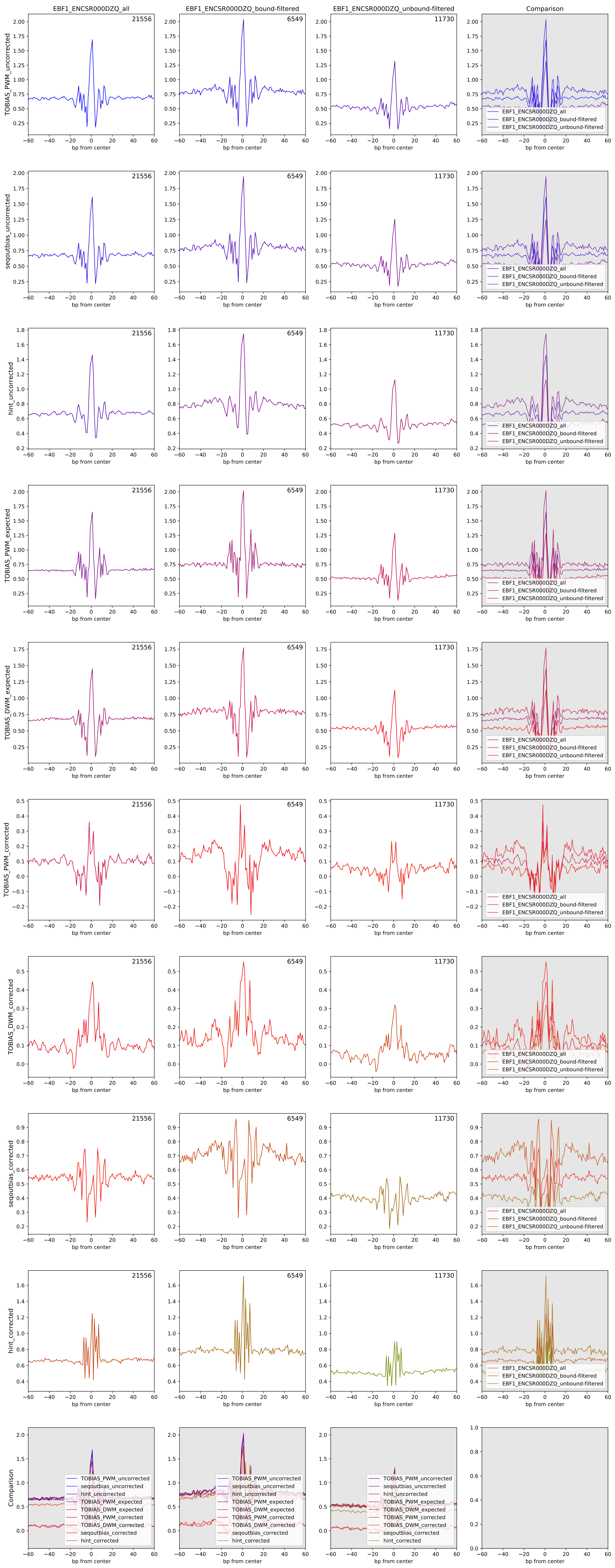

Aggregate footprints for TF ELF1 (ENCSR841NDX)

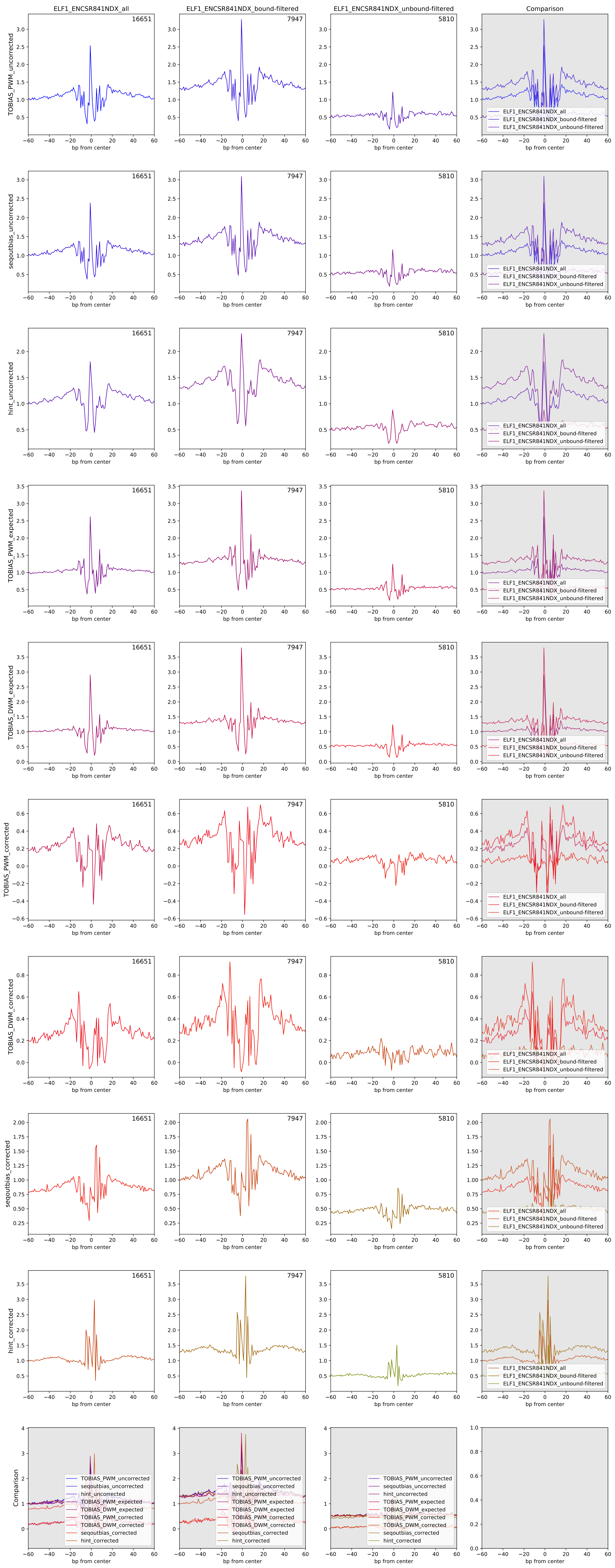

### Aggregate footprints for TF ELK1 (ENCSR000DZB)

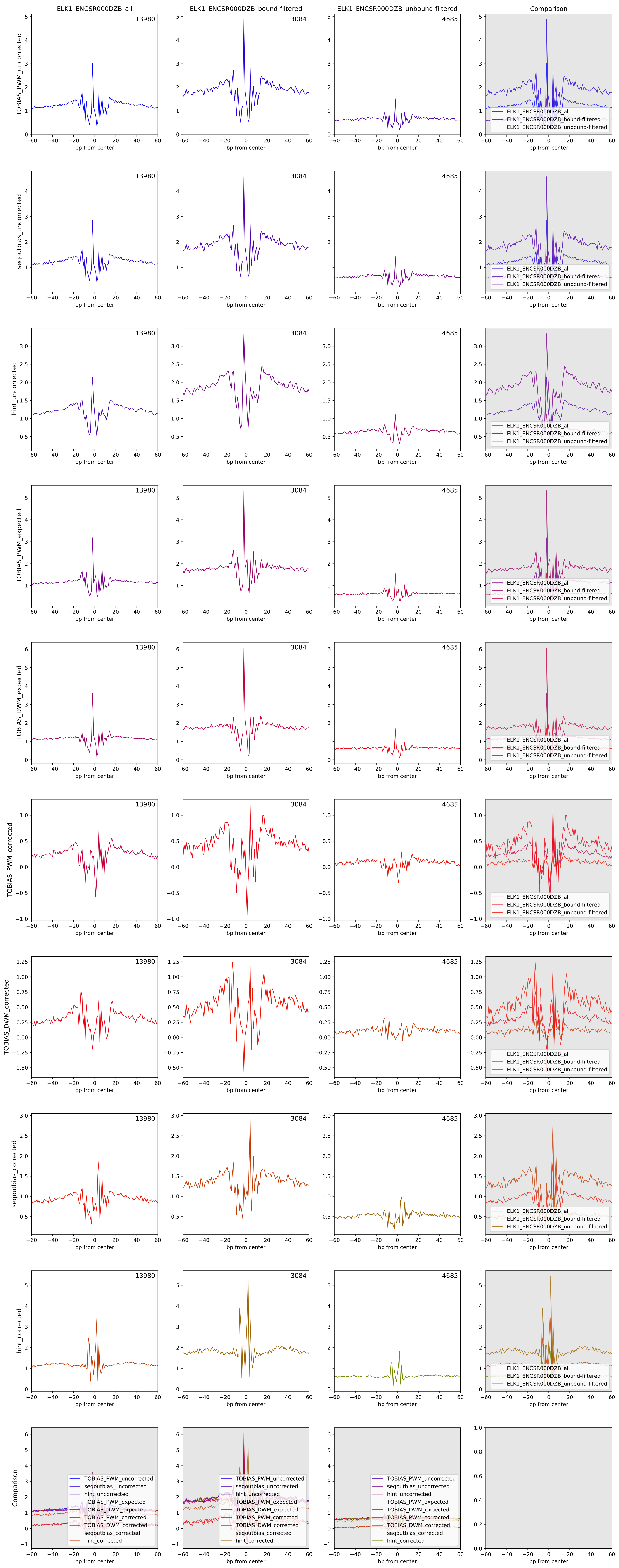

### Aggregate footprints for TF ETS1 (ENCSR000BKA)

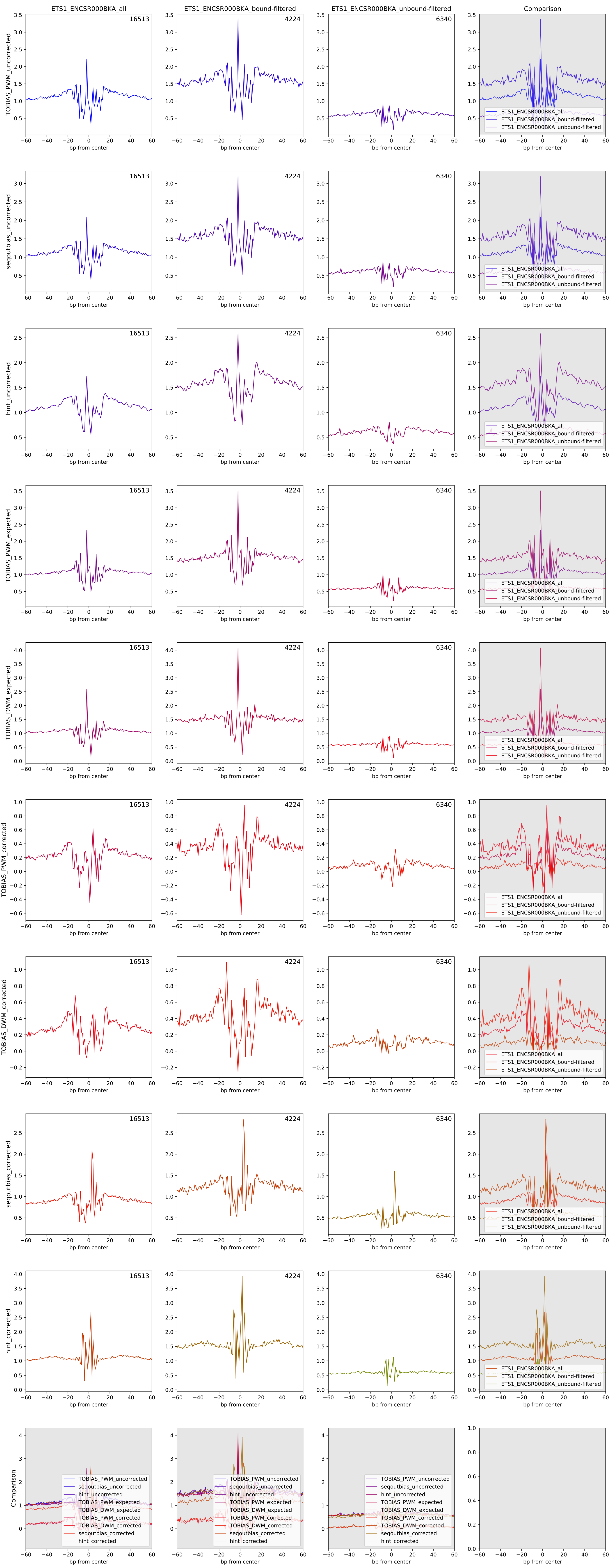

Aggregate footprints for TF ETV6 (ENCSR626VUC)

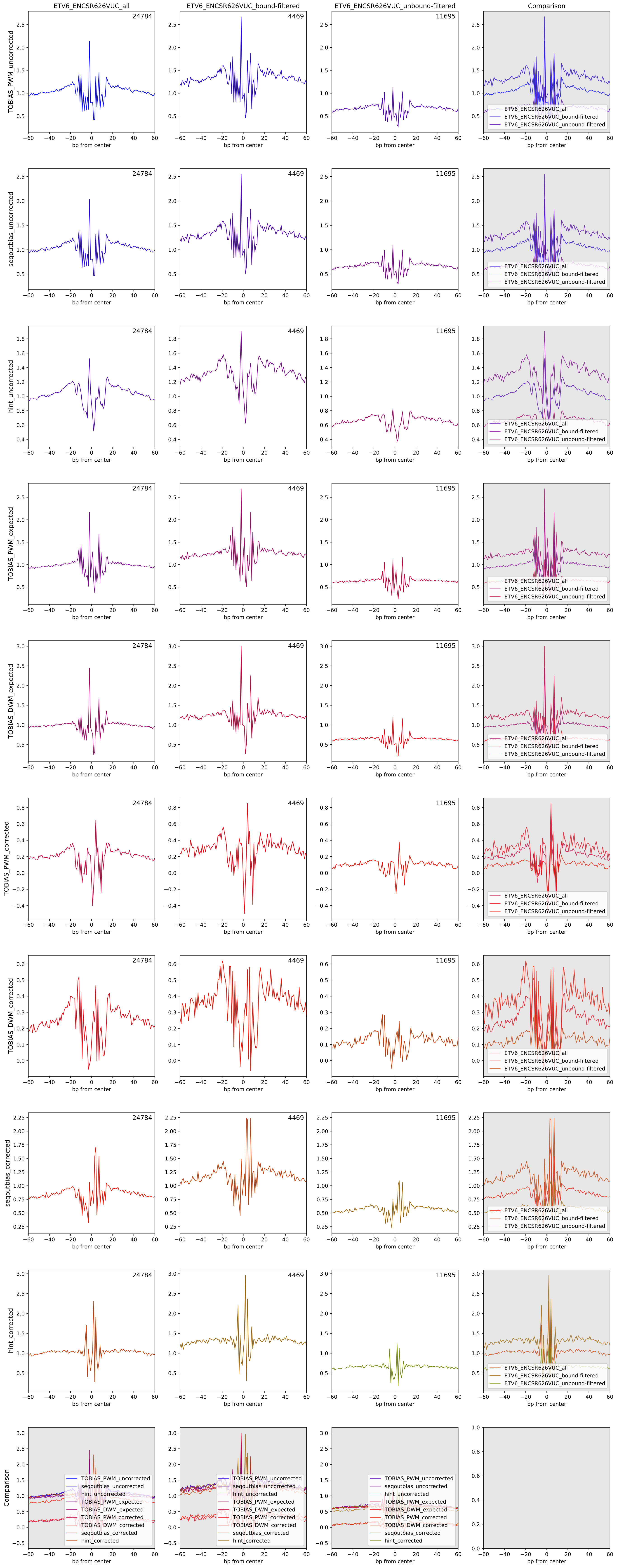

Aggregate footprints for TF GABPA (ENCSR331HPA)

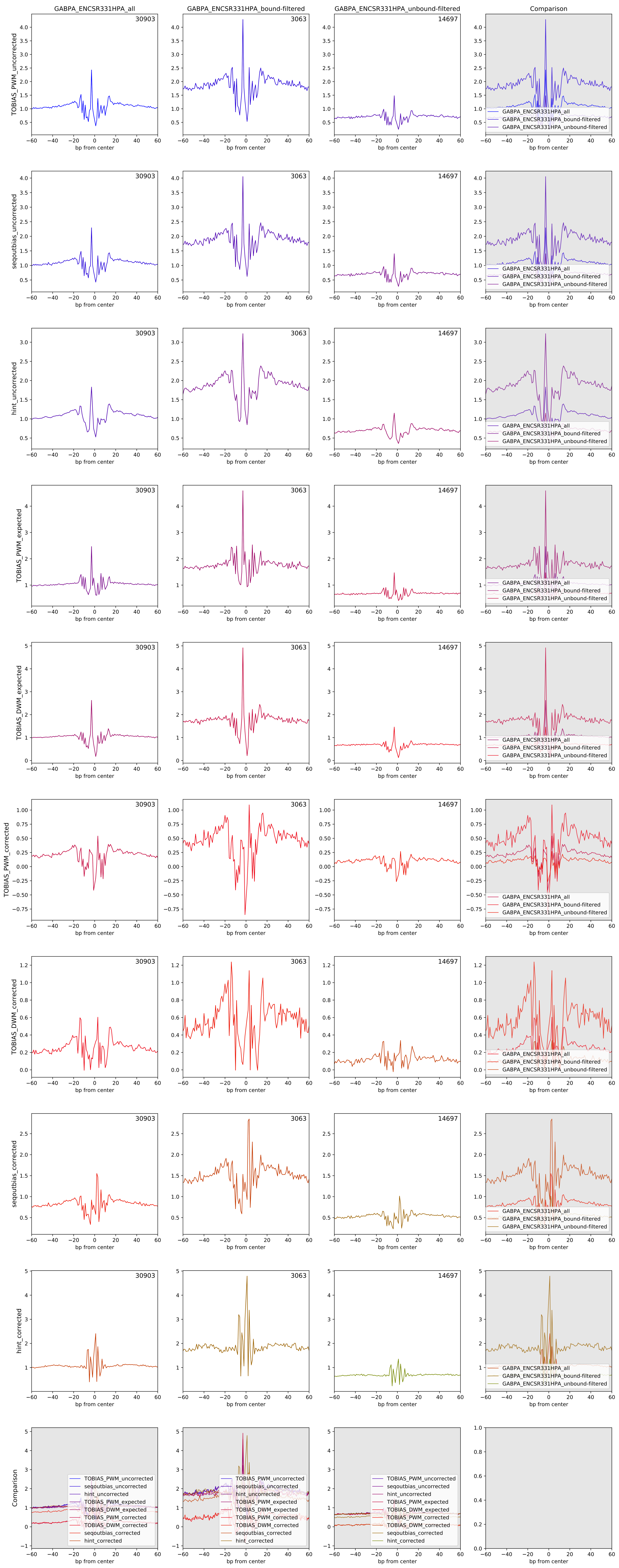

Aggregate footprints for TF HSF1 (ENCSR009MBP)

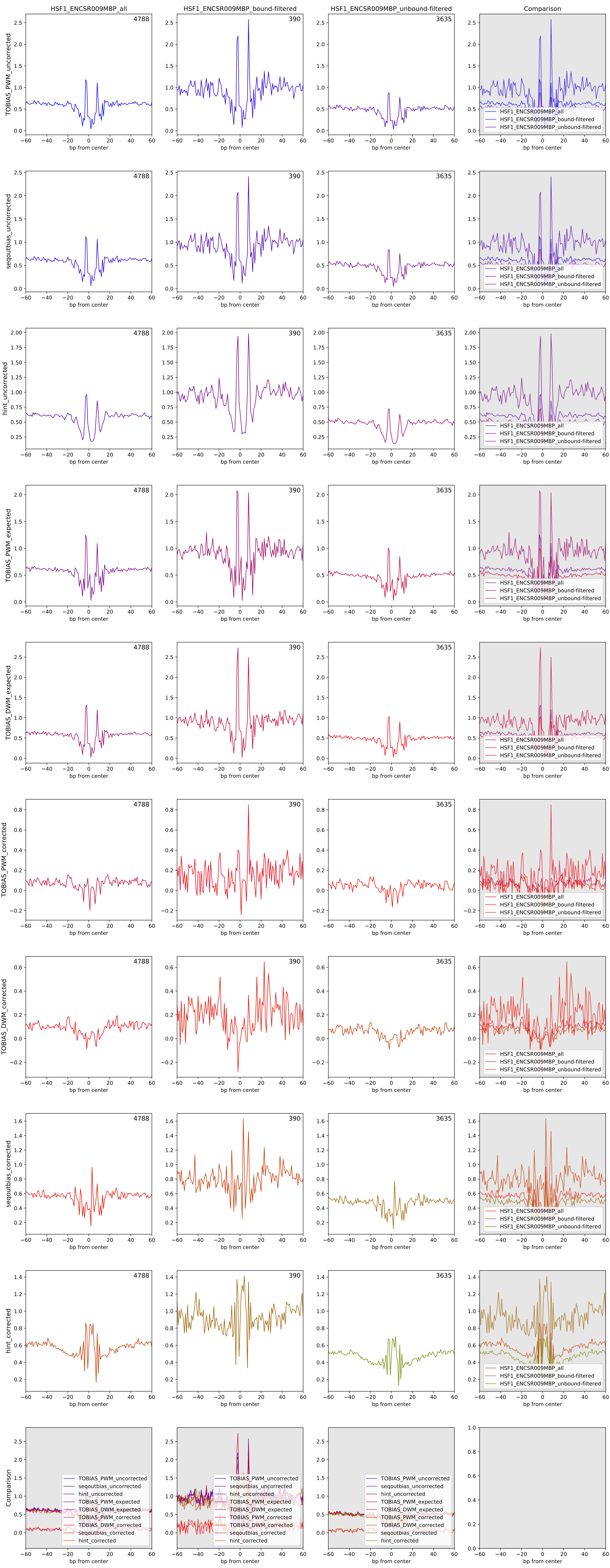

Aggregate footprints for TF JUND (ENCSR000DYS)

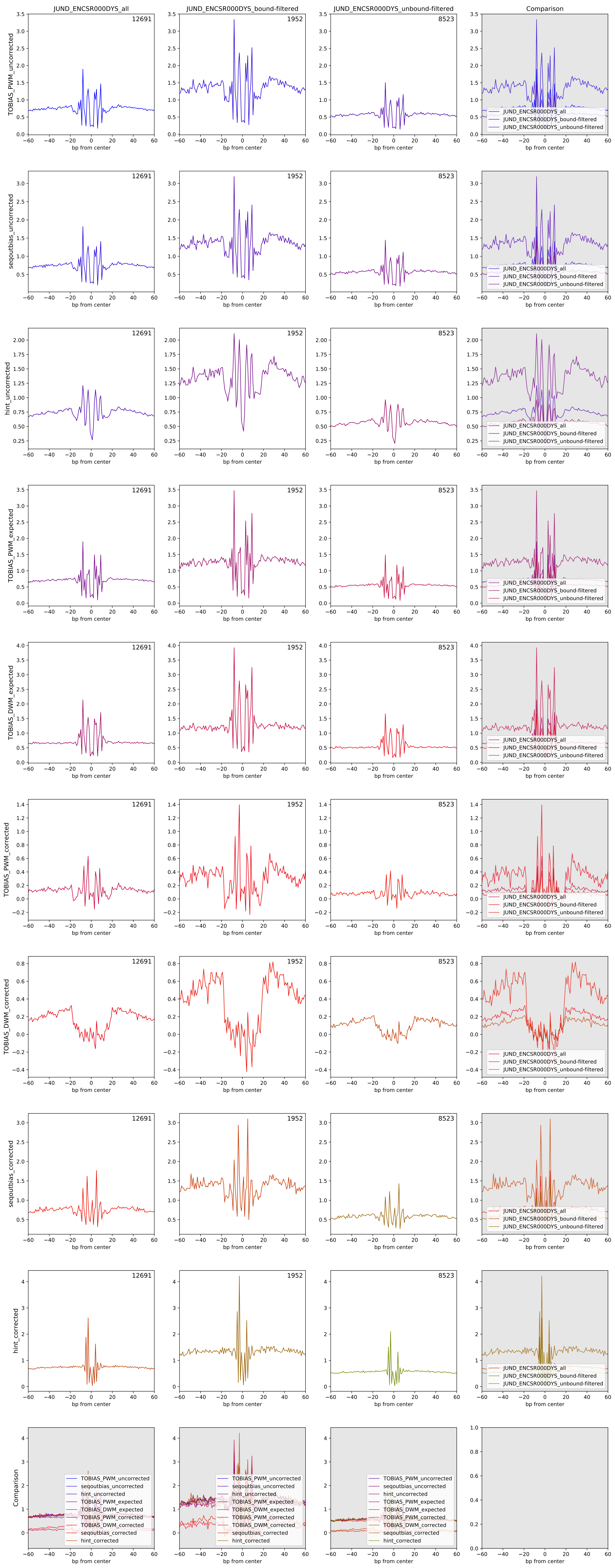

Aggregate footprints for TF MAFK (ENCSR000DYV)

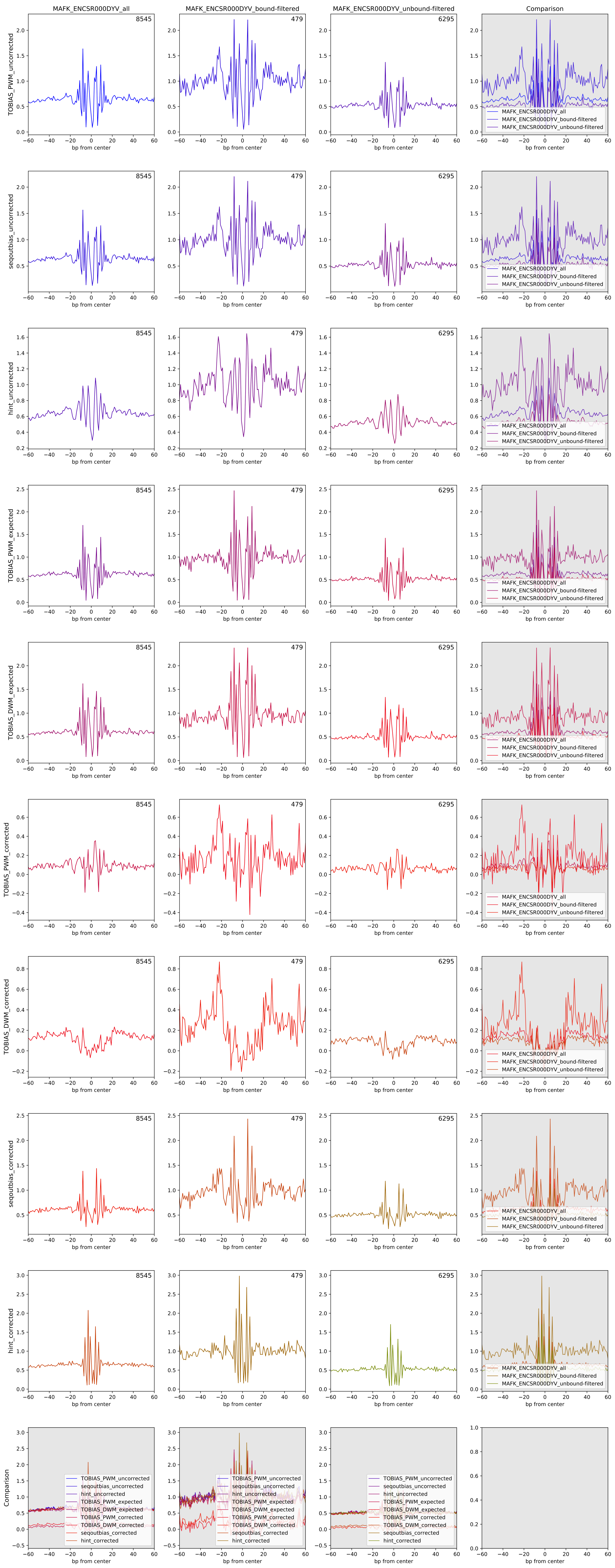

Aggregate footprints for TF MAX (ENCSR000DZF)

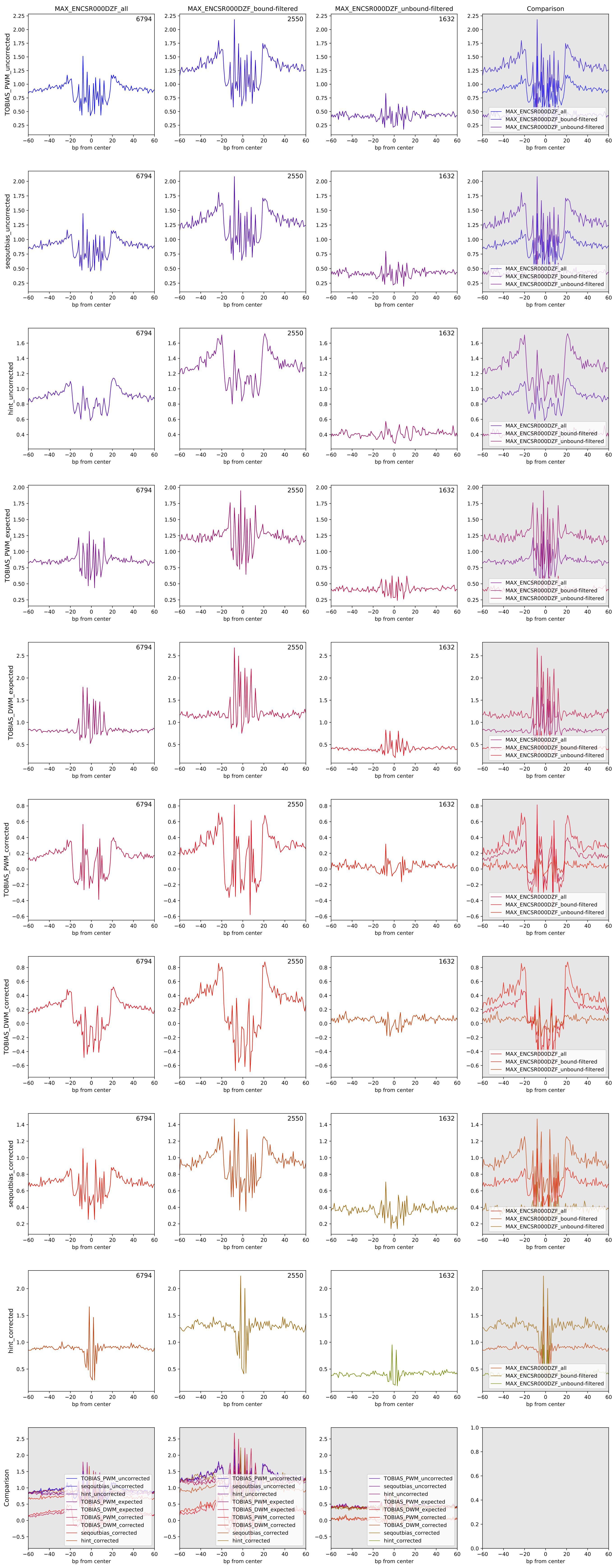

Aggregate footprints for TF MEF2A (ENCSR000BKB)

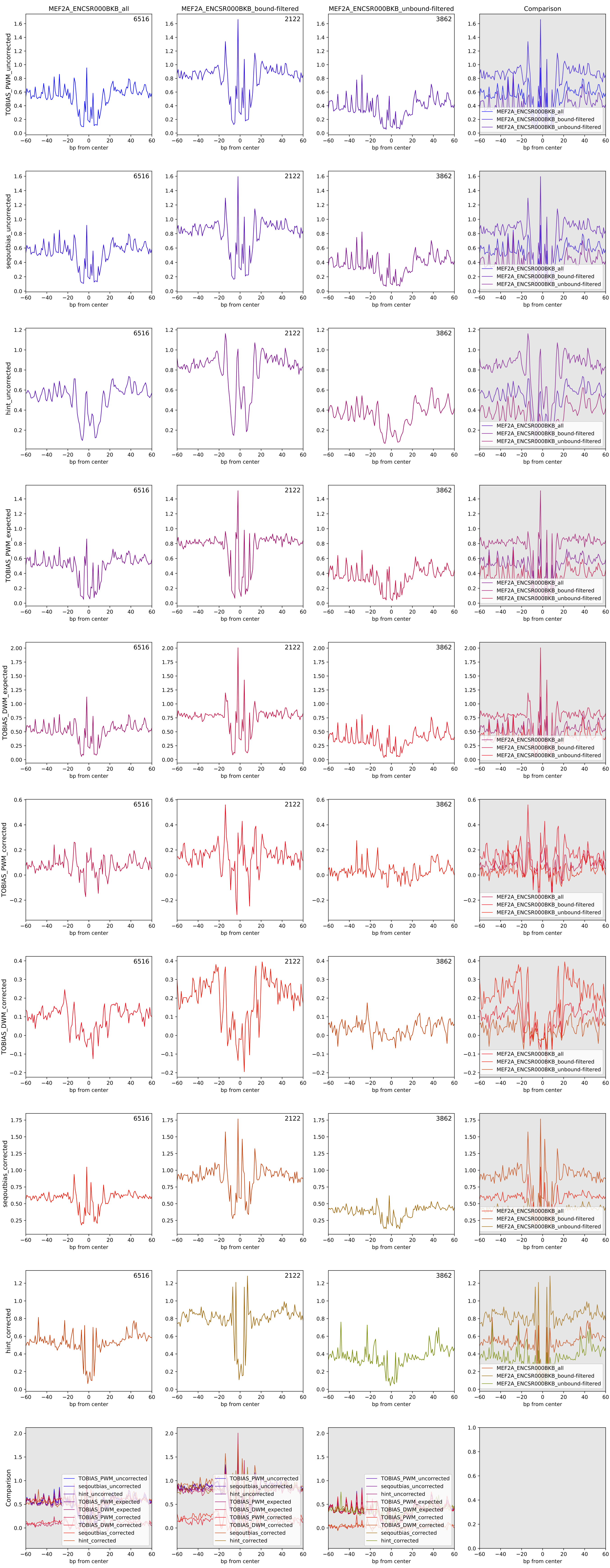

Aggregate footprints for TF MEF2C (ENCSR000BNG)

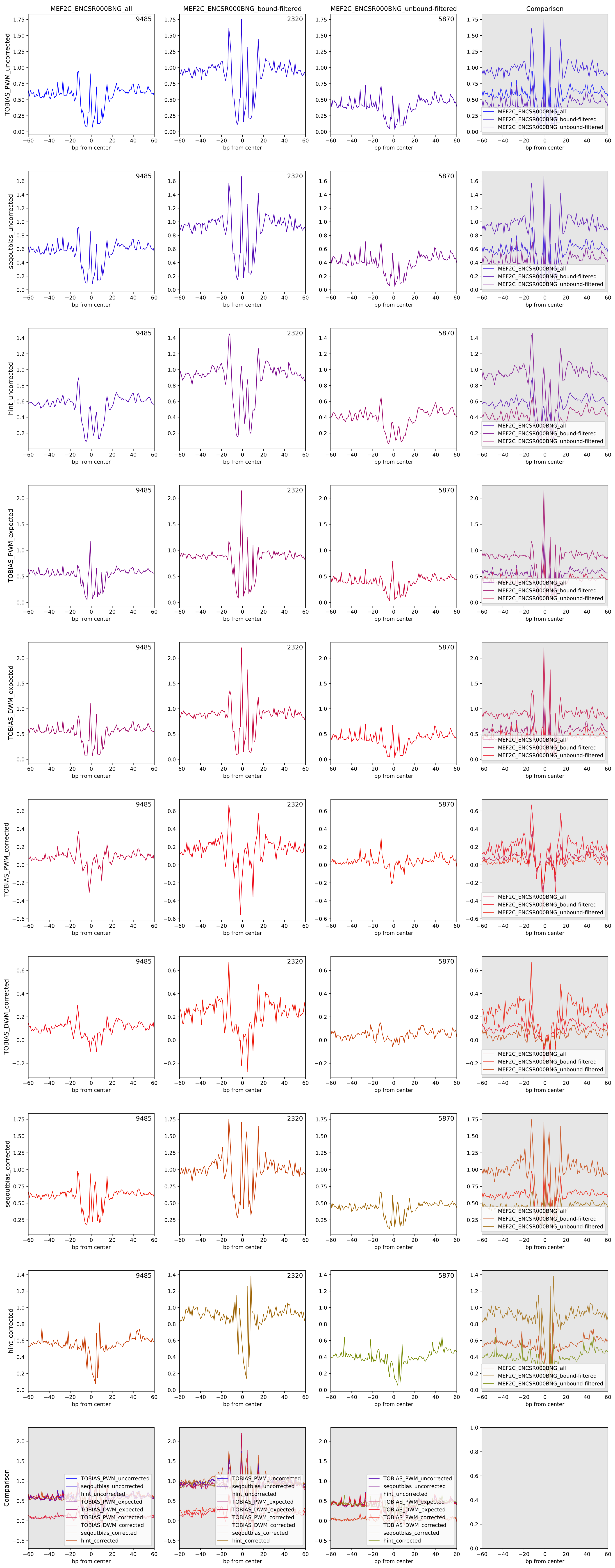

Aggregate footprints for TF MXI1 (ENCSR000DZI)

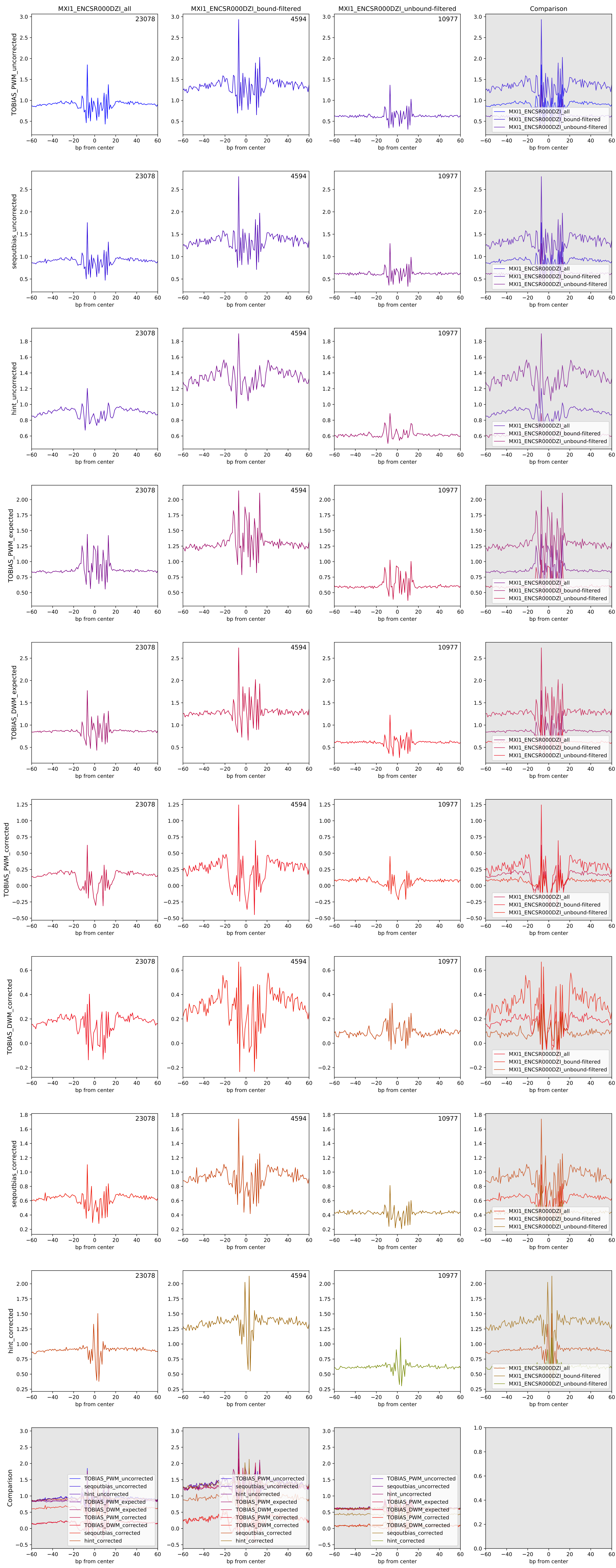

Aggregate footprints for TF NFYB (ENCSR000DNM)

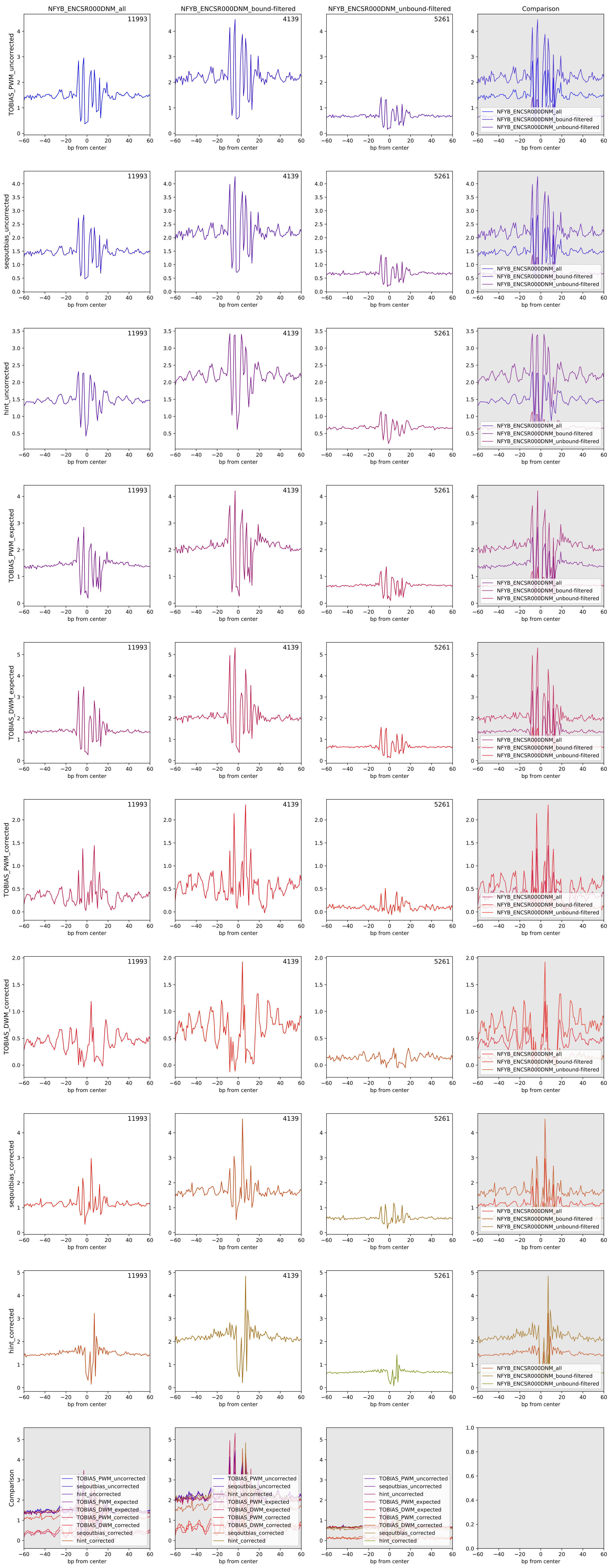

Aggregate footprints for TF NR2F1 (ENCSR514VYD)

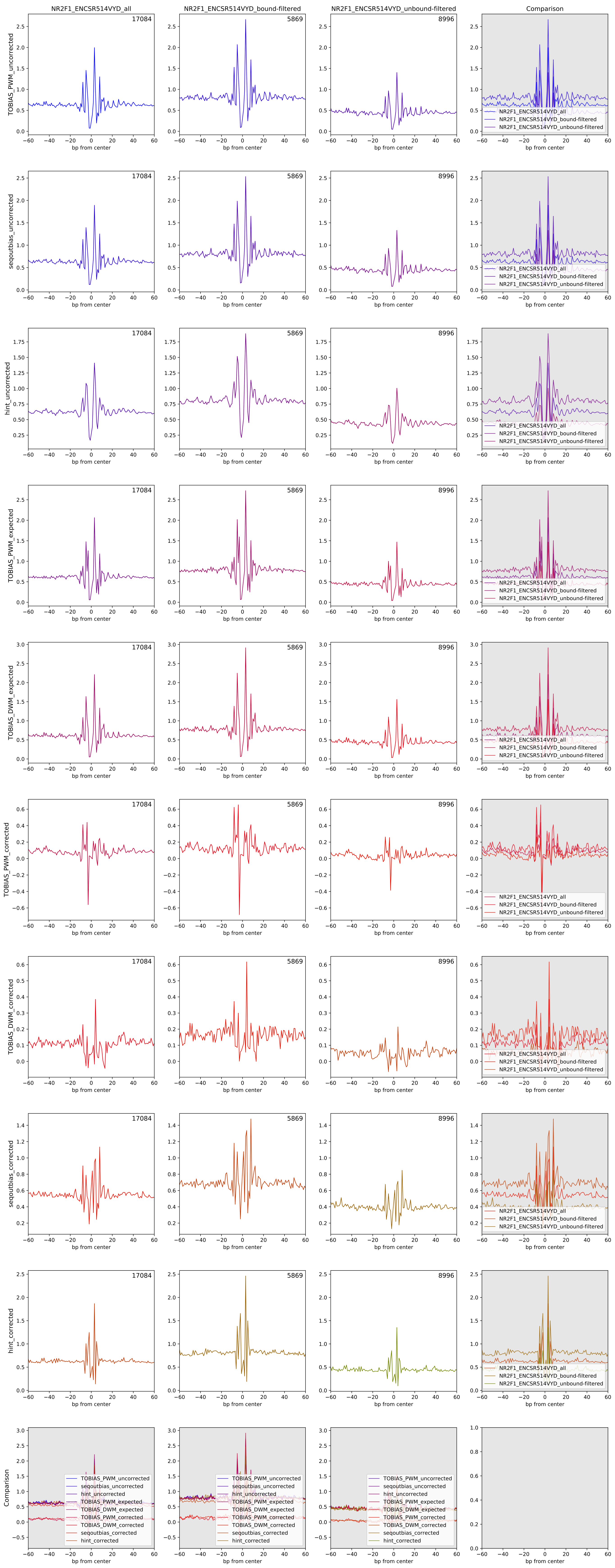

Aggregate footprints for TF NRF1 (ENCSR000DZO)

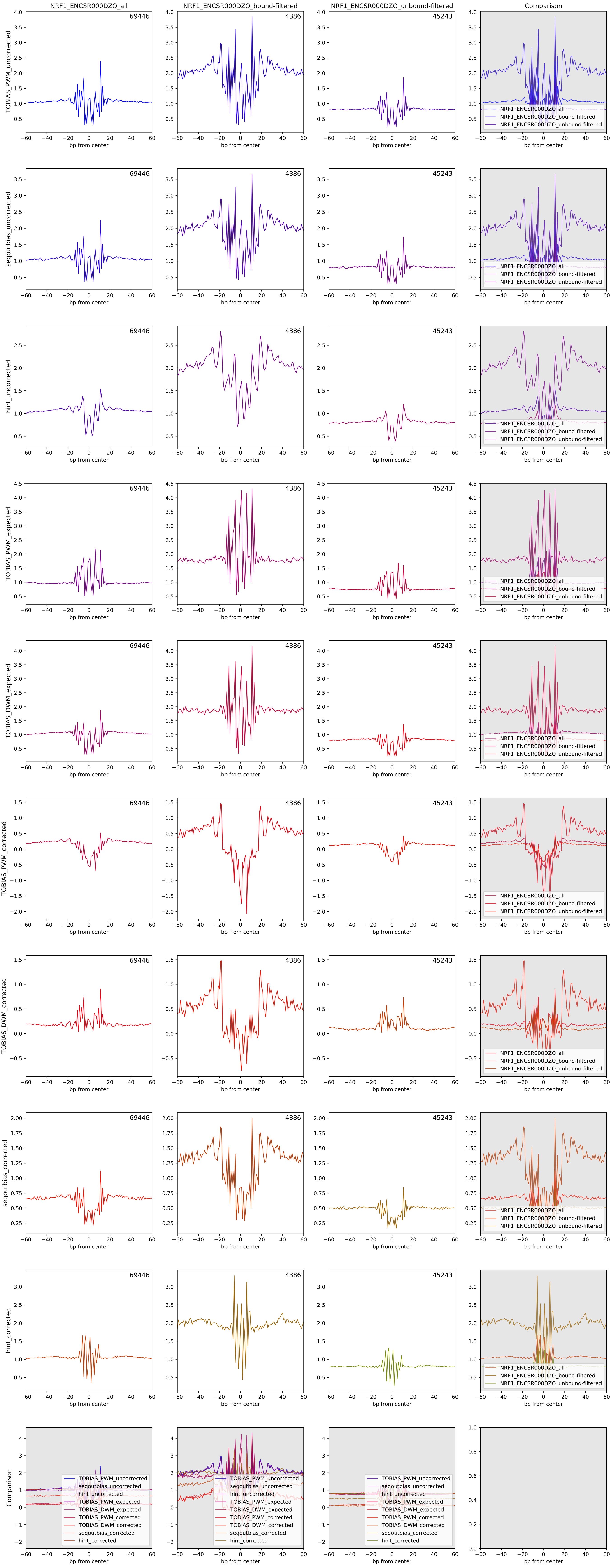

Aggregate footprints for TF PAX5 (ENCSR000BHD)

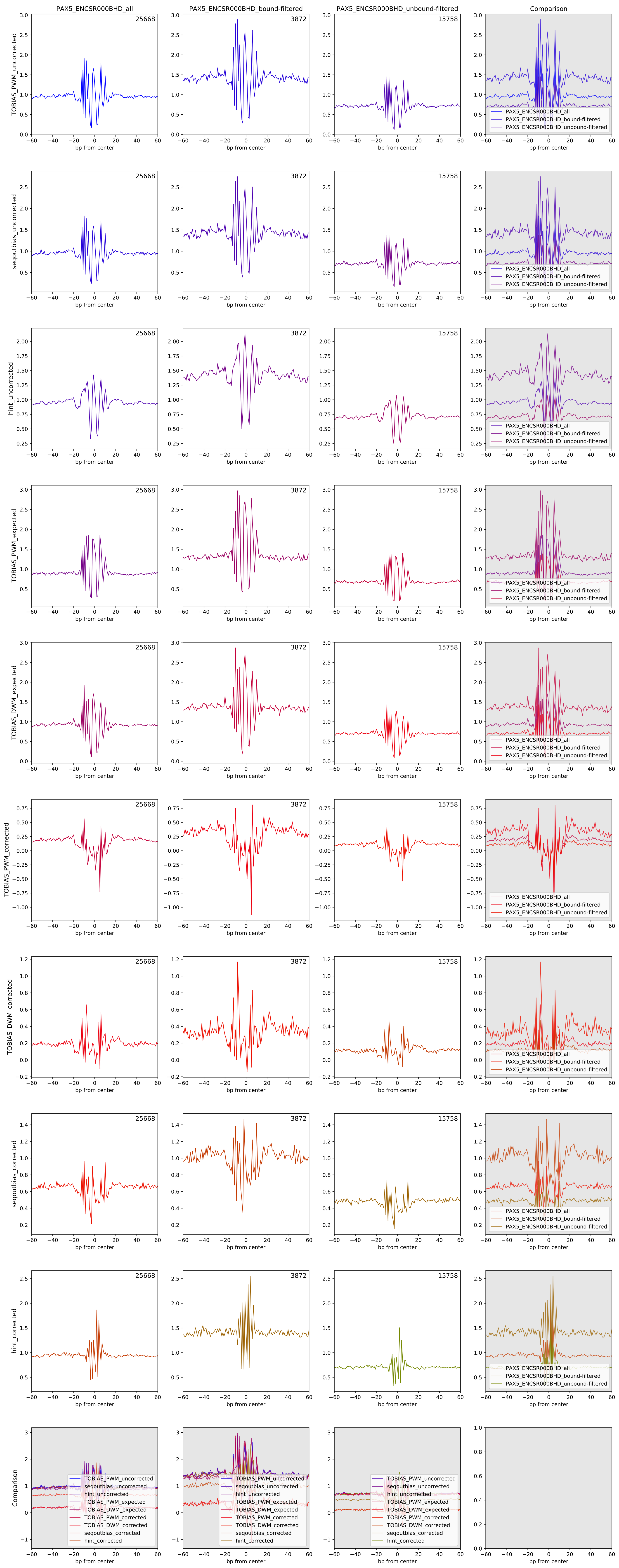

Aggregate footprints for TF PBX3 (ENCSR000BGR)

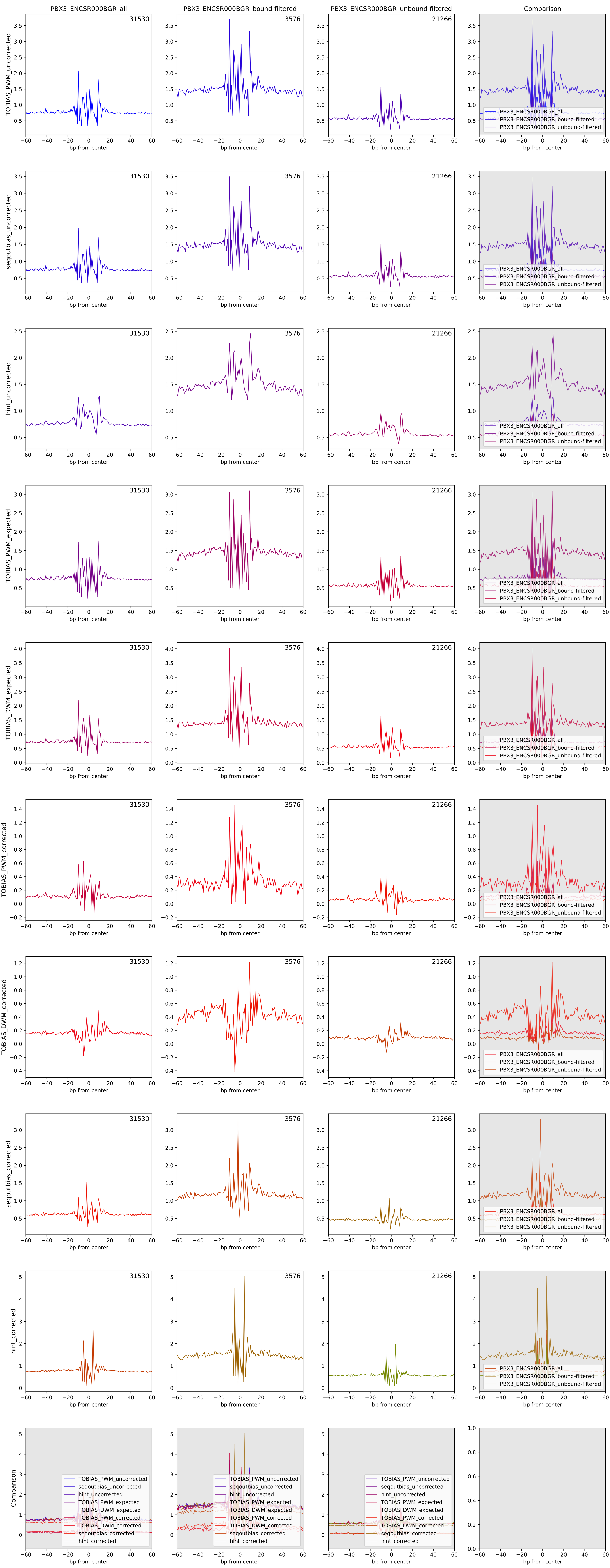

Aggregate footprints for TF PKNOX1 (ENCSR711XNY)

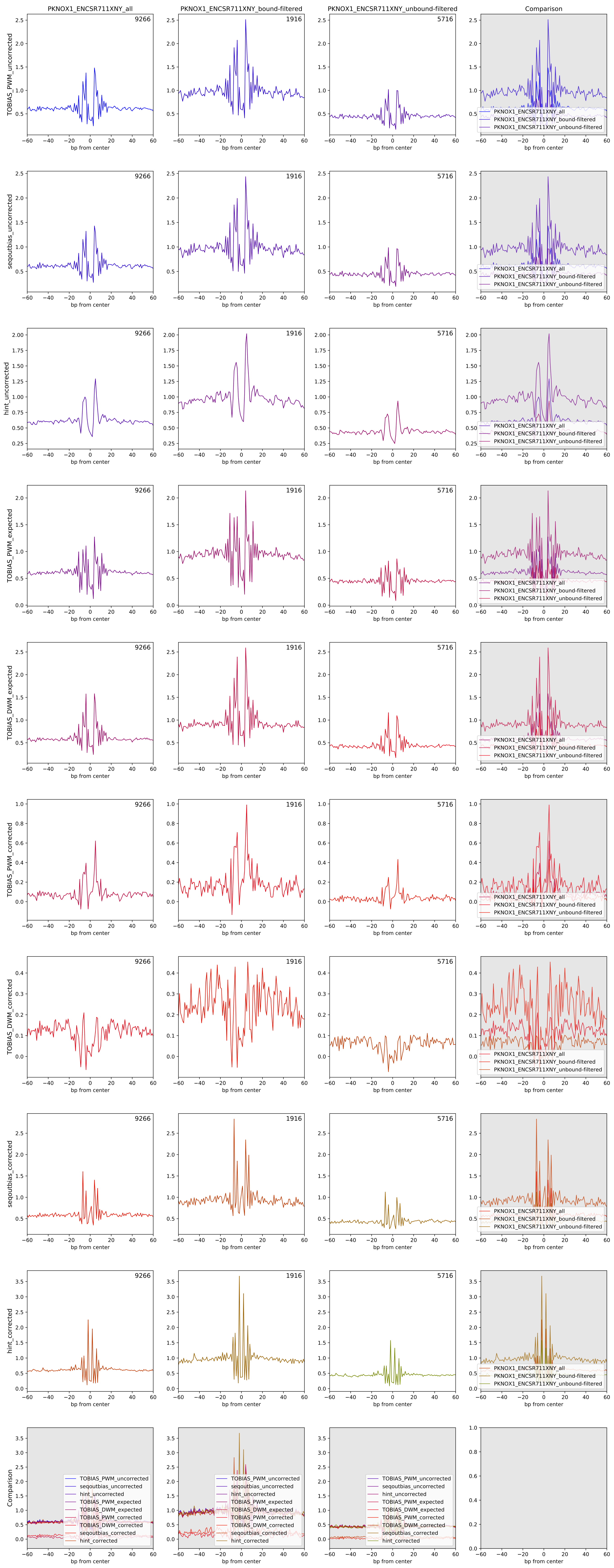

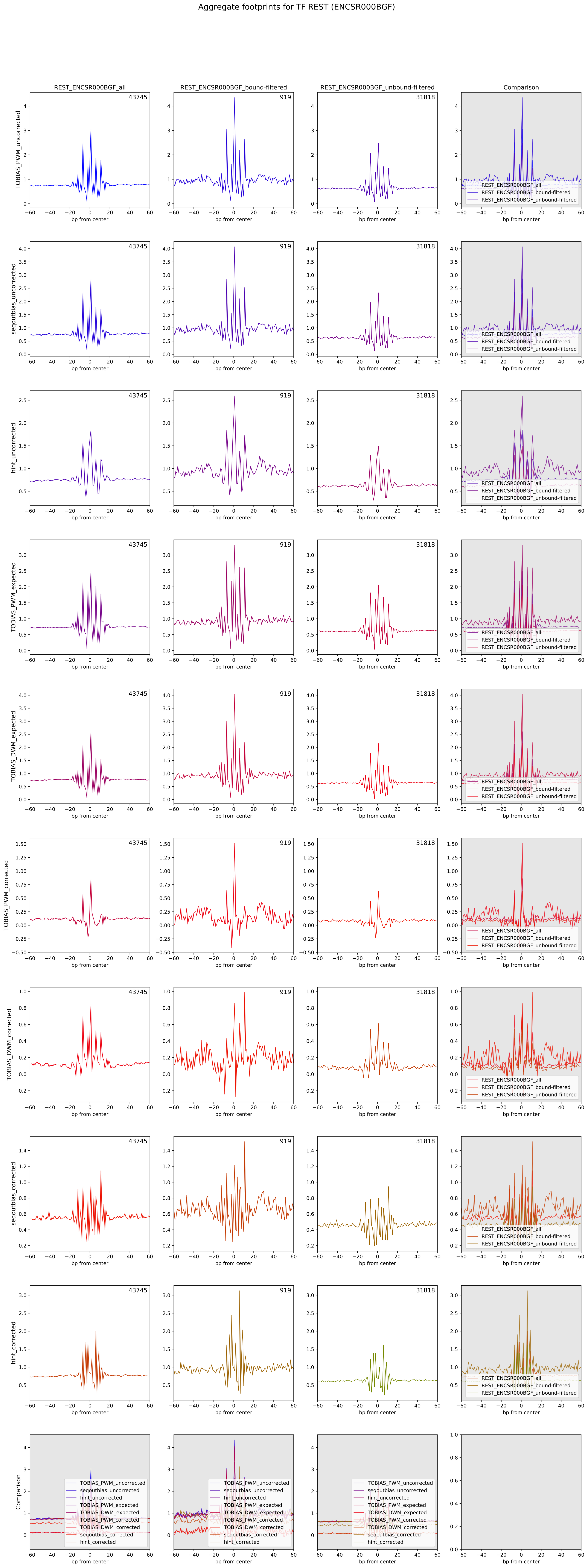

Aggregate footprints for TF REST (ENCSR000BGF)
REST_ENCSR000BGF_all
REST_ENCSR000BGF_bound-filtered
REST_ENCSR000BGF_unbound-filtered
Comparison
43745
919
31818
TOBIAS_PWM_uncorrected
seqoutbias_uncorrected
hint_uncorrected
TOBIAS_PWM_expected
TOBIAS_DWM_expected
TOBIAS_PWM_corrected
TOBIAS_DWM_corrected
seqoutbias_corrected
hint_corrected
Comparison
bp from center
REST_ENCSR000BGF_all
REST_ENCSR000BGF_bound-filtered
REST_ENCSR000BGF_unbound-filtered

Aggregate footprints for TF RUNX3 (ENCSR000BRI)

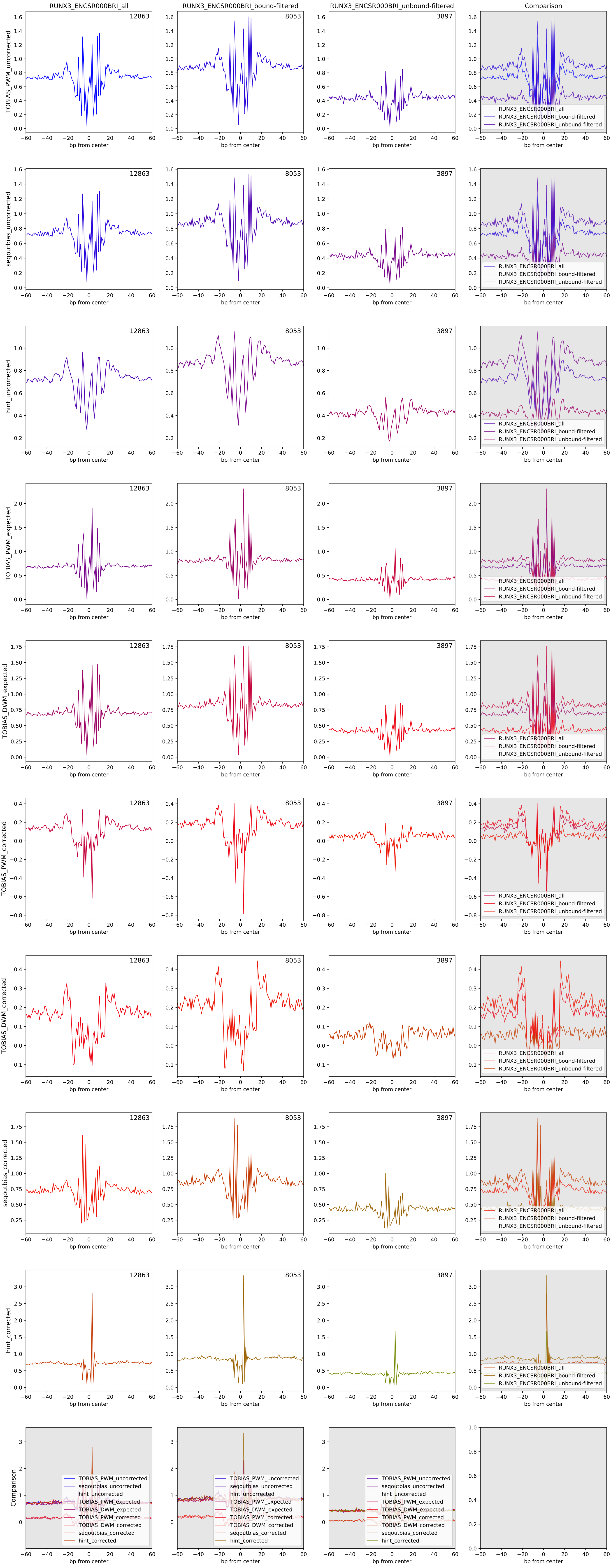

Aggregate footprints for TF SRF (ENCSR041XML)

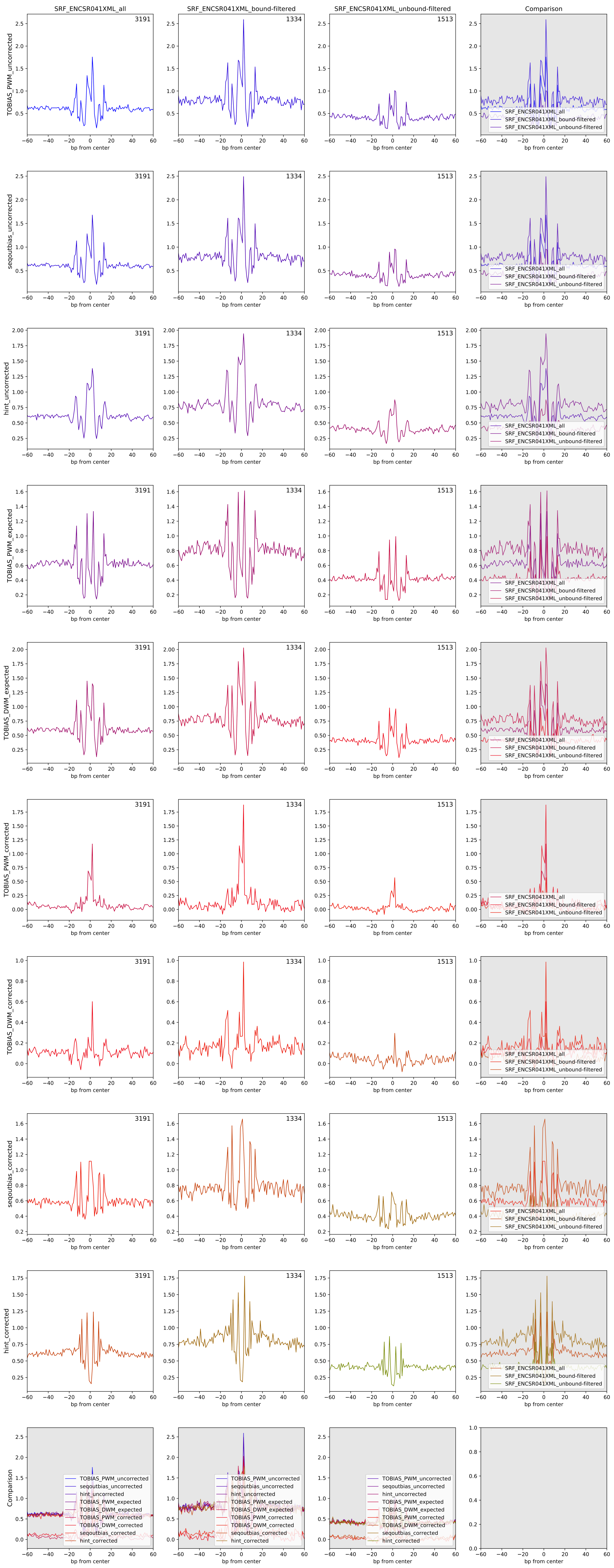

Aggregate footprints for TF TBX21 (ENCSR739IHN)

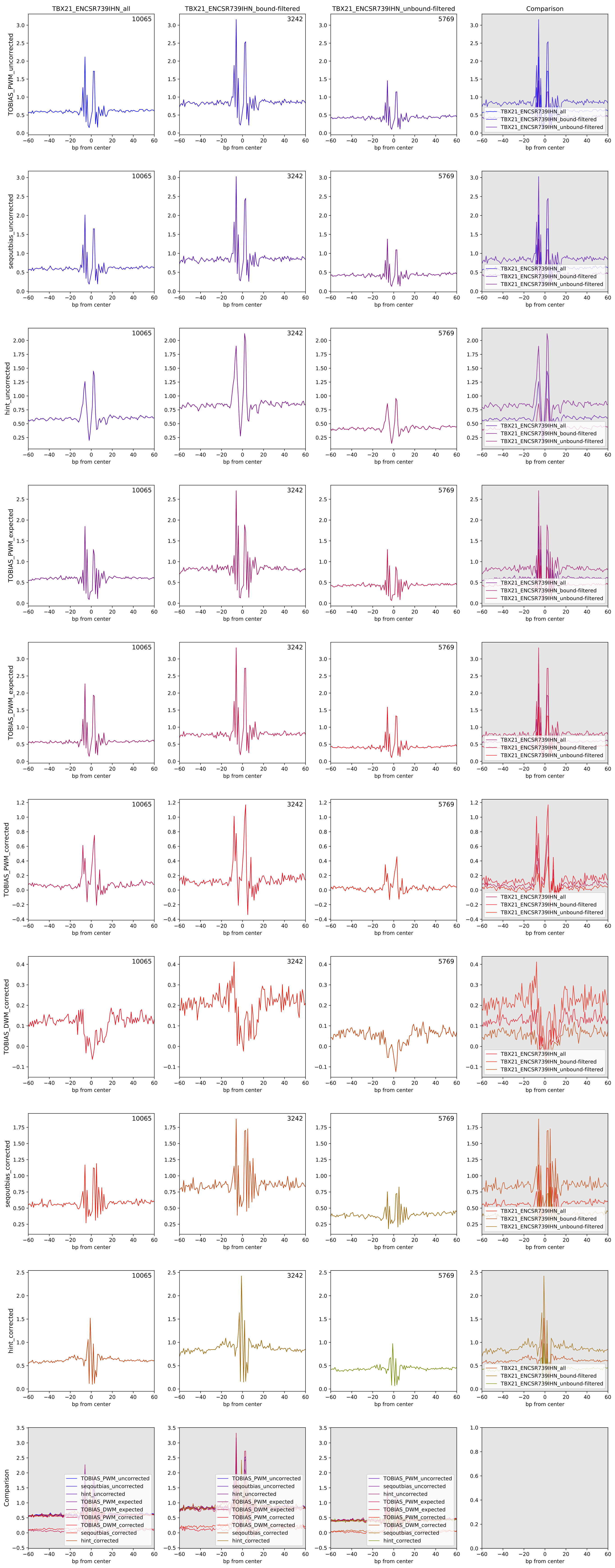

Aggregate footprints for TF TCF12 (ENCSR000BGZ)

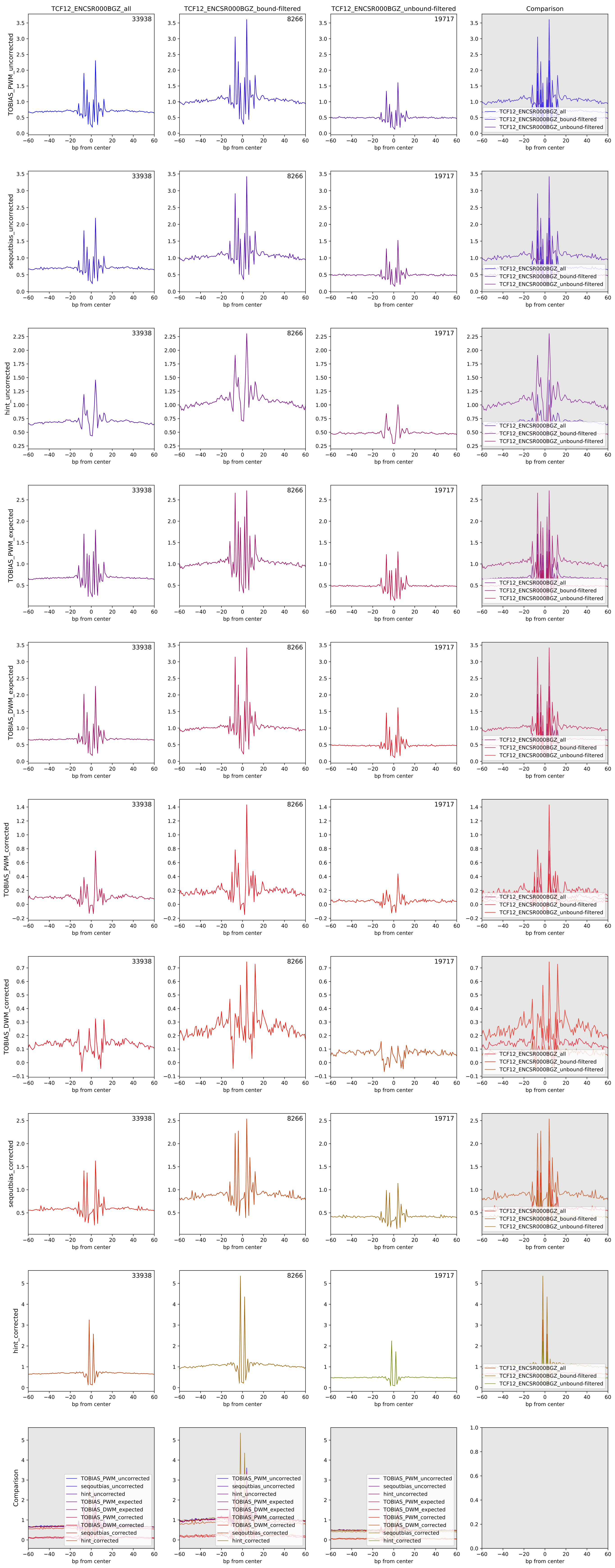

Aggregate footprints for TF TCF7 (ENCSR501DKS)

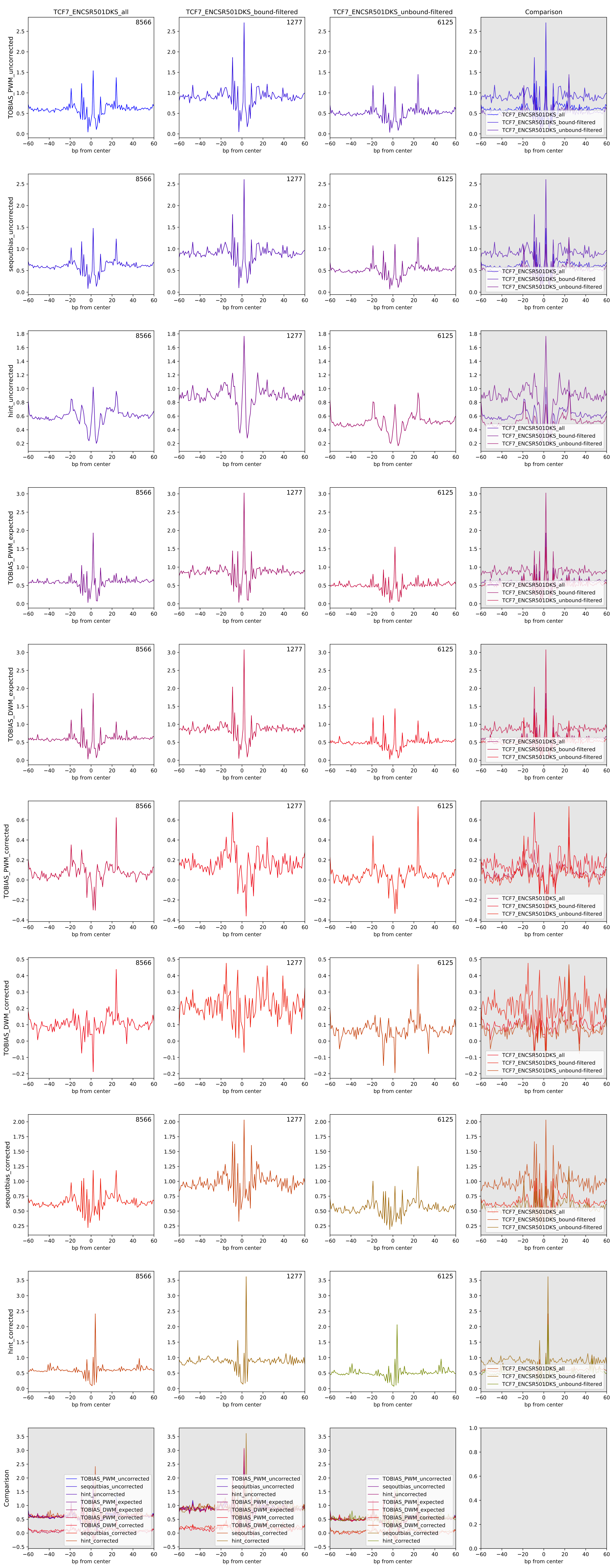

Aggregate footprints for TF USF1 (ENCSR000BGI)

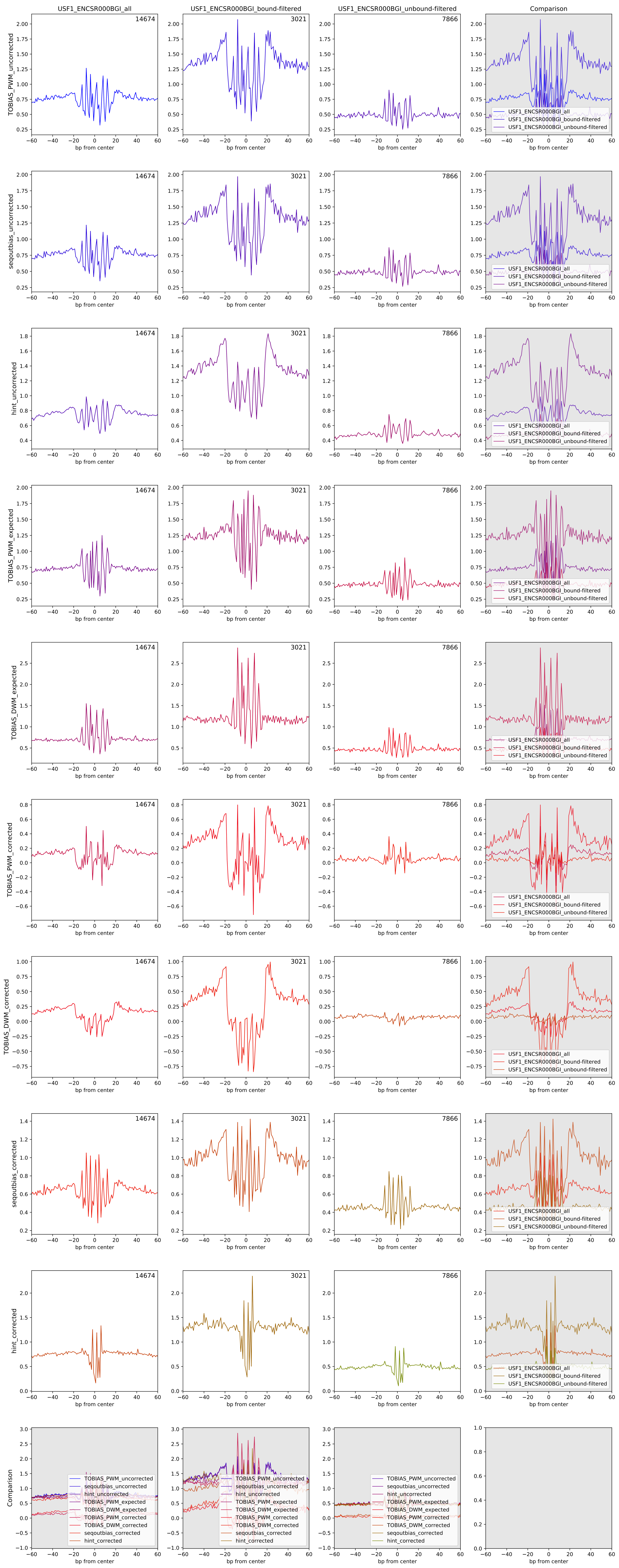

Aggregate footprints for TF USF2 (ENCSR000DZU)

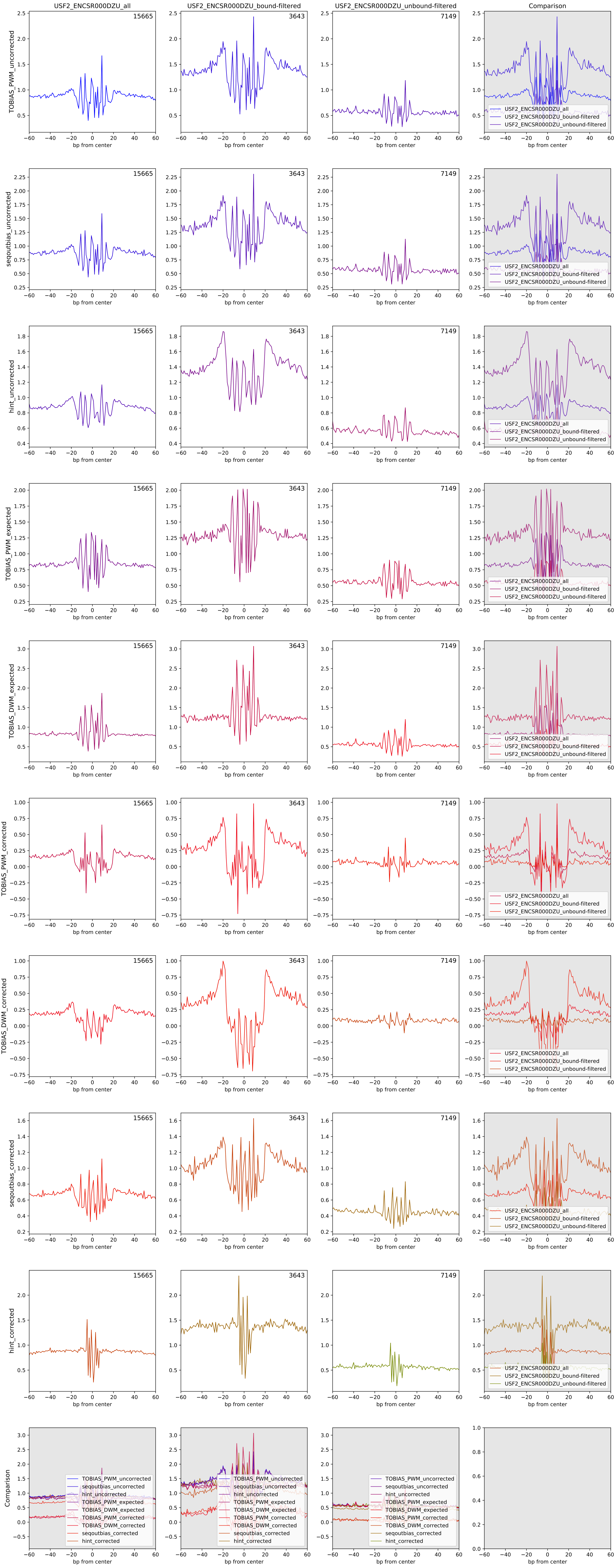

#### Aggregate footprints for TF YY1 (ENCSR000BNP)

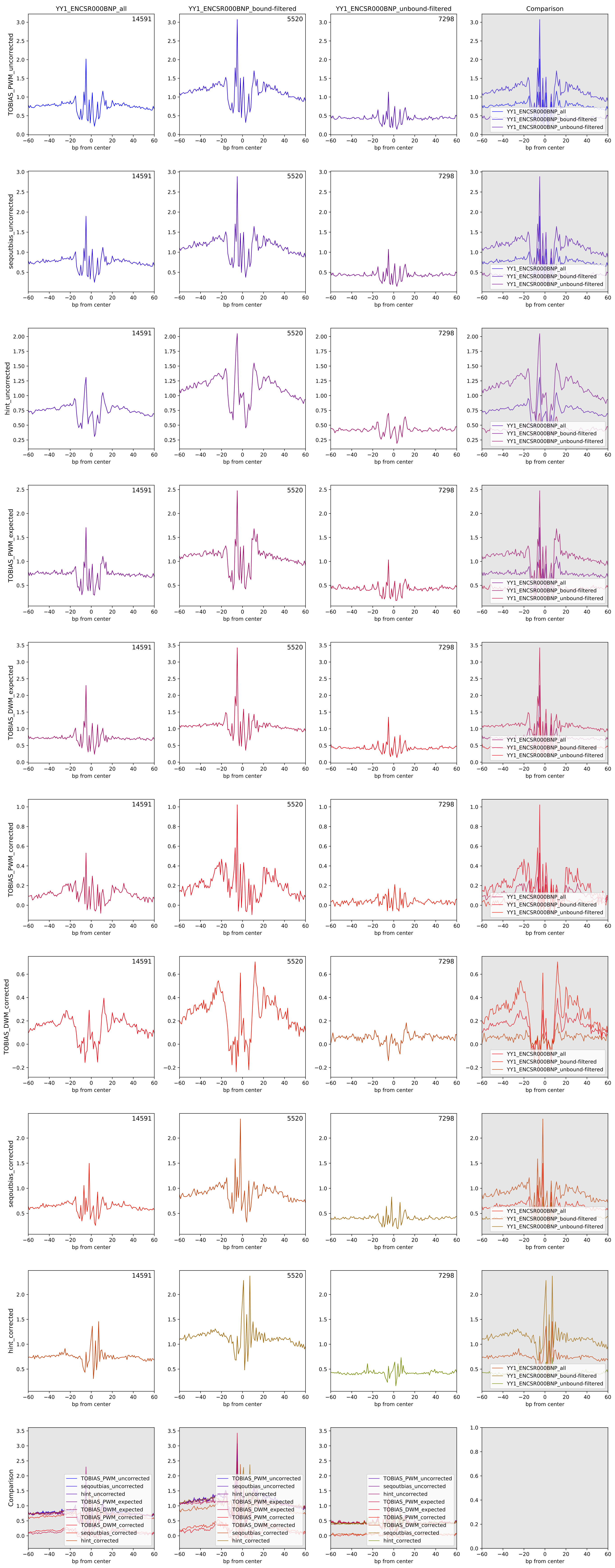

Aggregate footprints for TF ZBTB33 (ENCSR000BHC)

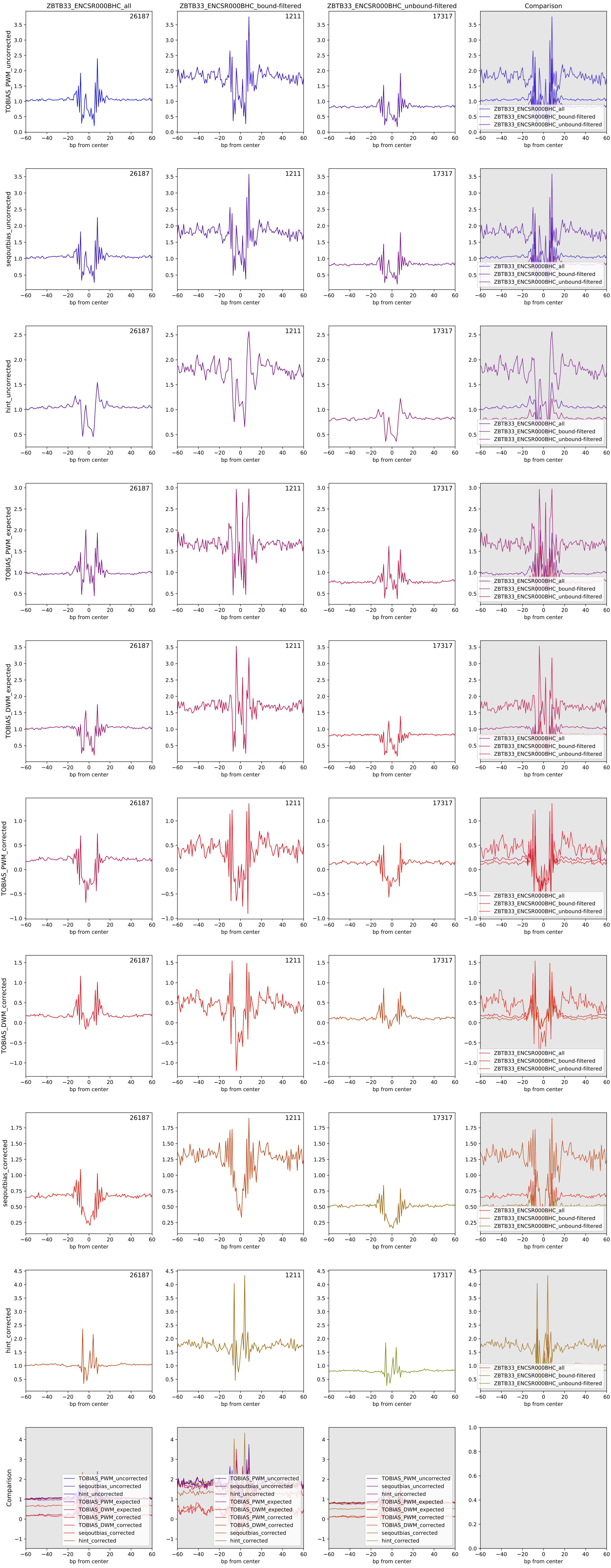

Aggregate footprints for TF ZNF143 (ENCSR000DZL)

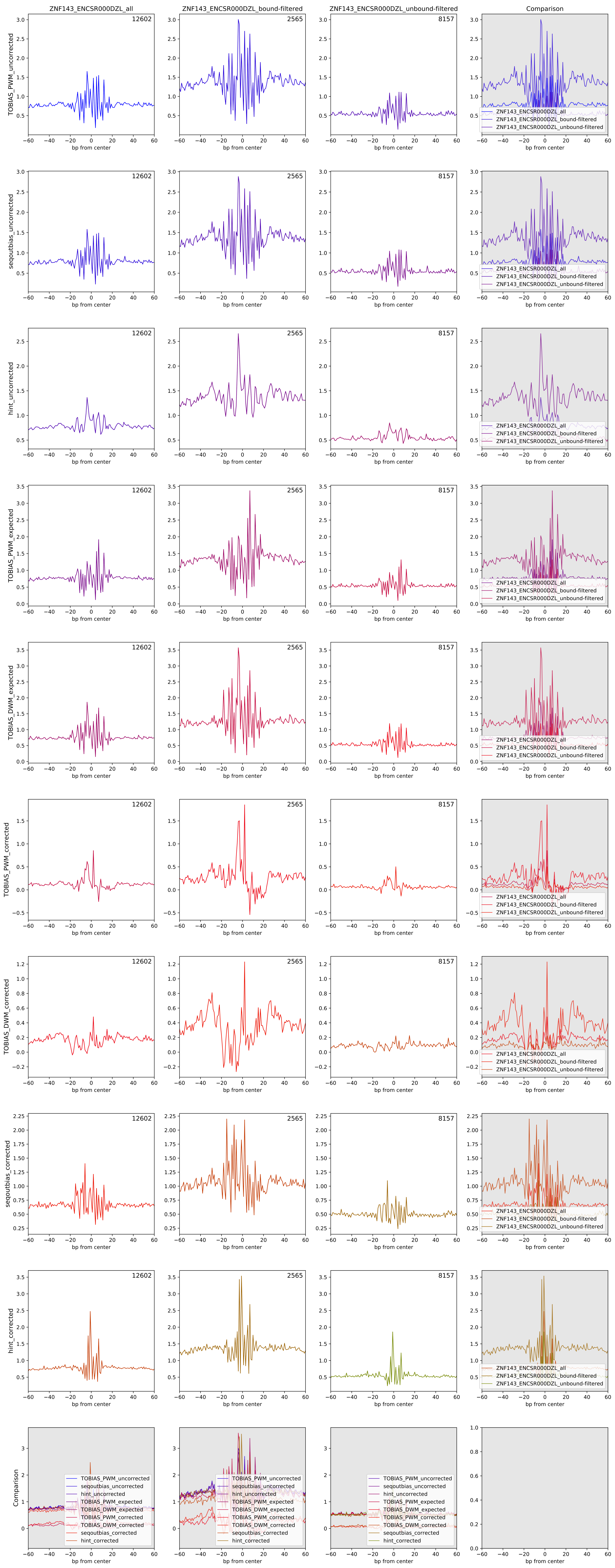

Aggregate footprints for TF ZNF24 (ENCSR072PWP)

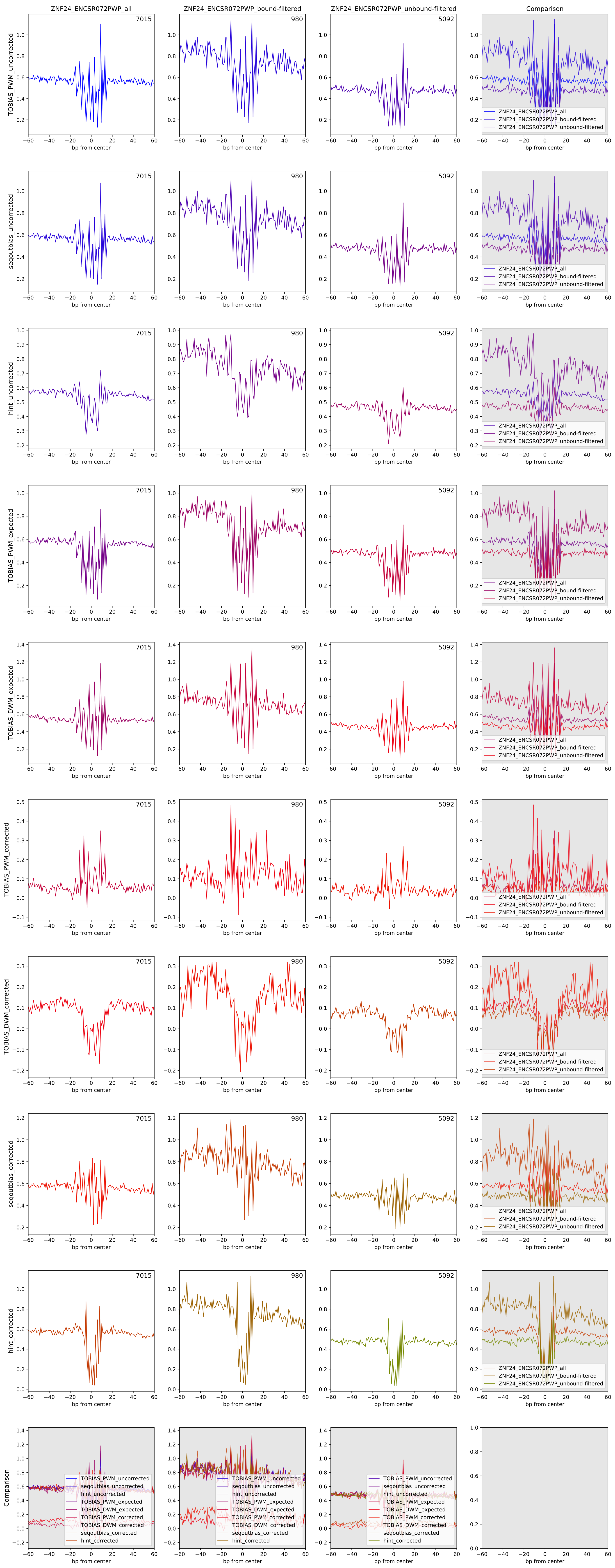

Aggregate footprints for TF ZNF384 (ENCSR000DYP)

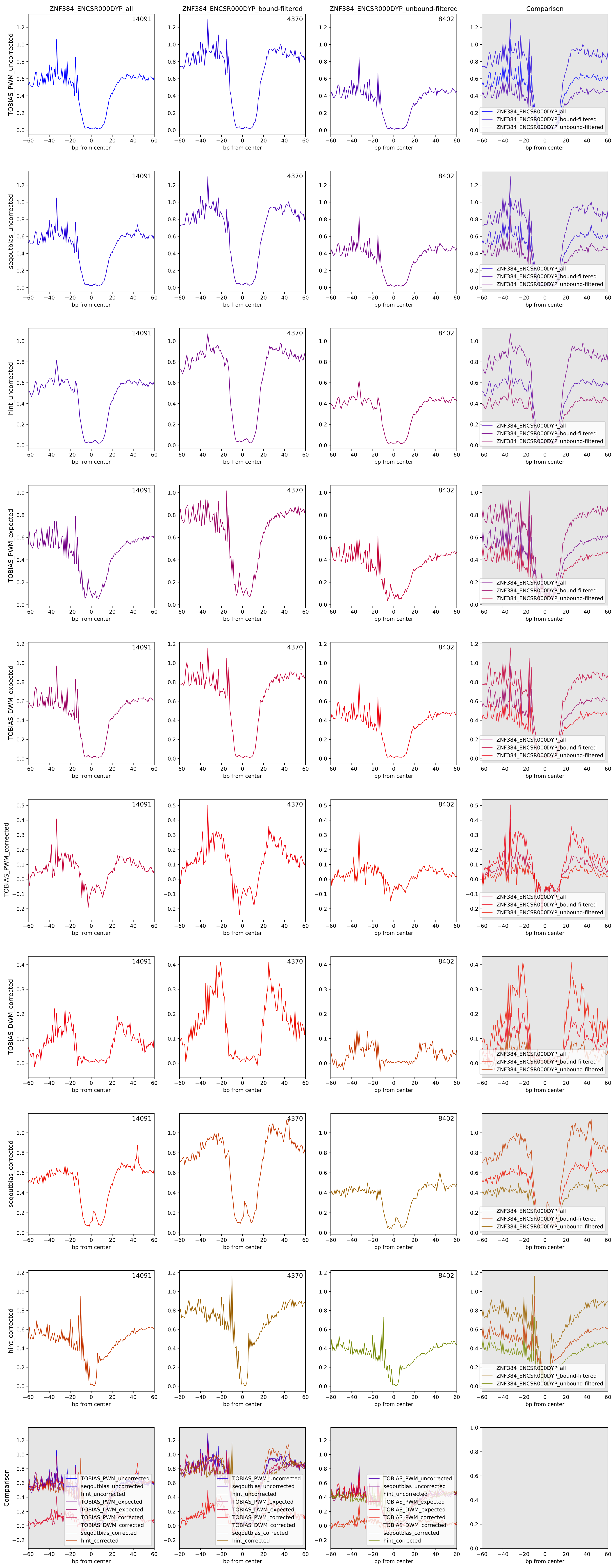
