## Supplemental File 2 for "Beyond accessibility: ATAC-seq footprinting unravels kinetics of transcription factor binding during zygotic genome activation"

### BINDETECT FIGURES

Page 2) Raw score distributions
Page 3) Normalized score distributions
Page 4) BINDetect plot (2C / 4C3PN)
Page 5) BINDetect plot (2C / 8C)
Page 6) BINDetect plot (2C / ICM)
Page 7) BINDetect plot (2C / naive_hESC)
Page 8) BINDetect plot (2C / hESC)
Page 9) BINDetect plot (4C3PN / 8C)
Page 10) BINDetect plot (4C3PN / ICM)
Page 11) BINDetect plot (4C3PN / naive_hESC)
Page 12) BINDetect plot (4C3PN / hESC)
Page 13) BINDetect plot (8C / ICM)
Page 14) BINDetect plot (8C / naive_hESC)
Page 15) BINDetect plot (8C / hESC)
Page 16) BINDetect plot (ICM / naive_hESC)
Page 17) BINDetect plot (ICM / hESC)
Page 18) BINDetect plot (naive_hESC / hESC)

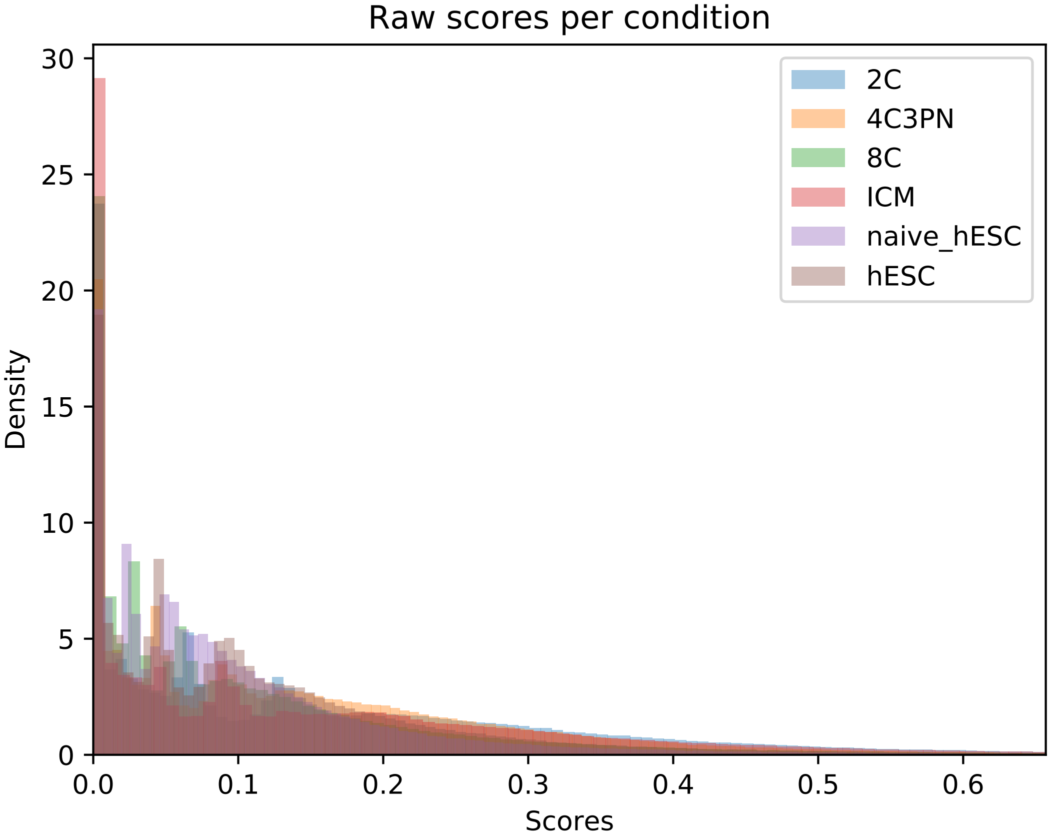

Raw scores per condition
2C
4C3PN
8C
ICM
naive_hESC
hESC
Density
Scores
0
5
10
15
20
25
30
0.0
0.1
0.2
0.3
0.4
0.5
0.6

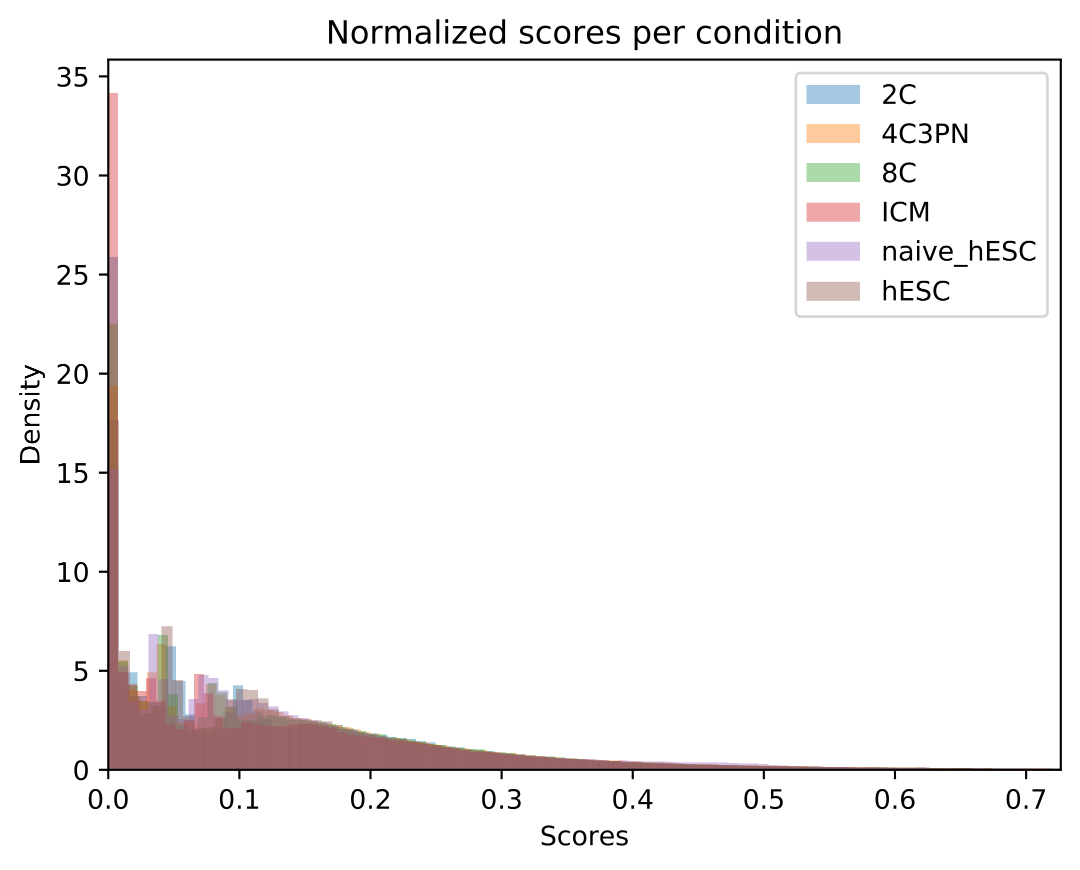

Normalized scores per condition
Density
Scores
2C
4C3PN
8C
ICM
naive_hESC
hESC
0
5
10
15
20
25
30
35
0.0
0.1
0.2
0.3
0.4
0.5
0.6
0.7

BINDetect volcano plot

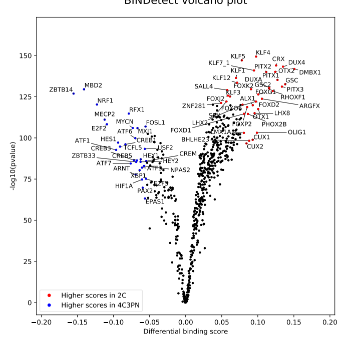


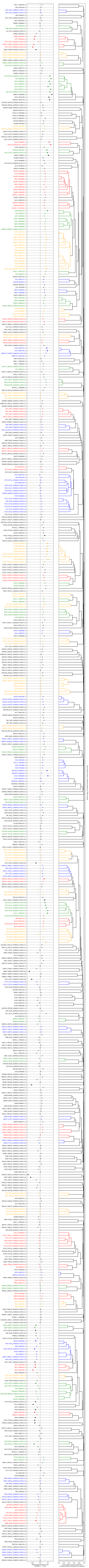

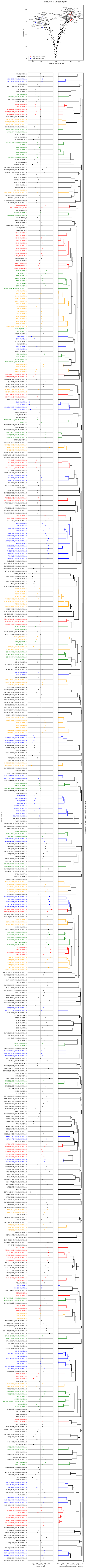
BINDetect volcano plot
ZNF467
VEZF1
PATZ1
KLF6
WT1
KLF1
SP4
SP3
ZNF281
MAZ
SALL4
EGR1
KLF3
FLI1
KLF9
FOXJ3
ZFP105_1
ZNF341
KLF5
ZIC3
ZNF740
ZNF354A
ZNF436
ZBTB17
TCF4
MEF2D
SRY
ETV5
EGR2
SP1
ETV4
ZNF331
MEF2A
AR
PTF1A
ERG
EBF1
KLF16
TLX2
FOXD3
PAX5
GATA6
INSM1
SREBF2
ONECUT2
GATA3
HNF1A
ASCL1
NKX2-5
ETV1
RARB
ESRRG
CUX2
RHOXF1
ETS1
TCF3
ZBTB14
GATA5
ELF2
PBX3
CUX1
ARID5A
ZFP187_1
HLX
TCFL5
BHLHE23
HOMEZ
YY2
HES1
IRF6_1
-log10(pvalue)
Differential binding score
Higher scores in 2C
Higher scores in 8C
SIX6_1 (PB0059.1)
SIX4 (PH0164.1)
SIX2 (SIX2_HUMAN.H11MO.0.A)
SIX1 (SIX1_HUMAN.H11MO.0.A)
HDX (PH0037.1)
GFI1 (GFI1_HUMAN.H11MO.0.C)
NFIL3 (MA0025.1)
GMEB2 (MA0862.1)
TEF (MA0843.1)
DBP (DBP_HUMAN.H11MO.0.B)
HLF (HLF_HUMAN.H11MO.0.C)
GMEB1 (MA0615.1)
CREB5 (MA0840.1)
ATF7 (MA0834.1)
ATF2 (ATF2_HUMAN.H11MO.0.B)
CREB1 (CREB1_HUMAN.H11MO.0.A)
ATF1 (ATF1_HUMAN.H11MO.0.B)
CREM (CREM_HUMAN.H11MO.0.C)
CEBPB (CEBPB_HUMAN.H11MO.0.A)
CEBPA (CEBPA_HUMAN.H11MO.0.A)
CEBPD (CEBPD_HUMAN.H11MO.0.C)
CEBPE (CEBPE_HUMAN.H11MO.0.A)
CEBPG (CEBPG_HUMAN.H11MO.0.B)
ATF4 (ATF4_HUMAN.H11MO.0.A)
MYBL2 (MA0777.1)
BARHL2 (MA0635.1)

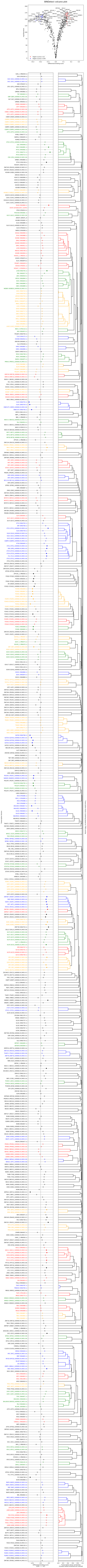

BINDetect volcano plot
-log10(pvalue)
Differential binding score
Higher scores in 2C
Higher scores in ICM
SIX6_1 (PB0059.1)
SIX4 (PH0164.1)
SIX2 (SIX2_HUMAN.H11MO.0.A)
SIX1 (SIX1_HUMAN.H11MO.0.A)
HDX (PH0037.1)
GFI1 (GFI1_HUMAN.H11MO.0.C)
NFIL3 (MA0025.1)
GMEB2 (MA0862.1)
TEF (MA0843.1)
DBP (DBP_HUMAN.H11MO.0.B)
HLF (HLF_HUMAN.H11MO.0.C)
GMEB1 (MA0615.1)
CREB5 (MA0840.1)
ATF7 (MA0834.1)
ATF2 (ATF2_HUMAN.H11MO.0.B)
CREB1 (CREB1_HUMAN.H11MO.0.A)
ATF1 (ATF1_HUMAN.H11MO.0.B)
CREM (CREM_HUMAN.H11MO.0.C)
CEBPB (CEBPB_HUMAN.H11MO.0.A)
CEBPA (CEBPA_HUMAN.H11MO.0.A)
CEBPD (CEBPD_HUMAN.H11MO.0.C)
CEBPE (CEBPE_HUMAN.H11MO.0.A)
CEBPG (CEBPG_HUMAN.H11MO.0.B)
ATF4 (ATF4_HUMAN.H11MO.0.A)
MYBL2 (MA0777.1)
BARHL2 (MA0635.1)
HOMEZ (PH0044.1)
OTX2 (OTX2_HUMAN.H11MO.0.A)
GSC (MA0648.1)
GSC2 (MA0891.1)
PITX1 (MA0682.1)
OTX1 (MA0711.1)
PITX3 (MA0714.1)
CRX (CRX_HUMAN.H11MO.0.B)
DMBX1 (MA0883.1)
PITX2 (PH0138.1)
RHOXF1 (MA0719.1)
ZNF329 (ZN329_HUMAN.H11MO.0.C)

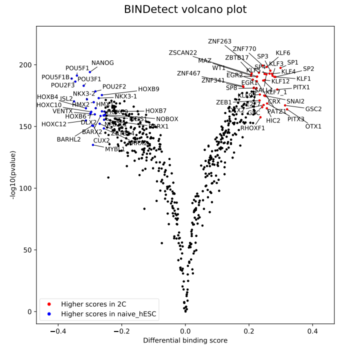
BINDetect volcano plot
-log10(pvalue)
Differential binding score
Higher scores in 2C
Higher scores in naive_hESC
NANOG
POU5F1
POU5F1B
POU3F1
POU2F3
POU2F2
HOXB9
NKX3-2
NKX3-1
HOXB4
ISL2
HOXC10
HMX2
HMX1
VENTX
HOXB7
HOXB6
NOBOX
HOXC12
DLX2
BARX1
BARX2
BARHL2
CUX2
MYBL1
ZNF263
ZNF770
ZSCAN22
ZBTB17
SP3
KLF6
MAZ
KLF3
SP1
WT1
SP4
KLF5
SP2
KLF4
ZNF467
EGR2
KLF1
ZNF341
EGR1
KLF12
SP8
PITX1
SALL4
KLF7_1
KLF9
ZEB1
CRX
SNAI2
E2F7
GSC2
DLX2
GSC
PATZ1
HIC2
PITX3
OTX1
RHOXF1
200
150
100
50
0
−0.4
−0.2
0.0
0.2
0.4

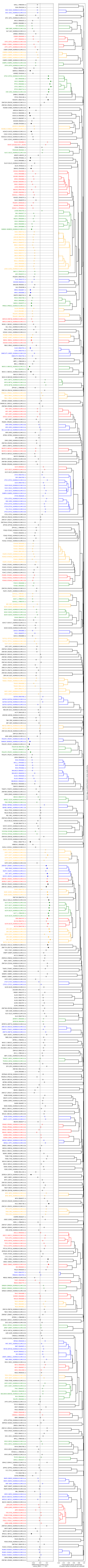
SIX6_1 (PB0059.1)
SIX4 (PH0164.1)
SIX2 (SIX2_HUMAN.H11MO.0.A)
SIX1 (SIX1_HUMAN.H11MO.0.A)
HDX (PH0037.1)
GFI1 (GFI1_HUMAN.H11MO.0.C)
NFIL3 (MA0025.1)
GMEB2 (MA0862.1)
TEF (MA0843.1)
DBP (DBP_HUMAN.H11MO.0.B)
HLF (HLF_HUMAN.H11MO.0.C)
GMEB1 (MA0615.1)
CREB5 (MA0840.1)
ATF7 (MA0834.1)
ATF2 (ATF2_HUMAN.H11MO.0.B)
CREB1 (CREB1_HUMAN.H11MO.0.A)
ATF1 (ATF1_HUMAN.H11MO.0.B)
CREM (CREM_HUMAN.H11MO.0.C)
CEBPB (CEBPB_HUMAN.H11MO.0.A)
CEBPA (CEBPA_HUMAN.H11MO.0.A)
CEBPD (CEBPD_HUMAN.H11MO.0.C)
CEBPE (CEBPE_HUMAN.H11MO.0.A)
CEBPG (CEBPG_HUMAN.H11MO.0.B)
ATF4 (ATF4_HUMAN.H11MO.0.A)
MYBL2 (MA0777.1)
BARHL2 (MA0635.1)
HOMEZ (PH0044.1)
OTX2 (OTX2_HUMAN.H11MO.0.A)
GSC (MA0648.1)
GSC2 (MA0891.1)
PITX1 (MA0682.1)
CREB3 (MA0638.1)
CREB3L1 (MA0839.1)
XBP1 (MA0844.1)
ATF6 (ATF6A_HUMAN.H11MO.0.B)
ZBTB33 (KAISO_HUMAN.H11MO.0.A)
ZBED1 (MA0749.1)
ZNF121 (ZN121_HUMAN.H11MO.0.C)
PAX5 (PAX5_HUMAN.H11MO.0.A)
SPZ1 (MA0111.1)
HIC1 (HIC1_HUMAN.H11MO.0.C)
IRF6_1 (PB0036.1)
RFX2 (RFX2_HUMAN.H11MO.0.A)
RFX1 (RFX1_HUMAN.H11MO.0.B)
RFX4 (MA0799.1)
RFX3 (RFX3_HUMAN.H11MO.0.B)
RFXDC2_1 (PB0056.1)
RFX5 (RFX5_HUMAN.H11MO.0.A)
TFCP2 (MA0145.3)
GRHL1 (MA0647.1)
GRHL2 (GRHL2_HUMAN.H11MO.0.A)
PROX1 (MA0794.1)
ZFP42 (ZFP42_HUMAN.H11MO.0.A)
YY1 (TYY1_HUMAN.H11MO.0.A)
YY2 (MA0748.1)
SOX8 (MA0868.1)
MAFK (MAFK_HUMAN.H11MO.0.A)
MAFG (MAFG_HUMAN.H11MO.0.A)
MAF (MAF_HUMAN.H11MO.0.A)
MAFF (MAFF_HUMAN.H11MO.0.B)
MAFB (MAFB_HUMAN.H11MO.0.B)
NRL (MA0842.1)
NFE2 (NFE2_HUMAN.H11MO.0.A)
BACH2 (BACH2_HUMAN.H11MO.0.A)
NFE2L2 (NF2L2_HUMAN.H11MO.0.A)
BACH1 (BACH1_HUMAN.H11MO.0.A)
JUN (JUN_HUMAN.H11MO.0.A)
JUND (JUND_HUMAN.H11MO.0.A)
JDP2 (MA0655.1)
FOS (FOS_HUMAN.H11MO.0.A)
FOSB (FOSB_HUMAN.H11MO.0.A)
FOSL2 (FOSL2_HUMAN.H11MO.0.A)
FOSL1 (FOSL1_HUMAN.H11MO.0.A)
JUNB (JUNB_HUMAN.H11MO.0.A)
ZNF554 (ZN554_HUMAN.H11MO.0.C)
BATF (BATF_HUMAN.H11MO.0.A)
BATF3 (BATF3_HUMAN.H11MO.0.B)
NFE2L1 (NF2L1_HUMAN.H11MO.0.C)
ZNF549 (ZN549_HUMAN.H11MO.0.C)
PAX9 (MA0781.1)
PAX6 (PAX6_HUMAN.H11MO.0.C)
ZNF354C (MA0130.1)
FOXH1 (FOXH1_HUMAN.H11MO.0.A)
RUNX3 (RUNX3_HUMAN.H11MO.0.A)
RUNX1 (RUNX1_HUMAN.H11MO.0.A)
RUNX2 (RUNX2_HUMAN.H11MO.0.A)
CBFB (PEBB_HUMAN.H11MO.0.C)
ZNF382 (ZN382_HUMAN.H11MO.0.C)
−0.2
0.0
0.2
Differential binding score
(naive_hESC ← → 2C)
0.0
0.2
0.4
0.6
0.8
1.0
Transcription factor similarities
(Clusters below threshold are colored)

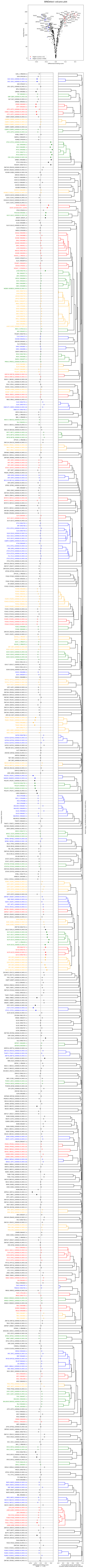
BINDetect volcano plot
-log10(pvalue)
Differential binding score
Higher scores in 2C
Higher scores in hESC

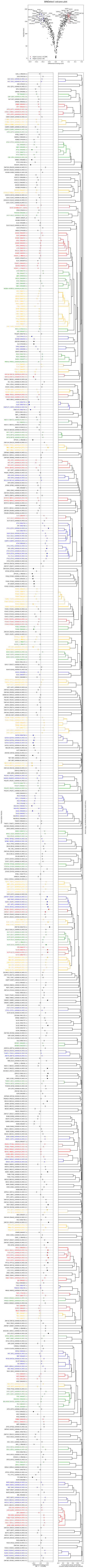
BINDetect volcano plot
-log10(pvalue)
Differential binding score
Higher scores in 4C3PN
Higher scores in 8C
ZFP105_1
MEF2D
MEF2A
FOXJ3
SRY
ZNF354A
FOXD3
GATA3
AR
FOXJ2
RHOXF1
ONECUT2
PITX1
TLX2
FOXB1
DMBX1
FOXG1
GATA5
CUX2
ZFP187
GATA2
GATA6
CUX1
HNF1A
HLX
ARID5A
OLIG1
ARID3B
BHLHE23
YY2
TCFL5
VEZF1
PATZ1
ZNF281
ZNF467
WT1
EGR1
ZBTB17
FLI1
SREBF2
ERG
ZNF331
ZNF740
KLF6
MAZ
ETV4
PTF1A
ZIC3
NKX2-5
KLF16
SP3
TCF3
MYF6
LYL1
ASCL1
KLF9
NFIC
ZNF563
NR1H4
HSM1
REST
PBX3
ETV1
ZIC2_1
GLIS2
ZNF423
Differential binding score
(8C ← → 4C3PN)
Transcription factor similarities
(Clusters below threshold are colored)

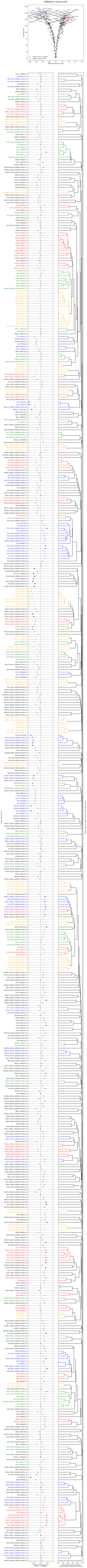
BINDetect volcano plot
-log10(pvalue)
Differential binding score
Higher scores in 4C3PN
Higher scores in ICM

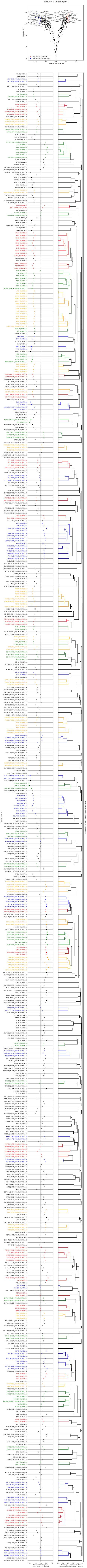

BINDetect volcano plot
-log10(pvalue)
Differential binding score
Higher scores in 4C3PN
Higher scores in naive_hESC
Differential binding score
(naive_hESC ← → 4C3PN)
Transcription factor similarities
(Clusters below threshold are colored)

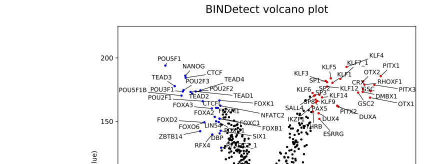
BINDetect volcano plot
-log10(pvalue)
Differential binding score
Higher scores in 4C3PN
Higher scores in hESC
POU5F1
NANOG
CTCF
TEAD3
POU2F3
TEAD4
POU5F1B
POU3F1
POU2F2
TEAD1
TEAD2
POU2F1
FOXK1
FOXA3
CTCFL
FOXM1
NFATC2
FOXA2
FOXD2
FOXC1
FOXO6
LIN54
FOXB1
ZBTB14
SIX1
DBP
RFX4
GMEB2
KLF4
KLF7_1
PITX1
KLF5
KLF3
KLF1
OTX2
SP1
CRX
RHOXF1
KLF6
SP2
KLF12
GSC
PITX3
SP3
DMBX1
KLF14
SP8
KLF9
GSC2
OTX1
SALL4
PAX5
PITX2
DUXA
IKZF1
DUX4
THRB
ESRRG

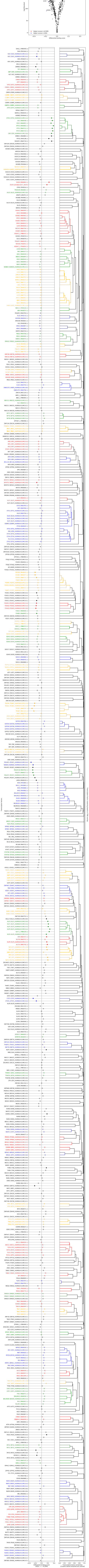
SIX6_1 (PB0059.1)
SIX4 (PH0164.1)
SIX2 (SIX2_HUMAN.H11MO.0.A)
SIX1 (SIX1_HUMAN.H11MO.0.A)
HDX (PH0037.1)
GFI1 (GFI1_HUMAN.H11MO.0.C)
NFIL3 (MA0025.1)
GMEB2 (MA0862.1)
TEF (MA0843.1)
DBP (DBP_HUMAN.H11MO.0.B)
HLF (HLF_HUMAN.H11MO.0.C)
GMEB1 (MA0615.1)
CREB5 (MA0840.1)
ATF7 (MA0834.1)
ATF2 (ATF2_HUMAN.H11MO.0.B)
CREB1 (CREB1_HUMAN.H11MO.0.A)
ATF1 (ATF1_HUMAN.H11MO.0.B)
CREM (CREM_HUMAN.H11MO.0.C)
CEBPB (CEBPB_HUMAN.H11MO.0.A)
CEBPA (CEBPA_HUMAN.H11MO.0.A)
CEBPD (CEBPD_HUMAN.H11MO.0.C)
CEBPE (CEBPE_HUMAN.H11MO.0.A)
CEBPG (CEBPG_HUMAN.H11MO.0.B)
ATF4 (ATF4_HUMAN.H11MO.0.A)

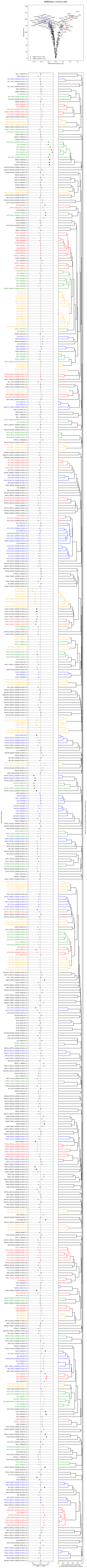

BINDetect volcano plot
-log10(pvalue)
Differential binding score
Higher scores in 8C
Higher scores in ICM

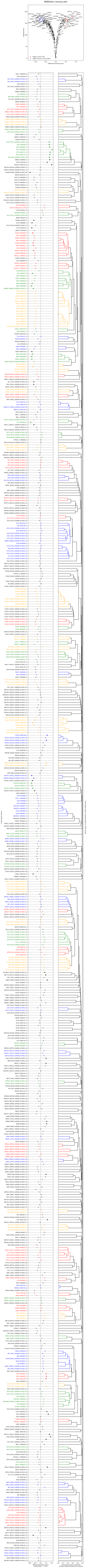

BINDetect volcano plot
-log10(pvalue)
Differential binding score
Higher scores in 8C
Higher scores in naive_hESC

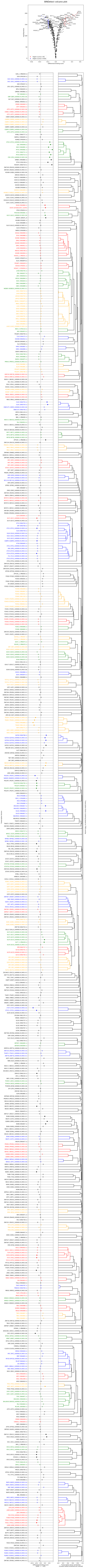
BINDetect volcano plot
-log10(pvalue)
Differential binding score
Higher scores in 8C
Higher scores in hESC

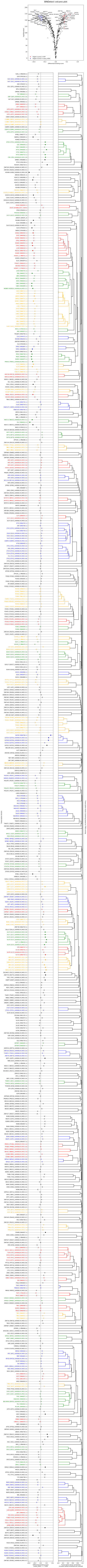

BINDetect volcano plot
-log10(pvalue)
Differential binding score
Higher scores in ICM
Higher scores in naive_hESC

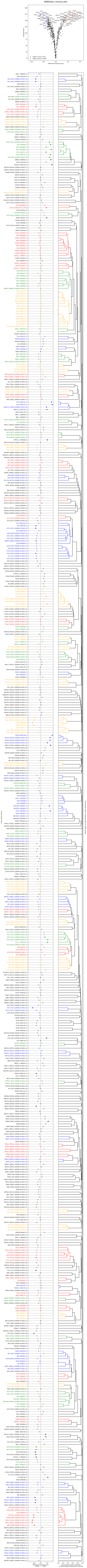
BINDetect volcano plot
-log10(pvalue)
Differential binding score
Higher scores in ICM
Higher scores in hESC
FOSL2
FOSL1
ZIC3
CTCF
TEAD2
TEAD1
TEAD3
FOS
ETV4
FOSB
TEAD4
LYL1
JUNB
NFE2
TCF3
NFE2L2
BACH2
SIX2
ETV1
SIX1
RFX1
RFX3
RFX4
ERF
MECP2
ATF7
KLF4
GATA2
RHOXF1
KLF7_1
GATA6
GATA5
GATA3
PITX1
OTX2
GSC
ZNF121
FOXK2
GATA4
DMBX1
KLF12
GLI3
CRX
PITX3
KLF5
PAX5
FOXG1
CUX1
ESRRG
CUX2
DUX4
ZFP187_1
OLIG1
ONECUT2
ARGFX
BHLHE23
GSC2
YY2
ONECUT1
OTX1

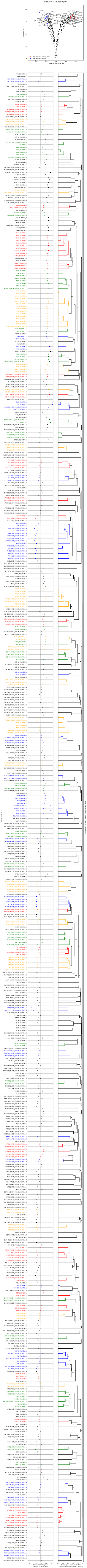
BINDetect volcano plot
-log10(pvalue)
Differential binding score
Higher scores in naive_hESC
Higher scores in hESC
