## Supplemental File 3 for "Beyond accessibility: ATAC-seq footprinting unravels kinetics of transcription factor binding during zygotic genome activation"

### BINDETECT FIGURES

Page 2) Raw score distributions
Page 3) Normalized score distributions
Page 4) BINDetect plot (early2C / 2C)
Page 5) BINDetect plot (early2C / 4C)
Page 6) BINDetect plot (early2C / 8C)
Page 7) BINDetect plot (early2C / ICM)
Page 8) BINDetect plot (early2C / mESC)
Page 9) BINDetect plot (2C / 4C)
Page 10) BINDetect plot (2C / 8C)
Page 11) BINDetect plot (2C / ICM)
Page 12) BINDetect plot (2C / mESC)
Page 13) BINDetect plot (4C / 8C)
Page 14) BINDetect plot (4C / ICM)
Page 15) BINDetect plot (4C / mESC)
Page 16) BINDetect plot (8C / ICM)
Page 17) BINDetect plot (8C / mESC)
Page 18) BINDetect plot (ICM / mESC)

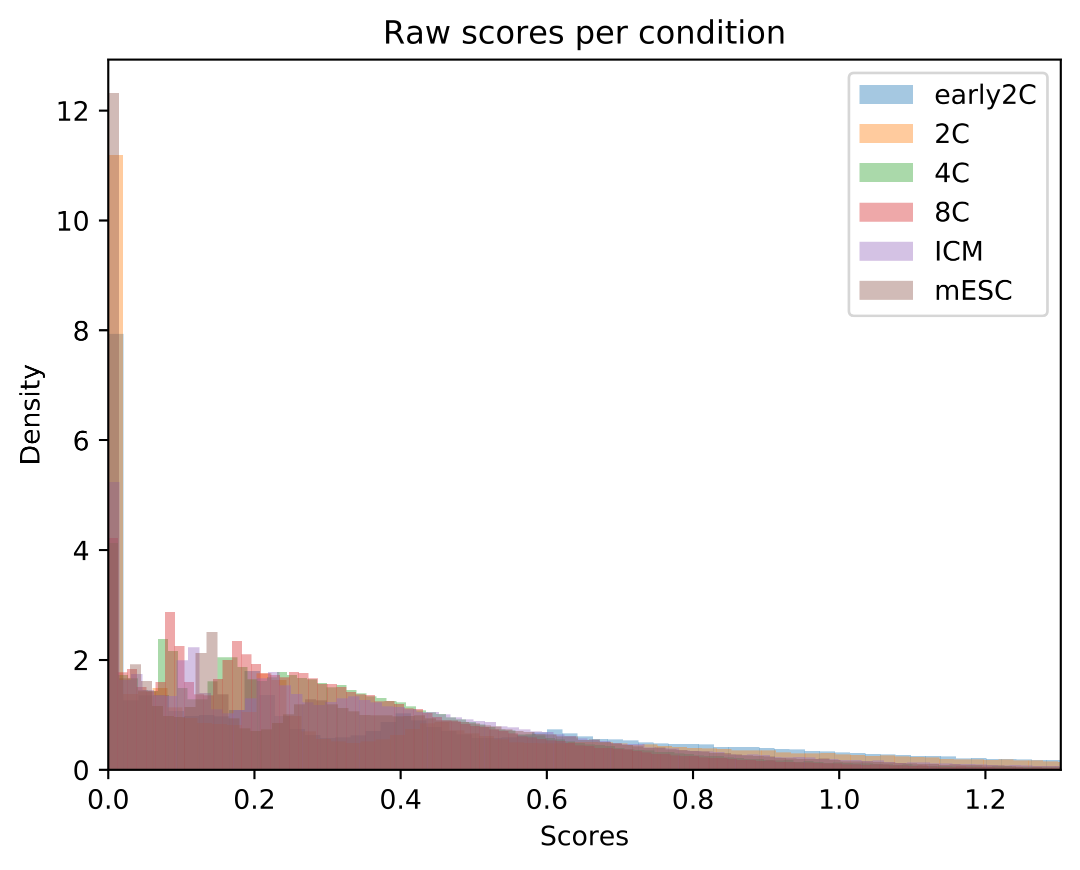

Raw scores per condition
early2C
2C
4C
8C
ICM
mESC
Density
Scores
0
2
4
6
8
10
12
0.0
0.2
0.4
0.6
0.8
1.0
1.2

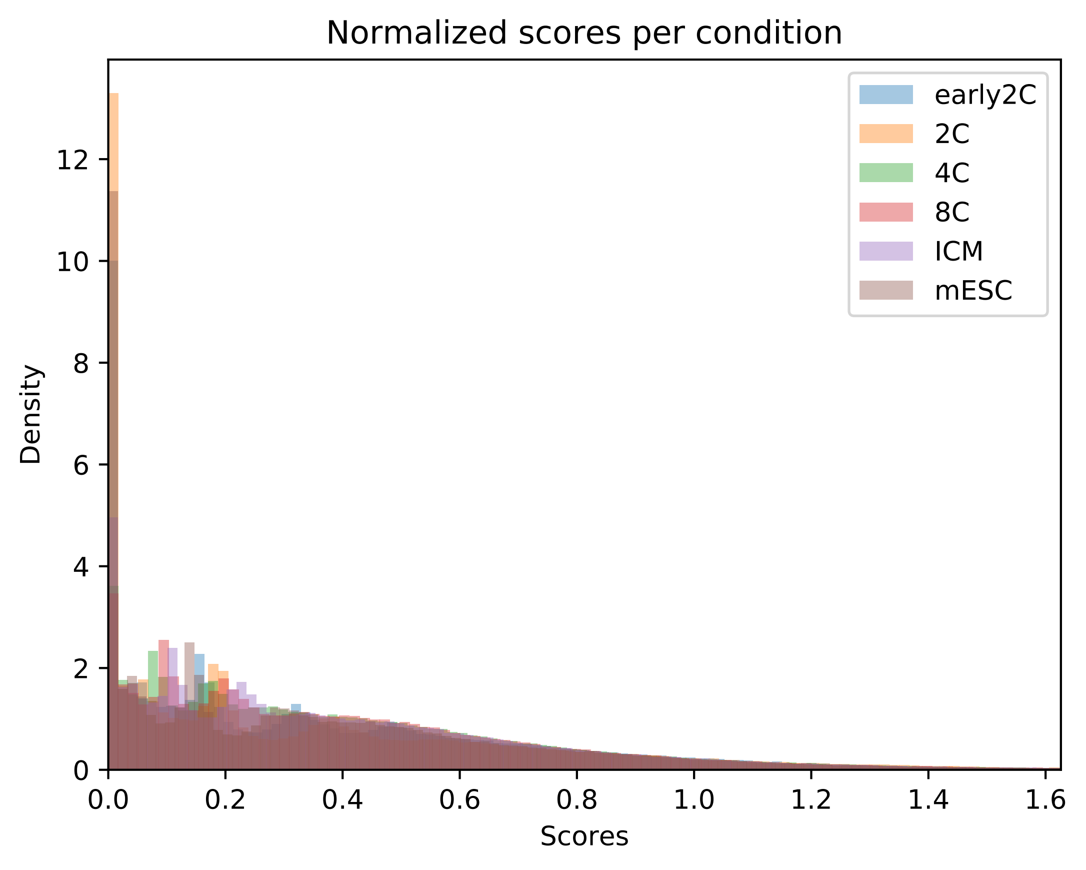
Normalized scores per condition
Density
Scores
early2C
2C
4C
8C
ICM
mESC
0
2
4
6
8
10
12
0.0
0.2
0.4
0.6
0.8
1.0
1.2
1.4
1.6

BINDetect volcano plot

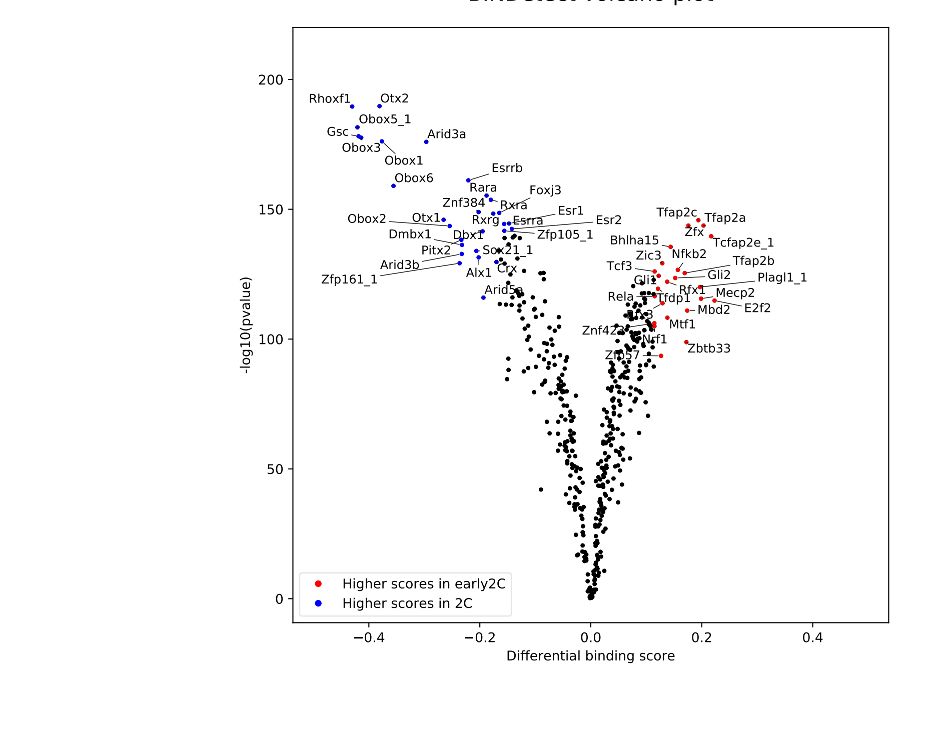

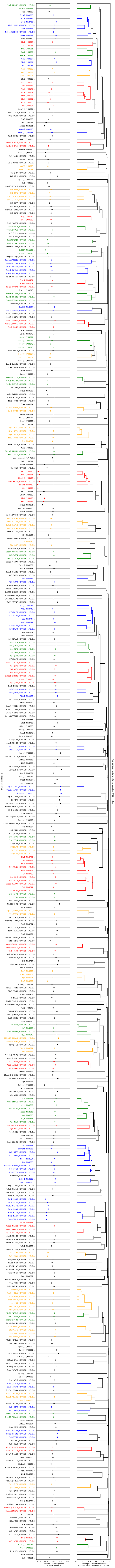

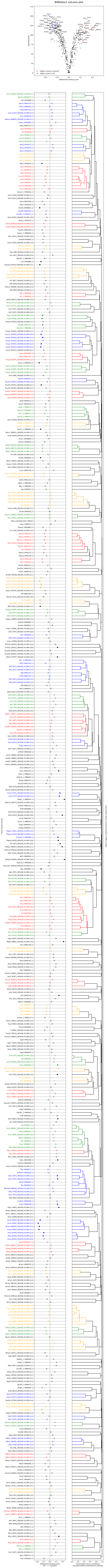
BINDetect volcano plot

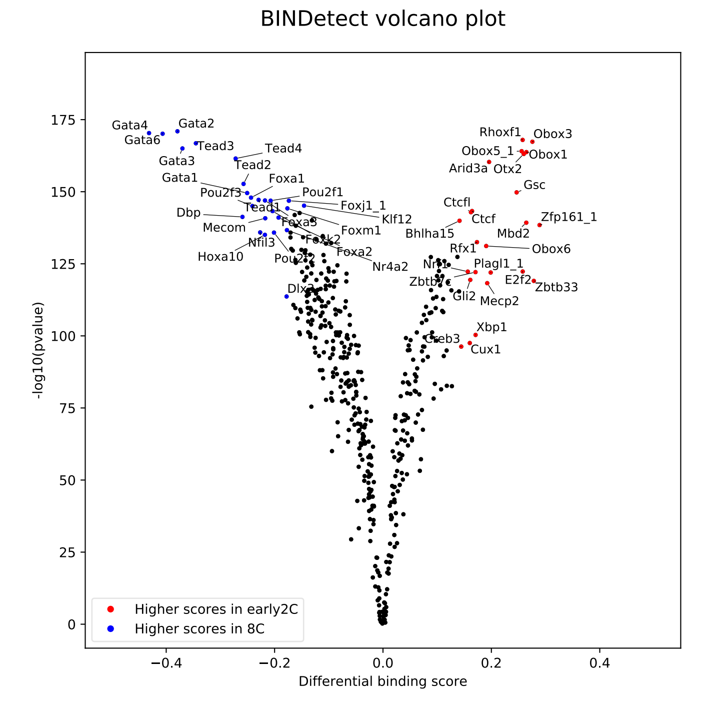

BINDetect volcano plot
Gata4
Gata2
Gata6
Tead3
Tead4
Gata3
Tead2
Gata1
Foxa1
Pou2f3
Pou2f1
Tead1
Foxj1_1
Dbp
Klf12
Foxa3
Mecom
Foxm1
Foxk2
Nfil3
Foxa2
Hoxa10
Pou2f2
Nr4a2
Dlx2
Rhoxf1
Obox3
Obox5_1
Obox1
Arid3a
Otx2
Gsc
Ctcfl
Ctcf
Zfp161_1
Bhlha15
Mbd2
Rfx1
Obox6
Nr1
Plagl1_1
E2f2
Zbtb33
Gli2
Mecp2
Xbp1
Creb3
Cux1
Higher scores in early2C
Higher scores in 8C
-log10(pvalue)
Differential binding score


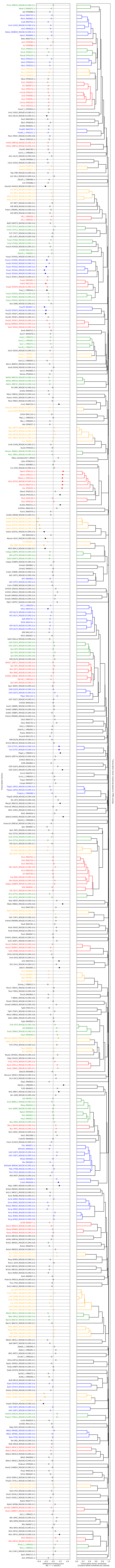

Prrx2 (PRRX2_MOUSE.H11MO.0.C)
Nkx6-2 (MA0675.1)
Isl2 (PH0088.1)
Shox2 (MA0720.1)
Mixl1 (MA0662.1)
Lhx8 (MA0705.1)
Lhx2 (LHX2_MOUSE.H11MO.0.A)
Lbx1 (MA0618.1)
Nobox (NOBOX_MOUSE.H11MO.0.C)
Hesx1 (MA0894.1)
Noto (MA0710.1)
Lbx2 (PH0090.1)
Isx (PH0089.1)
Evx1 (PH0031.1)
Emx2 (PH0027.1)
Rhox6 (PH0159.1)
Msx2 (PH0107.1)
Gbx2 (PH0034.1)
Gbx1 (PH0033.1)

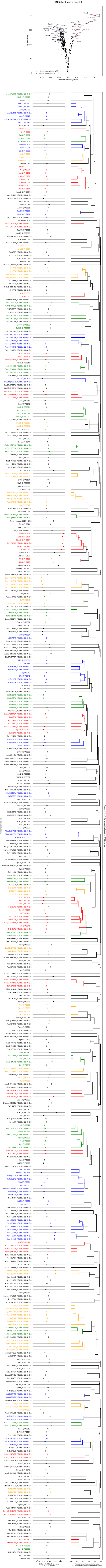
BINDetect volcano plot

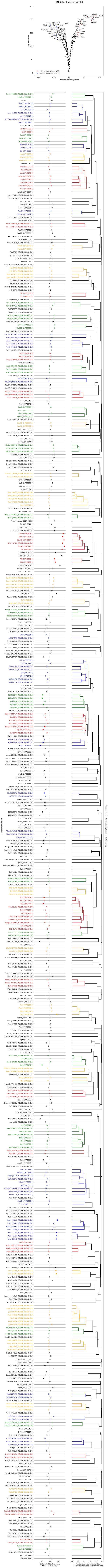
BINDetect volcano plot

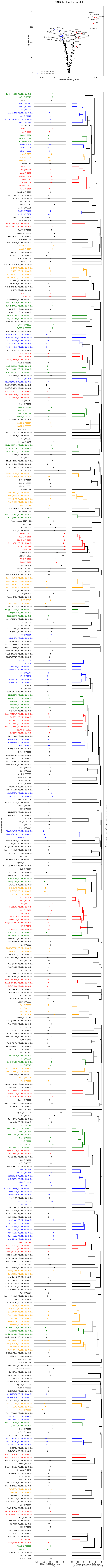
BINDetect volcano plot

BINDetect volcano plot

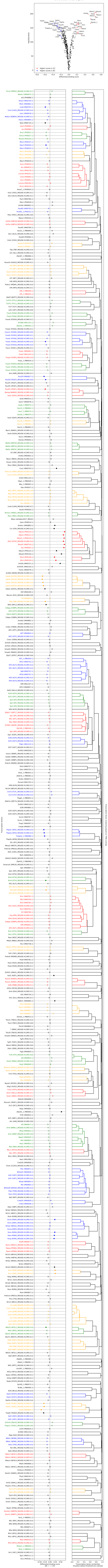

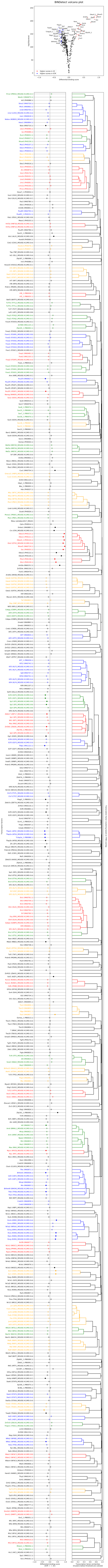
BINDetect volcano plot

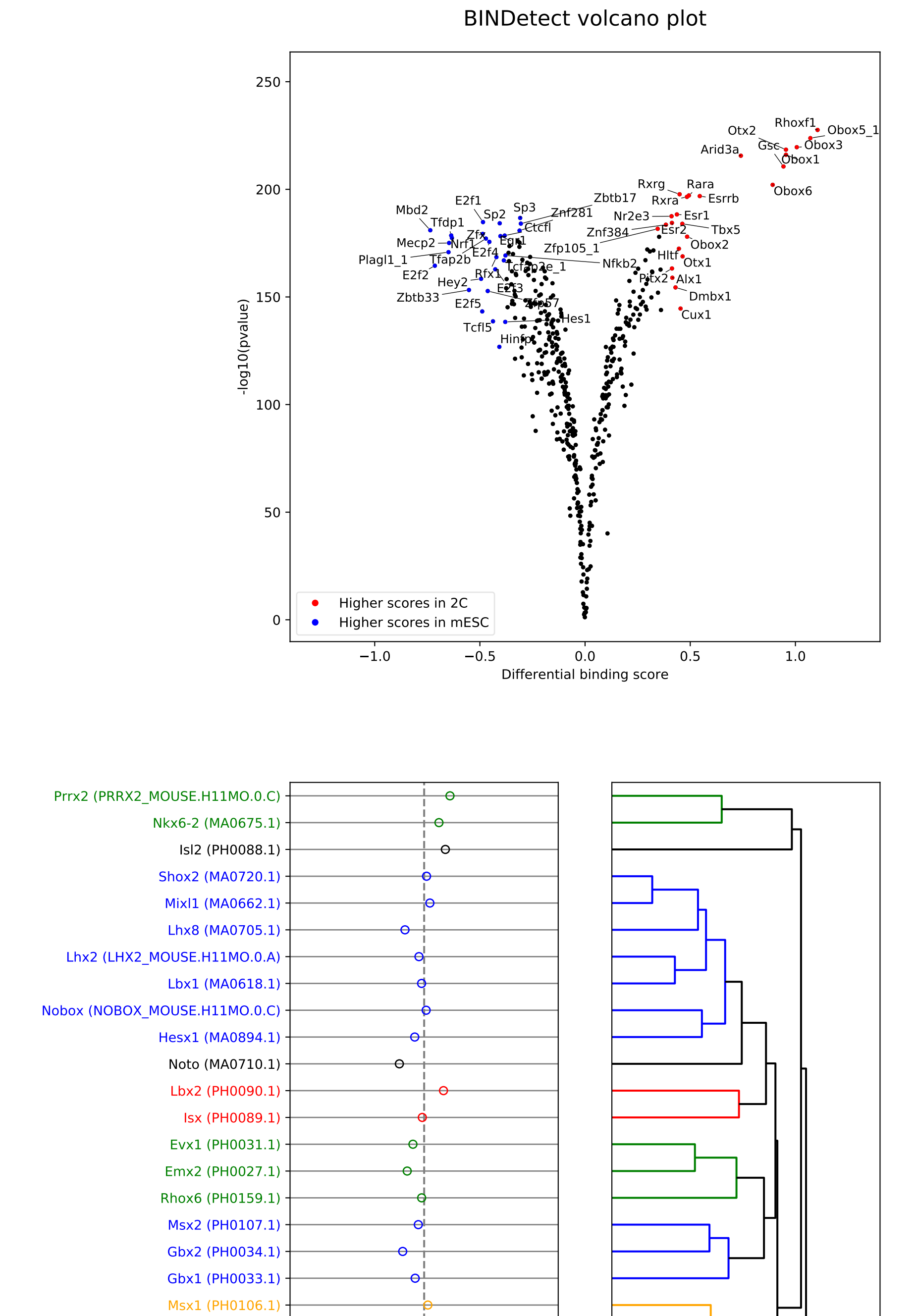
BINDetect volcano plot

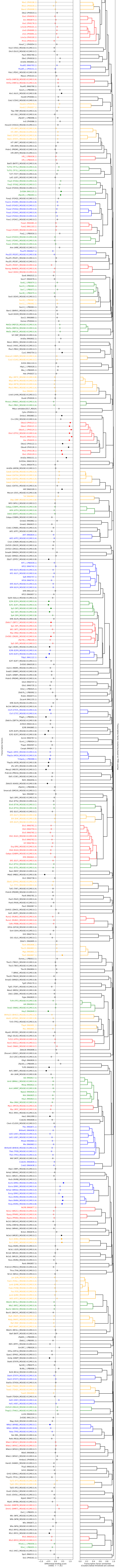

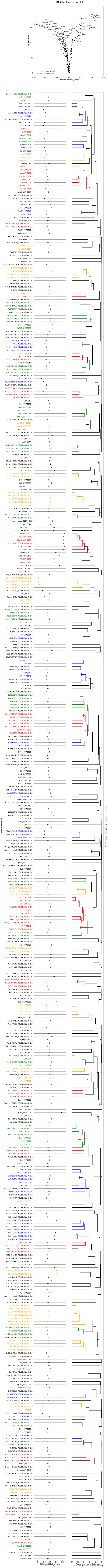
BINDetect volcano plot

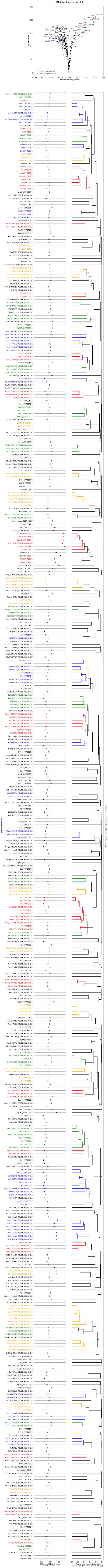
BINDetect volcano plot

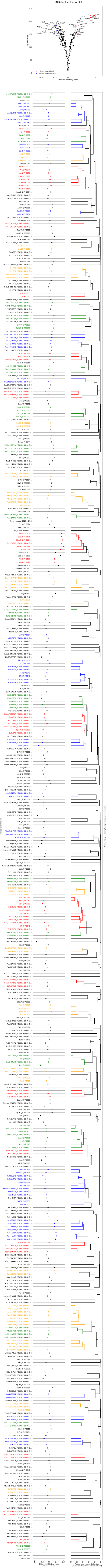
BINDetect volcano plot

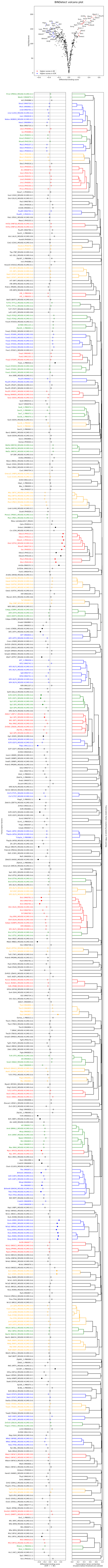
BINDetect volcano plot

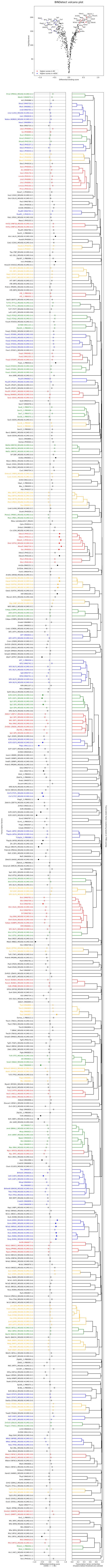
BINDetect volcano plot

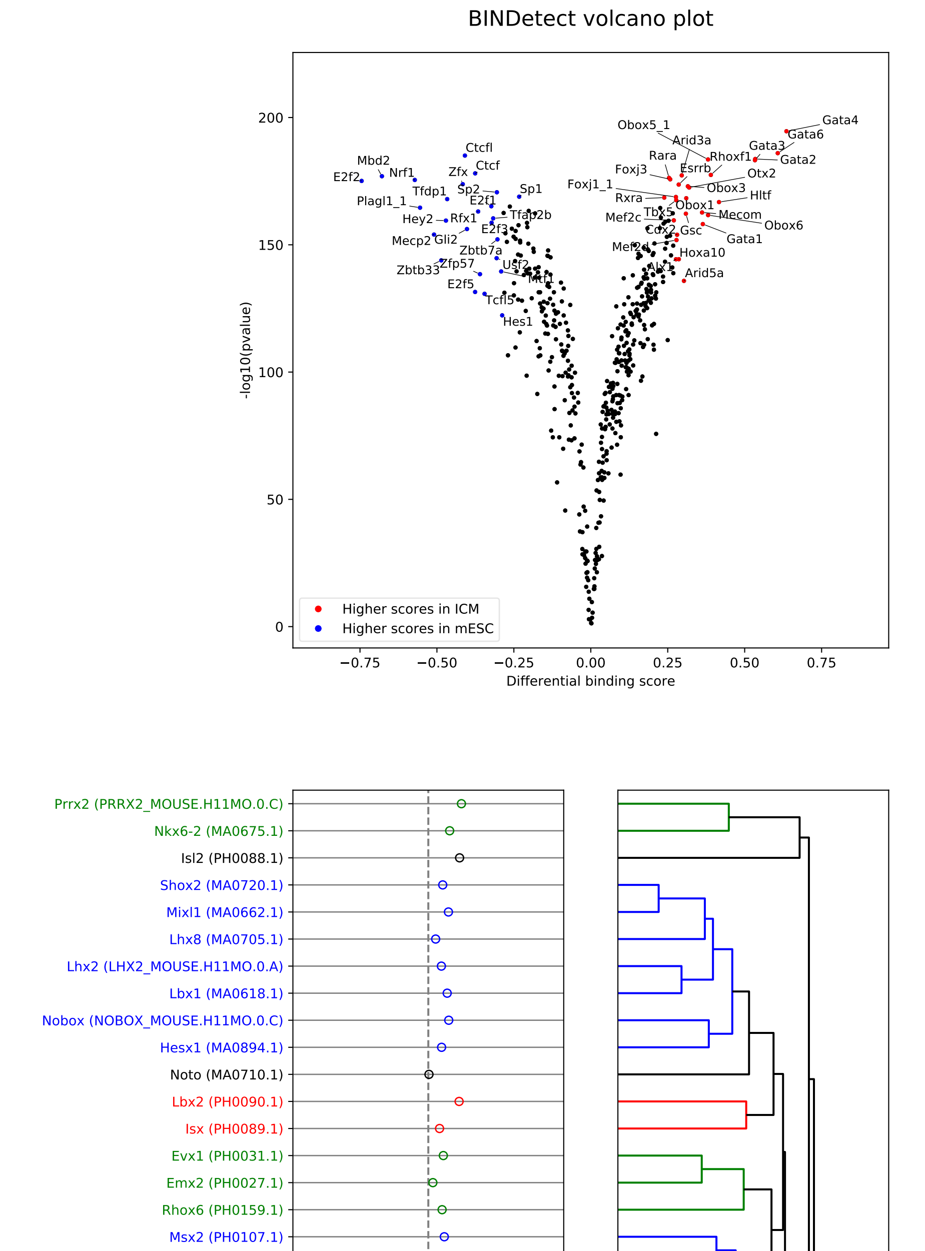
BINDetect volcano plot

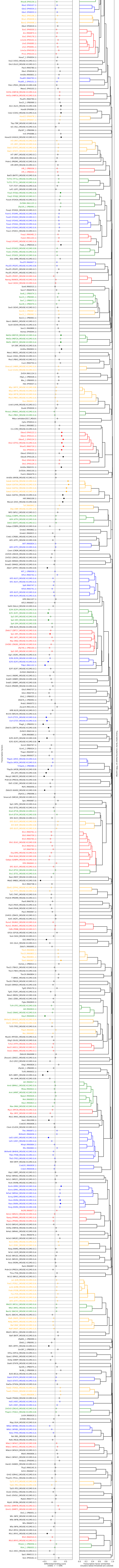
