## Supplemental File 4 for "Beyond accessibility: ATAC-seq footprinting unravels kinetics of transcription factor binding during zygotic genome activation"

### BINDETECT FIGURES

Page 2) Raw score distributions
Page 3) Normalized score distributions
Page 4) BINDetect plot (mDuxNeg / mDuxPos)

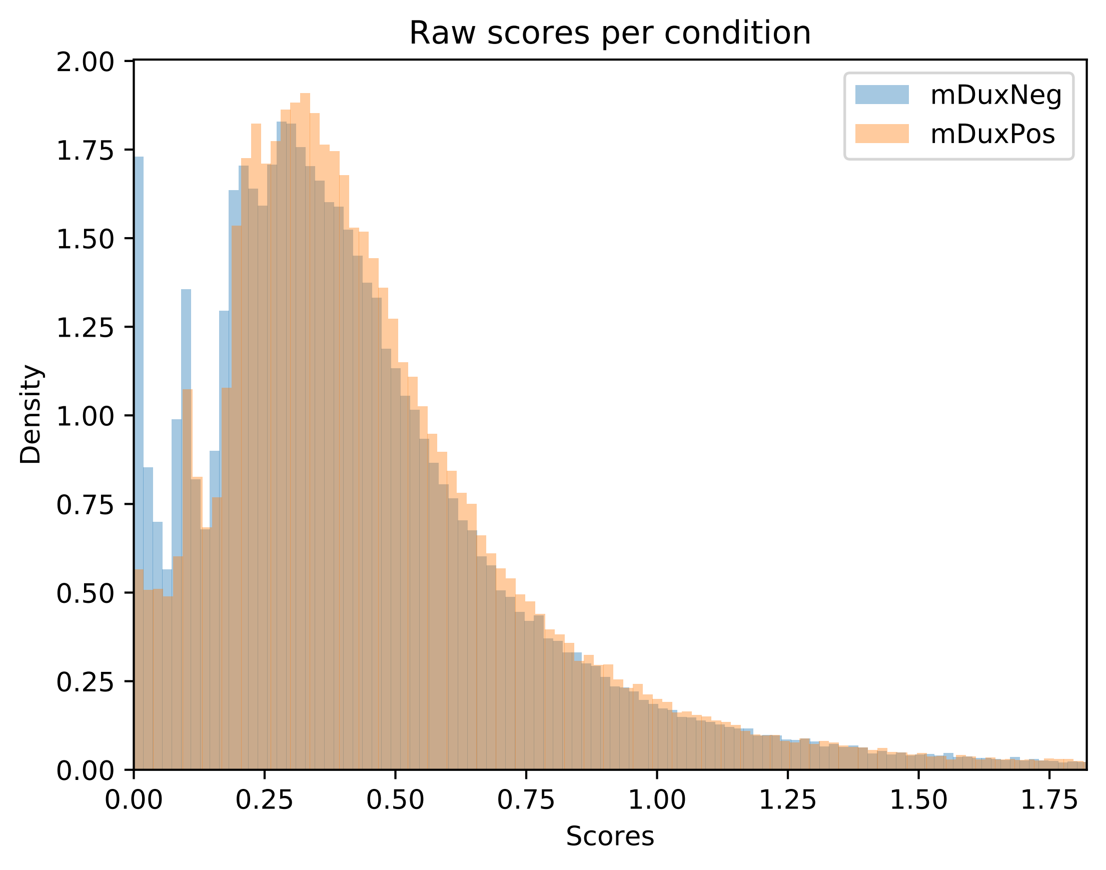

Raw scores per condition
mDuxNeg
mDuxPos
Density
Scores
2.00
1.75
1.50
1.25
1.00
0.75
0.50
0.25
0.00
0.00
0.25
0.50
0.75
1.00
1.25
1.50
1.75

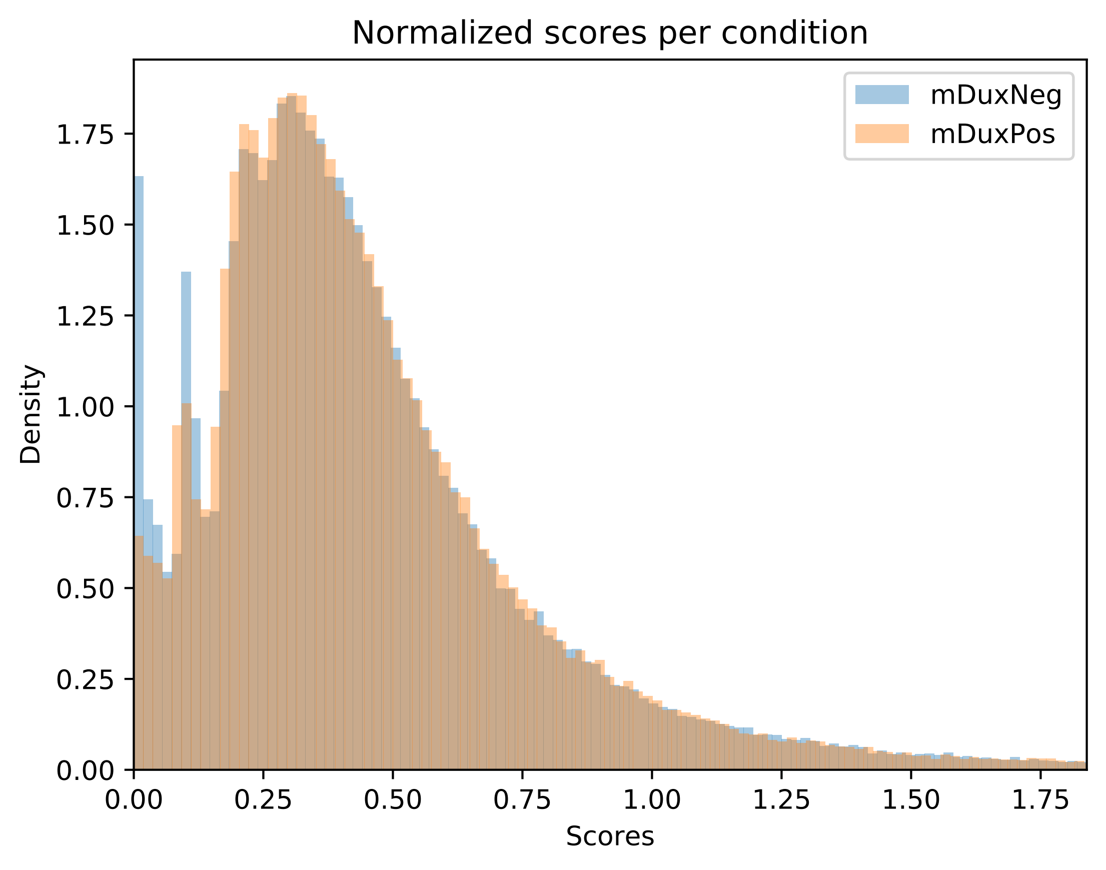
Normalized scores per condition
mDuxNeg
mDuxPos
Density
Scores
0.00
0.25
0.50
0.75
1.00
1.25
1.50
1.75

BINDetect volcano plot

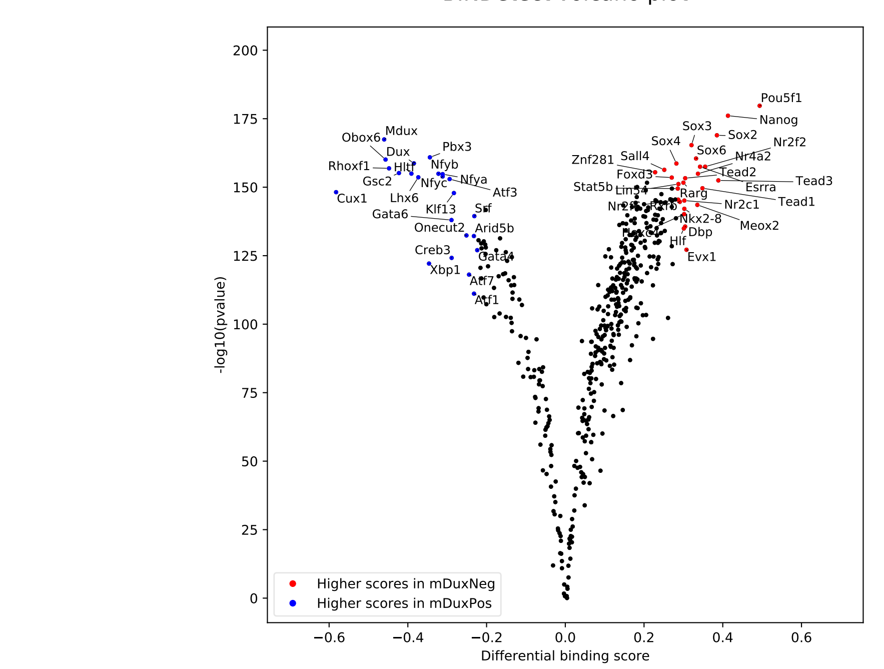

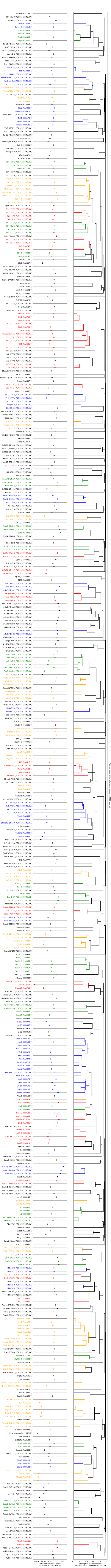
